# SPARK: Physiology-informed inference of latent cardiac autonomic dynamics from RR intervals

**DOI:** 10.64898/2026.11.06.764976

**Authors:** Matías Castillo-Aguilar, Noah Beelders, Cristian Núñez-Espinosa

## Abstract

Cardiac autonomic activity is observed only through beat-to-beat (RR) intervals, and separating it into parasym-pathetic-like and sympathetic-like components is an inverse problem that RR alone is not expected to solve uniquely. We introduce SPARK (Stochastic Phase-domain Autonomic Reconstruction Kinetics), a state-space model in which two independent, mean-reverting latent drives modulate heart rate through an integrate-and-fire pacemaker. The model is combined with exact continuous-time Kalman filtering and with an explicit physiological identification structure: time-scale priors, a kinetic ordering and a spectral partition of heart rate variability. SPARK offers a model-conditional description of autonomic trajectories that complements conventional heart rate variability; its latent states are reconstructions, not measurements of neural activity. Across recordings, fitting each recording from many different starting points gave a median of 23.5 distinct solutions without physiological information versus 1.0 with it, and one process absorbed essentially all the variability in 75% versus 4% of the fits; the fit diagnostics showed residual structure in most recordings. Reconstructions were then examined: in 10 tilt-table recordings, the vagal-like state decreased and the sympathetic-like state increased in 10 of 10; exploratory comparisons between 20 young and 20 older adults showed differences in reconstructed variability and faster-component kinetics, although several latent summaries were mathematically redundant; and in 263 exercise recordings, median recovery times, which are closely tied to the fitted time constants, were 42 s (vagal-like) and 341 s (sympathetic-like). How a stressor-induced change is split between the two drives remained sensitive to the spectral anchor.

**Highlights:**

- RR alone may leave vagal-like and sympathetic-like drives ambiguous
- A phase-domain state-space model filters irregular RR intervals in continuous time
- Physiological priors and a spectral anchor restrict the set of latent solutions
- Latent states are examined during tilt, across aging and in exercise recovery
- How change is split between the two drives remains conditional on the anchor

## 1 Introduction

Cardiac autonomic regulation operates over multiple time scales. The sinoatrial node, the primary chronotropic interface, translates continuous neural efferent traffic into a sequence of discrete cardiac contractions [1,2]. Parasympathetic signaling, mediated by acetylcholine, acts rapidly to slow pacemaker firing, whereas sympathetic signaling, mediated by norepinephrine, acts more slowly to accelerate it [3,4]. Physiological challenges such as postural change or exercise therefore elicit time-resolved autonomic responses, including withdrawal of vagal influence, delayed sympathetic engagement and subsequent recovery, whose time courses are of physiological interest in their own right [5].

The inter-beat (RR) interval series is the most accessible observable of this regulation, but it reflects the integrated influence of several physiological processes rather than the activity of either autonomic branch itself [3]. Direct measurement of autonomic nerve activity by microneurography is invasive and technically constrained, and the growing use of wearable and photoplethysmographic acquisition further motivates inference from interval data alone [6]. Recovering the latent autonomic dynamics from RR is therefore a problem of inference from an indirect observable [5,7].

Heart rate variability (HRV) analysis has long provided the standard non-invasive description of RR variability, summarizing the series with time-domain statistics and frequency-domain spectral measures [5,8–10]. These descriptors are widely used and informative, but they quantify the output of the system rather than offering a generative representation of its latent dynamics. Conventional spectral partitioning assumes stationarity, which is not satisfied during transient perturbations [5,11], and the low-frequency band reflects a mixture of sympathetic, vagal and baroreflex influences [8,12]. Descriptive measures also do not model how interval timing is generated by the sinoatrial pacemaker [5].

Generative alternatives model the sinoatrial node and its autonomic inputs, from multiscale biophysical simulations [3,13] to integrate-and-fire oscillators in which a phase accumulates at a rate set by autonomic inputs and a beat occurs when the phase reaches a threshold [14]. For a classical drift–diffusion phase, that is, one whose diffusive component is a Brownian motion, interval durations follow an inverse Gaussian distribution [15,16]. Embedding this mechanism in a state-space model, however, requires solving a nonlinear first-passage problem, which is usually handled by local linearization, sigma-point filtering or Euler–Maruyama discretization [7,17,18]. Such approximations can distort the continuous-time dynamics across irregular intervals [7]. Deep-learning models, although effective for pattern recognition, typically do not yield latent variables with explicit physiological meaning.

A second and conceptually distinct difficulty concerns identification. If RR is regarded as the outcome of a parasympathetic-like process, a sympathetic-like process and observation noise, recovering the two processes from a single interval series is an inverse problem [19]. Different combinations of time scales, amplitudes and noise can generate similar interval series, so the observations alone are not guaranteed to determine a unique decomposition. Under these conditions, unconstrained maximum-likelihood estimation can produce likelihood ridges, scale ambiguities and blending of the two processes; we examine whether this occurs in the present recordings (Section 3.1).

A good fit to RR is therefore necessary but not sufficient evidence of a meaningful decomposition: a solution may reproduce the observed intervals closely and still be physiologically implausible, for example by attributing all variability to one process or by assigning implausible time scales. We consider independent physiological information not as a device to improve fit, but as part of the identification structure that defines which latent solutions are admissible. Two sources of independent physiological information are used here: the characteristic time scales of vagal and sympathetic effector responses together with the ordering of their kinetics [2,20], and spectral evidence, including seminal pharmacological-blockade studies, on how modulation by the two branches is distributed across frequency [21–23]. These assumptions are explicit and external to the RR data, and how far they shape the inferred trajectories should be examined, not assumed.

We implement this strategy in SPARK (Stochastic Phase-domain Autonomic Reconstruction Kinetics). In the language of nonlinear stochastic dynamics, the model is a pair of mean-reverting processes with well-separated time scales that drive a threshold-crossing phase oscillator with resetting, and the question is how much of this hidden structure can be recovered from the crossing times alone. SPARK comprises three parts: the generative model, the identification structure that restricts its admissible solutions, and the inference that connects both to the data. For inference we use the Kalman filter, which is exact for this model because the measurement equation is conditionally linear and Gaussian (see below); other filters, such as sigma-point or particle filters, could be applied to the same model, but they would be approximate and are not needed for this formulation.

The individual components are established: mean-reverting Ornstein–Uhlenbeck processes [24], integrate-and-fire pacing [14], Kalman filtering [25] and spectral partitions of heart rate variability [21]. What this study adds is their combination, in three parts. First, a phase-domain (terminal-threshold) conditioning: each recorded RR interval is treated as a deterministic observation window, and the model asks what accumulated phase is implied at its end. This replaces the strict first-passage condition with a terminal-threshold condition that approximates it closely in the high-drift, low-noise regime of the sinoatrial node [26] and makes the measurement equation Gaussian and linear in the time-integrals of the latent drives [27]. Because the drives are Ornstein–Uhlenbeck processes, their transitions across irregular intervals are computed analytically. Second, an explicit identification structure, in which the time-scale priors, the kinetic ordering and a wavelet-derived spectral anchor define the admissible latent solutions. Third, a systematic assessment, in real recordings, of how far the decomposition is determined by the RR data and how far by those assumptions, through identifiability and sensitivity analyses.

To our knowledge, this combination of phase-domain conditioning, exact continuous-time propagation and an explicit physiological identification structure has not been applied to RR-interval inference before; we claim no novelty for the individual components, and the resulting states have not been validated against nerve activity.

We evaluated the framework using the following sequence of questions. Does RR alone leave the latent decomposition under-determined, and does physiological information reduce this ambiguity? Do the latent states show the expected direction of change during an orthostatic perturbation? How do they behave across healthy aging? Do they characterize adaptation and recovery in older adults during a rest–exercise–rest protocol? Conventional HRV descriptors are used as comparators rather than competitors, and independent measurements available in subsets of the recordings (tilt angle, arterial pressure and respiration) are external comparators and not inputs to the model. We emphasize at the outset that the latent states are reconstructions conditional on the stated assumptions: the aim is a model-based description of autonomic dynamics, not a measurement of neural activity.

## 2 Methods

This section describes the model, the assumptions that make its latent decomposition identifiable, inference and diagnostics, and the three datasets used for the perturbation, aging and exercise analyses. Derivations are in the supplementary material. The numerical settings of the optimizer and of the analyses are the defaults of the accompanying R engine, and the configuration actually used in every analysis (for instance, the number of starting points and the analysis windows) is stored with the results.

### 2.1 Conceptual model

SPARK represents the sinoatrial node as an integrate-and-fire pacemaker whose net drift is modulated by two latent autonomic drives. Let ϕ(*t*) denote the sinoatrial phase. Between two consecutive beats at times *t*_*k*−1_ and *t*_*k*_, starting from ϕ(*t*_*k*−1_) = 0, the phase evolves as a drift–diffusion process,

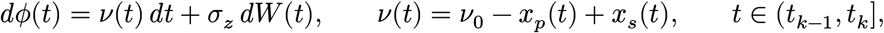

where *W* (*t*) is a standard Wiener process (Brownian motion), and a beat occurs the first time ϕ reaches a normalized threshold of 1 [14], after which ϕ is set to zero and the process restarts. The net drift *v*(*t*) is the instantaneous pacing rate (Hz), *v*_0_ is the baseline pacing rate (the mean rate when both drives are at their baseline), *σ*_*z*_ is the diffusion coefficient of threshold-level noise, and *x*_*p*_(*t*) and *x*_*s*_(*t*) are latent parasympathetic-like and sympathetic-like drives, respectively, expressed in Hz as deviations from their homeostatic baseline. Hereafter we call them the vagal and sympathetic drives for brevity. These labels denote model components distinguished by their kinetics and spectral allocation (see *Model assumptions and identification structure*), not measured neural activity.

### 2.2 Latent autonomic dynamics

The drives are independent, mean-reverting Ornstein–Uhlenbeck (OU) processes [24], which represent the tendency of each branch to return to its homeostatic level after a transient stimulus:

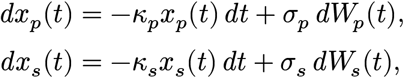

where *k*_*p*_ and *k*_*s*_ are clearance rates (s^−1^) with e-folding times *τ*_*p*_ = 1/*k*_*p*_ and *τ*_*s*_ = 1/*k*_*s*_. To keep amplitude independent of kinetics, the diffusion coefficients are parameterized through the stationary variances, 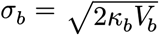, with 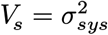 and 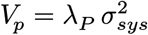; *λ*_*P*_ = *V*_*p*_/*V*_*s*_ is the relative vagal-to-sympathetic variance. The global scale *σ*^2^ multiplies all the covariances of the model but leaves the Kalman gain unchanged, so it can be estimated analytically instead of by numerical optimization (see *Inference and uncertainty*). To make the observation equation linear, the state is augmented with the running integrals of the drives, *dI*_*p*_ = *x*_*p*_ *dt* and *dI*_*s*_ = *x*_*s*_ *dt*, giving 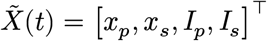 and 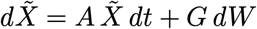 with constant matrices *A* and *G*. The transition over any observed interval Δ*t*_*k*_ is then given by the matrix exponential, Φ_*k*_ = exp(*A* Δ*t*_*k*_), with a closed-form process covariance (see the “Matrix exponential” and “Estimation algorithm” sections of the supplementary material, which also list the model quantities and their estimation status).

### 2.3 Observation model: phase-domain inversion

In the strict first-passage formulation, the observed interval is the random time *T* = inf{*t* > 0 : ϕ(*t*) ≥ 1}, whose distribution is inverse Gaussian for a constant drift [15]; estimating a time-varying drift from it within a state-space model is a nonlinear problem [7,17]. We instead condition on the observed interval: the recorded duration Δ*t*_*k*_ is a fixed window, and under a terminal-threshold condition the phase accumulated over [*t*_*k*−1_, *t*_*k*_] equals one,

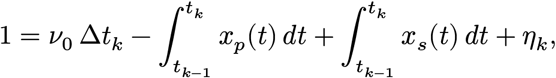

where *η*_*k*_ collects the Gaussian threshold-level noise, with variance 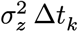 and 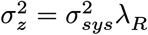. In the strict formulation the beat occurs the first time ϕ touches the threshold, so ϕ must stay below 1 during the whole interval. The terminal-threshold condition uses only the value of the phase at *t*_*k*_ and drops this path-survival requirement; it therefore neglects the paths that reach the threshold earlier within the interval and then fall back below it. This is accurate when the drift is large relative to the noise, because such paths are then rare (an explicit bound is given in the “Asymptotic convergence” section of the supplementary material). In the augmented state the observation is linear,

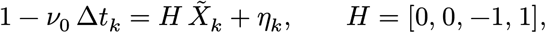

so standard Kalman recursions apply. After each measurement update the integral states and their covariances are reset to zero, whereas *x*_*p*_ and *x*_*s*_ continue to evolve, which represents the restart of phase accumulation at each beat while preserving the trans-beat memory of the drives. To limit the influence of ectopic beats and detection artifacts, the threshold-noise variance at beat *k* is inflated by a multiplier that grows with the largest relative change between adjacent intervals, which removes abrupt jumps from the likelihood without interrupting the propagation of the drives. The multiplier uses both the preceding and the following interval and is therefore not strictly causal; it affects only the weighting of the observation. It is a heuristic robustness device and not a model of arrhythmia, so recordings with substantial non-sinus activity are outside the intended scope of the framework.

### 2.4 Model assumptions and identification structure

The decomposition of RR into two latent drives is not guaranteed to be unique, and we therefore make explicit the physiological information that restricts the admissible solutions (Table 1). These specifications are model assumptions informed by external evidence; they are not estimated from the present data, and their influence on the inferred trajectories is evaluated in the identifiability analyses described below. Each is justified in turn after the table.

**Table 1:**
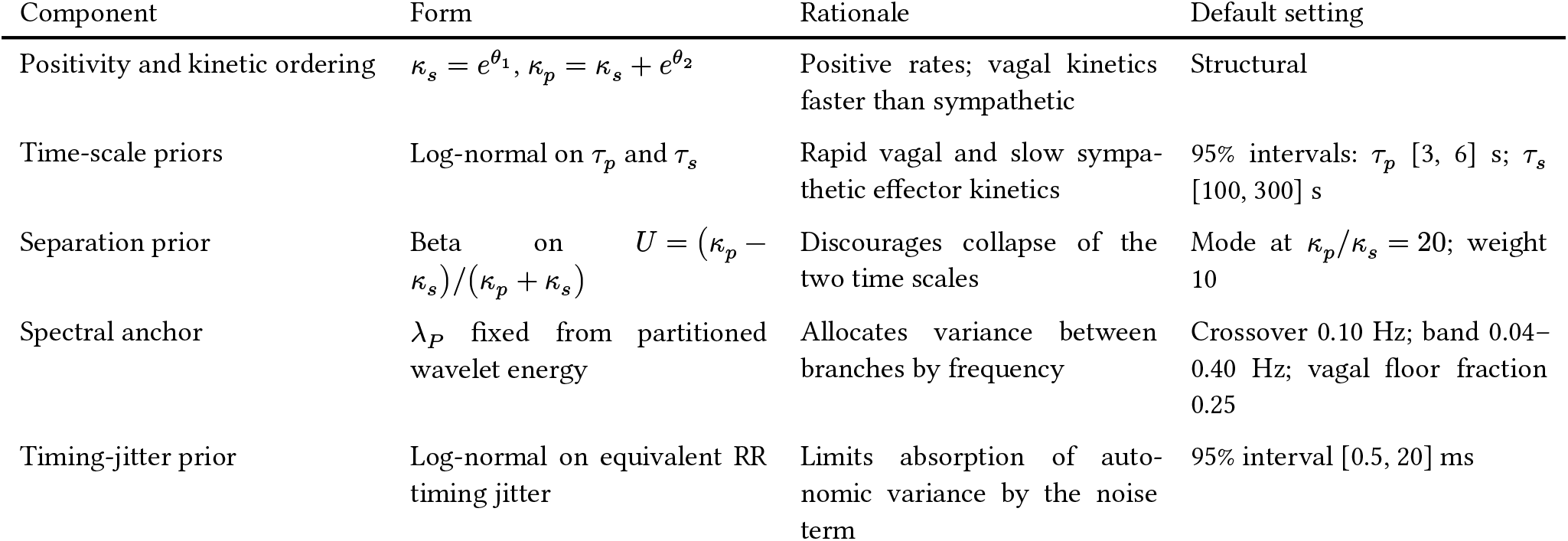
Components of the identification structure.

| Component | Form | Rationale | Default setting |
|---|---|---|---|
| Positivity and kinetic ordering | $\kappa_s = e^{\theta_1}$, $\kappa_p = \kappa_s + e^{\theta_2}$ | Positive rates; vagal kinetics faster than sympathetic | Structural |
| Time-scale priors | Log-normal on τ<sub>p</sub> and τ<sub>s</sub> | Rapid vagal and slow sympathetic effector kinetics | 95% intervals: τ<sub>p</sub> [3, 6] s; τ<sub>s</sub> [100, 300] s |
| Separation prior | Beta on $U = (\kappa_p - \kappa_s)/(\kappa_p + \kappa_s)$ | Discourages collapse of the two time scales | Mode at κ<sub>p</sub>/κ<sub>s</sub> = 20; weight 10 |
| Spectral anchor | λ<sub>P</sub> fixed from partitioned wavelet energy | Allocates variance between branches by frequency | Crossover 0.10 Hz; band 0.04–0.40 Hz; vagal floor fraction 0.25 |
| Timing-jitter prior | Log-normal on equivalent RR timing jitter | Limits absorption of autonomic variance by the noise term | 95% interval [0.5, 20] ms |

#### Structural constraints

Exponential links ensure positive rates, and *k*_*p*_ = *k*_*s*_ + exp(*θ*_2_) imposes *k*_*p*_ > *k*_*s*_, which removes label switching, that is, the interchangeability of the two drives (a solution in which the fast and the slow process exchange roles would otherwise fit the intervals equally well).

#### Time-scale information

Independent log-normal priors on *τ*_*p*_ and *τ*_*s*_, and a Beta prior on *U* that discourages the two time scales from collapsing onto one another, reflect the expected separation between rapid vagal and slow sympathetic effector kinetics [2,20]. The limits for *τ*_*p*_ align with rapid parasympathetic signaling and respiratory cycles, and those for *τ*_*s*_ with the inertia and low-pass behavior of sympathetic modulation [20,22,28]. The priors are fixed across recordings and not derived from the individual series.

#### Spectral information

The ratio *λ*_*P*_ is a deterministic function of the global wavelet spectrum of the heart rate [29,30], which is partitioned into vagal and sympathetic energy with logistic frequency weights: sympathetic weight dominates at low frequencies, vagal weight rises at a crossover of 0.10 Hz, sympathetic weight falls to 5% at the boundary between the low-frequency (LF) and high-frequency (HF) bands at 0.15 Hz, and the vagal branch keeps a floor fraction of the low-frequency weight to reflect its contribution under non-stationarity. The weights encode external evidence on how modulation by the two branches is distributed across frequency, including pharmacological-blockade studies [21–23,31,32]. Because *λ*_*P*_ is computed from the series being fitted, it is data-informed and thus not independent of the data; the physiological information resides in the weighting function, and equating the ratio of spectral energies with the ratio of latent variances is an approximation (“Spectral anchoring” section of the supplementary material).

#### Timing jitter

The equivalent RR timing jitter is the standard deviation, in milliseconds, of the error in the timing of each beat that the threshold-level noise produces; it is the noise term that could otherwise absorb autonomic variability. A prior on it, with a 95% interval of 0.5–20 ms, spans the range expected for electrocar-diographic and photoplethysmographic acquisition.

### 2.5 Inference and uncertainty

For a given parameter vector, the Kalman recursion (with the Joseph form of the covariance update) processes the observed intervals sequentially. The baseline *v*_0_ enters the observation equation linearly and, together with the global scale 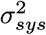, is profiled out analytically by generalized least squares, using two filters that share a single Kalman gain; the profiled log-likelihood is computed over the beats with an artifact multiplier below 100. The remaining parameters *θ* = (ln *k*_*s*_, ln(*k*_*p*_ − *k*_*s*_), ln *λ*_*R*_) are estimated by maximum a posteriori (MAP) estimation, that is, by minimizing the negative log-posterior (the negative profiled log-likelihood plus the negative log-prior terms of Table 1) with the Broyden–Fletcher–Goldfarb–Shanno (BFGS) quasi-Newton algorithm in coordinates scaled by the spread of the time-scale priors, started from three points (the centers of the priors, and two points with negligible and with dominant observation noise), keeping the run with the lowest objective.

States were estimated by forward filtering conditional on recording-level parameter estimates and the spectral anchor. Because parameter estimation and spectral anchoring use the complete recording, and artifact weighting uses neighboring intervals, the overall procedure is retrospective rather than strictly causal. The reported 95% intervals for the states and for the filtered RR are conditional on the MAP kinetic parameters and do not incorporate uncertainty in those parameters. Standard errors for *k*_*p*_, *k*_*s*_ and *λ*_*R*_ come from the numerical Hessian of the MAP objective and the delta method; profiled, fixed and derived quantities carry no standard error. Algorithmic details are given in the “Estimation algorithm” section of the supplementary material. The model was implemented in R, and a browser-based implementation with the same specification (default settings as in this study: *M*_3_, vagal floor 0.25) is available for single recordings (see Code availability).

### 2.6 Model diagnostics

#### Fit and convergence

Convergence is defined by the optimizer return code. Filtered tracking accuracy is quantified by the mean absolute percentage error (MAPE) and the root mean square error (RMSE) between the observed and filtered implied RR; these are in-sample measures, because the implied RR at each beat uses states updated with that beat, and they are not out-of-sample prediction error.

#### Innovation diagnostics

The standardized innovations 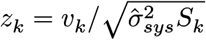, where *v*_*k*_ is the one-step prediction error of the observation and 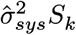 its variance, should be independent standard normal variates if the model is adequate. Under a correct model, *u*_*k*_ = Φ(*z*_*k*_), the probability integral transform of *z*_*k*_ by the standard normal distribution function Φ, is uniformly distributed on (0, 1). This was tested with the Kolmogorov–Smirnov (KS) distance *D* between the empirical distribution of *u*_*k*_ and the uniform distribution. The mapping to uniformity is the same idea as in the time-rescaling diagnostics of point-process models, but it is not an application of the the supplementary material). *D* was compared with the approximate recording-specific critical value 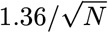 time-rescaling theorem: the innovations of SPARK are Gaussian, so the transform is Φ and not 1 − *e*^−Λ^ (see (approximate because the parameters are estimated from the same data; its behavior under a correctly specified model was examined by simulation). The distribution of *D* across recordings is reported instead of dichotomous *p*-value thresholds. Serial dependence was assessed with the partial autocorrelation function (PACF) of *z*_*k*_ over 40 lags, with bounds at 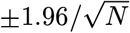, using beats with an artifact multiplier below 1.5 (“Goodness-of-fit diagnostic” section of the supplementary material).

### 2.7 Identifiability analyses

#### Model specifications

To evaluate whether RR alone identifies the decomposition and whether physiological information reduces the ambiguity, four nested specifications were compared (Table 2). The specification *M*_3_, which combines all the physiological information, is used for the fit diagnostics and for the perturbation, aging and exercise analyses. All specifications retain the structural constraints of Table 1, so that the comparison does not include label switching between the two drives. In *M*_0_ and *M*_1_, *λ*_*P*_ is a free parameter, whereas in *M*_2_ and *M*_3_ it is fixed by the spectral anchor. The specifications are alternative assemblies of the engine’s own likelihood and log-prior terms, and the compiled filter used inside the optimizer was verified against the engine’s filter (supplementary material).

**Table 2:**
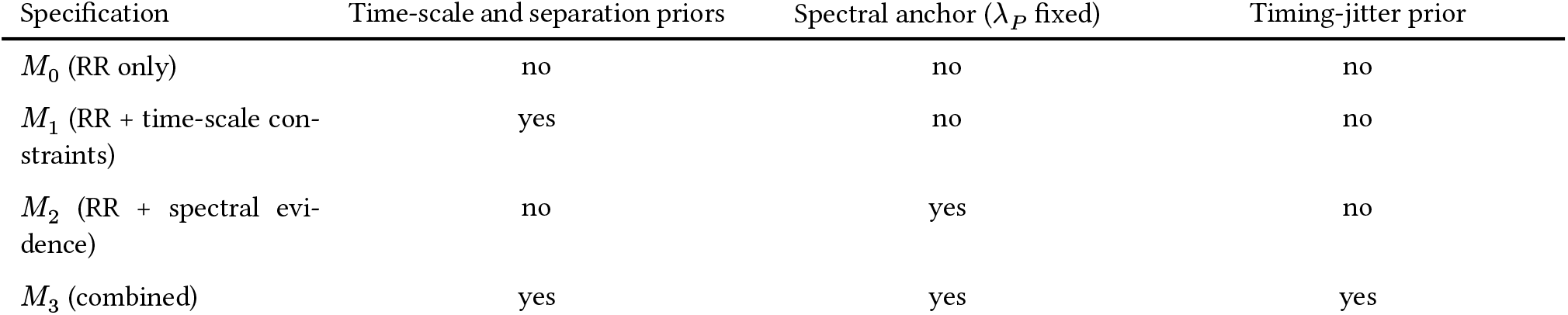
Model specifications used in the identifiability analyses.

| Specification | Time-scale and separation priors | Spectral anchor (*λ*<sub>*P*</sub> fixed) | Timing-jitter prior |
|---|---|---|---|
| *M*<sub>0</sub> (RR only) | no | no | no |
| *M*<sub>1</sub> (RR + time-scale con-straints) | yes | no | no |
| *M*<sub>2</sub> (RR + spectral evi-dence) | no | yes | no |
| *M*<sub>3</sub> (combined) | yes | yes | yes |

#### Dispersed multi-start fits

Each of *M*_0_–*M*_3_ was fitted by BFGS from the same set of *n*_*start*_ starting points (100 in this study; the first is the engine default and the others are drawn log-uniformly over wide ranges), in a deterministic random subset of up to 100 recordings per dataset (balanced across groups in the Fantasia database). For each recording and specification we computed the number of *distinct solutions*, the *near-equivalent fits* (profiled log-likelihood within 2 units of the best) with the dispersion of ln *τ*_*p*_, ln *τ*_*s*_ and of the reconstructed trajectories among them, the *share of x*_*p*_ *variance* and the proportion of fits in which one process absorbed essentially all variability (share below 0.05 or above 0.95; *one-process collapse*), the proportion of fits inside the 95% prior intervals, and the fit to RR; definitions are in the “Identifiability and sensitivity analyses: details” section of the supplementary material. Because the priors alter the objective, specifications are compared on the profiled log-likelihood and RR error, and not on the posterior objective.

#### Curvature, recovery under known truth and sensitivity

The numerical Hessian of the objective at the best fit of *M*_0_ and *M*_3_ was decomposed, and profile surfaces of the objective were computed over a 100 × 100 log-spaced grid of (*τ*_*p*_, *τ*_*s*_) for a subset of recordings. Synthetic RR series with known drives were generated in four scenarios (true time scales inside or outside the prior ranges, and vagal- or sympathetic-dominant variance) and fitted with each specification. Using *M*_3_ as the reference, each hyperparameter was varied one at a time and each prior component removed in turn, recording the change in *τ*_*p*_ and *τ*_*s*_, in the reconstructed states and in RR error (supplementary material).

### 2.8 Data, preprocessing and event definition

#### Data and cleaning

Each recording is a table of RR intervals derived from the electrocardiogram (ECG; single lead, recorded with a Polar H10 chest strap in the exercise cohort), with their times and, when available, blood pressure, respiration (from the Fantasia database) or the tilt-table angle (from the Physiologic Response to Changes in Posture, PRCP, database). RR intervals and covariates were cleaned with an adaptive running-median/median-absolute-deviation gate, chosen on simulated series with injected artifacts because it altered few artifact-free beats (about 0.2%) while correcting most artifacts (about 99.7%); analyses that use a covariate were restricted to recordings with at least 80% valid values in the analysis window. The latent state model itself uses only the RR intervals. Preparation, cleaning and their checks are described in the “Data preparation and its checks” section of the supplementary material.

#### Participants

The PRCP database contains ten healthy adults (the database reports neither age nor sex). The Fantasia database contains 20 young (21–34 years) and 20 older (68–85 years) healthy adults, with equal numbers of men and women in each group as reported by the database. The exercise cohort consists of community-dwelling adults aged 60 years or older from the Magallanes region of Chile, recruited for the study that produced these recordings, where the sample is described [33]; 263 recordings were analyzed. Sex and gender were not used as covariates in any analysis: the only subject characteristic analyzed was group membership in Fantasia.

#### Events

Every tilt and every stand-up of the PRCP protocol is an event analyzed separately against its own baseline. Onsets and returns were taken from the tilt-angle signal where available, from the protocol annotations otherwise, and, for recordings with neither and for the exercise recordings, from the heart-rate series itself: heart rate in 5-s bins was segmented into piecewise-constant levels, with the number of segments chosen by the Bayesian information criterion subject to a minimum difference of 0.10 Hz (about 6 bpm) between adjacent levels. Heart rate alone cannot tell a tilt from another maneuver that raises it, so such events are candidates, not confirmed tilts. Because they are detected from the same RR series that is modeled, timing relative to them is not independent evidence about the states, and the analyses therefore emphasize the relative timing and amplitude of the two components. The algorithms and their accuracy are given in the “Event detection” section of the supplementary material.

### 2.9 Validation framework: orthostatic perturbation

The first biological test of the reconstructed states uses tilt-table recordings from the PRCP dataset [34]. The aim is to assess whether the inferred components change in the expected direction during a perturbation. The components are named vagal-like and sympathetic-like because of their kinetics and spectral allocation (see *Model assumptions and identification structure*); a perturbation test can show that their behavior is physiologically coherent, but it cannot establish that either one is the parasympathetic or the sympathetic branch. Expected responses, based on the physiology of orthostatic stress, are a reduction of the vagal drive soon after the perturbation onset and a slower, more sustained increase of the sympathetic drive. Each maneuver was analyzed against its own baseline (60 to 180 s, beginning 120 s after the previous maneuver returned), followed by the maneuver itself and a 120-s recovery window. We computed (i) the difference in the mean of each state between phases, standardized by the baseline standard deviation (SD) of that state; (ii) whether each difference had the expected sign; and (iii) a step-response fit of each state, *x*(*t*) = *x*_0_ + *D* 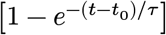 for *t* ≥ *t*_0_, yielding the response latency, the time constant and the relative onset lag *t*_0,*s*_ − *t*_0,*p*_ (fits at the search bounds are flagged). Between-recording summaries used one value per recording, the median over its tilts, so that a recording with more tilts did not weigh more; the share of individual tilts with the expected pattern is reported as a measure of within-recording consistency.

As a negative control, the same contrast was computed between the two halves of the resting baseline of each tilt, where no perturbation separates them, and compared with the tilt contrast (shares with exact 95% intervals and a paired signed-rank test of the absolute standardized changes over recordings; definition in the supplementary material). Reconstructions are displayed as time series with 95% intervals, as averages of tilts aligned on the onset (each relative to its own baseline) and as trajectories in the (*x*_*p*_, *x*_*s*_) plane in units of the baseline SDs. Secondary analyses (temporal resolution against a moving spectrum from the continuous wavelet transform, CWT, the arterial-pressure response and its lagged correlation with the states, the dose–response over tilt angle, and the responses by kind of maneuver) are described and reported in the “Secondary analyses of the perturbation” section of the supplementary material.

### 2.10 Generalization: aging

To examine whether the representation differs between young and older adults, we analyzed resting recordings from the Fantasia database [35], using only membership of the young or older group (taken from the record names) and a fixed analysis window (600 s starting at 600 s of the recording); no further covariates, such as continuous age or sex, were available. The question is which dimensions of the model-based description differ between groups, and not whether any single parameter is associated with age: the intrinsic rate *v*_0_, the time constants *τ*_*p*_ and *τ*_*s*_, the log-ratio ln(*k*_*p*_/*k*_*s*_), the relative vagal variance *λ*_*P*_, the global scale 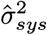, the equivalent timing jitter, the SD of each reconstructed state and their ratio, and the share of *x*_*p*_ variance. Group differences (older minus young) are summarized by the Hodges–Lehmann shift, the Wilcoxon rank-sum *p*-value and Cliff’s *δ* with a percentile bootstrap interval, with Benjamini–Hochberg control within each family of variables (latent and conventional HRV). Because the kinetic estimates are shaped by informative priors, group differences in *τ* are interpreted relative to those priors. Stability was examined by refitting two non-overlapping windows of each recording and by the intraclass correlation coefficient (ICC(3,1)) of the log-transformed values. With the group sizes available, all tests are exploratory. The respiratory and blood-pressure analyses of the same recordings are described and reported in the “Additional analyses of the Fantasia recordings” section of the supplementary material.

### 2.11 Exercise and recovery

The functional demonstration uses a two-minute step test with a rest–exercise–rest protocol in older adults [33]. Each recording is fitted individually, with time-invariant kinetic parameters, and the non-stationary exercise response is represented by the time-varying filtered states. The rest level of each state is its mean (and SD) from 20 s after the start to 5 s before the detected onset; the exercise response is the difference between the mean of the state over the last 15 s of exercise and the rest level (also expressed in rest SDs), together with the peak excursion; and the geometry of the (*x*_*p*_, *x*_*s*_) path, in units of the rest SDs, is summarized by its length, the area it encloses and its maximum quadratic energy *E* = (*u*^2^ + *v*^2^)/2, where *u* and *v* are the two states in units of their rest SDs (half the squared distance from the origin, the homeostatic baseline). These geometric summaries are not established statistics of HRV analysis; they are descriptive measures of the response introduced for this representation: the length is the distance travelled in the state plane, the enclosed area is the extent of the region that the path encloses, and the maximum energy reflects the largest distance from the rest point. No clinical or functional outcome measures and no subject characteristics were available for this cohort, and therefore no analyses of association with such variables are performed.

*Recovery time*. Because no single definition of recovery is canonical, four candidate definitions and three rules for the end of exercise were compared on simulated rest–exercise–recovery recordings with known recovery constants and end times, and the best-scoring combination was used; recordings that did not meet the criterion within the observation window were treated as right-censored (Kaplan–Meier medians are reported). The benchmark depends on the simulated profiles and is therefore a methodological check of the definitions, and not evidence about the real recordings (“Recovery-time benchmark” section of the supplementary material).

### 2.12 Conventional HRV comparators

Conventional HRV measures are used as comparators and not as competitors: the standard deviation of RR intervals (SDNN), the root mean square of successive RR differences (RMSSD), mean heart rate, and LF and HF power from a tapered periodogram of the resampled tachogram (the latter only for windows of at least 120 s). We ask whether the latent representation carries information that these descriptors do not already encode. This is examined descriptively in two ways: through rank correlations between latent summaries and HRV metrics within each dataset, and, in the perturbation and exercise recordings, through the relation between phase-specific HRV metrics and the corresponding latent responses. Leave-one-out analyses of how well HRV metrics predict the latent summaries, and of group discrimination in Fantasia, are described in the “Additional analyses of the Fantasia recordings” section of the supplementary material.

### 2.13 Statistical reporting

Quantities that vary across recordings are summarized as the median and the 2.5th–97.5th percentile range across recordings. This range describes between-recording dispersion; it is not a confidence interval for the median. Proportions are given with exact (Clopper–Pearson) 95% intervals; they are descriptive, since the tilts of one recording are not independent. None of the analyses was preregistered, and those on the PRCP, Fantasia and exercise recordings are exploratory. Fixed random seeds are used throughout, and the analysis code, configuration and session information are provided with the supplementary material.

## 3 Results

### 3.1 RR alone is insufficient for the latent decomposition

We first asked whether the RR series alone is sufficient to identify the latent vagal and sympathetic drives. To address this, the model was fitted without physiological information (*M*_0_) from dispersed starting values, and the resulting parameter estimates, likelihood values and reconstructed trajectories were compared across starts, in a subset of 150 recordings drawn from the three datasets. Evidence for under-determination would consist of solutions with comparable fit to RR but different latent decompositions, including solutions in which one process absorbs essentially all variability.

Without physiological information (*M*_0_; 150 recordings), the multi-start fits of a recording reached a median of 23.5 distinct solutions, with 60.0 fits within 2 log-likelihood units of the best, and one process absorbed essentially all variability in 75% of the fits. Under the full specification (*M*_3_), the corresponding values were 1.0 distinct solutions, 100.0 near-equivalent fits and one-process absorption in 4% of the fits. The dispersion of ln *τ*_*s*_ among near-equivalent fits under *M*_3_ was lower than that under *M*_0_ (0.00 versus 0.01; larger under *M*_0_ in 3 of 3 datasets: exercise, fantasia and prcp); it was small under both specifications, so the contrast rests mainly on the number of distinct solutions and on one-process solutions. The best RR error under *M*_3_ was higher than that under *M*_0_ (3.39 versus 1.00 ms; a factor of 3.4, small in absolute terms but not in relative terms). In these recordings, the RR series alone therefore left the latent decomposition less well determined than the constrained specification, consistent with the premise that physiological information is needed to restrict the admissible solutions. This shows that the constraints reduce the ambiguity; it does not show that the retained solution is the correct decomposition. The solutions reached from dispersed starting points are shown in Figure 1.

**Figure 1:**
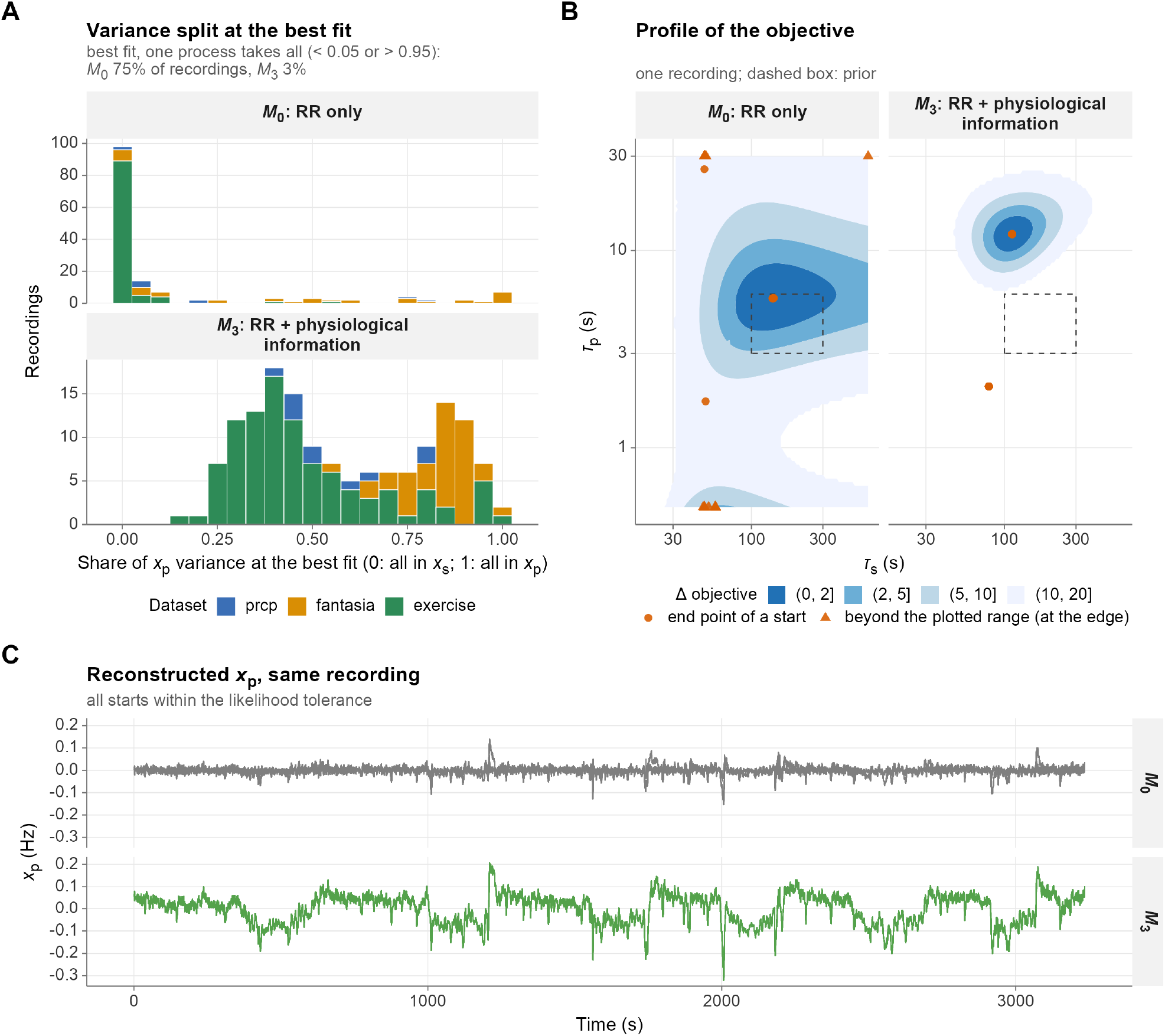
Non-identifiability under RR alone. (A) Share of *x*_*p*_ variance at the best fit of each recording, under *M*_0_ (RR only) and *M*_3_ (with physiological information); values at 0 or 1 mean that one process absorbs essentially all variability (the share of recordings is for the best fit only; the text reports the share of all fits). The single solution under *M*_3_ is conditional on the spectral anchor and the priors. (B) Profile of the objective over (*τ*_*s*_, *τ*_*p*_) for one recording, with the end points of its starts and the prior interval (dashed box); white areas inside the plotted range are grid points without a finite value. Under *M*_3_, 21 of the 100 starts of this recording stopped at a local optimum 191 log-likelihood units below the best fit (outside the contours); the others reached the same optimum. (C) Reconstructed vagal-like state *x*_*p*_ of the near-equivalent starts of the same recording.

### 3.2 Physiological information reduces solution ambiguity

The question is whether the constraints restrict the admissible solutions without imposing an arbitrary trajectory. This is assessed by comparing *M*_0_–*M*_3_ for dispersion of solutions, stability of the reconstructed trajectories, location of parameters within physiological ranges, and the change in RR fit (Table 3). Improved likelihood is not used as the criterion, since a well-chosen constraint can reduce likelihood while reducing ambiguity. As a caveat, constraining the model is expected to cost some RR fit; this cost is reported below and not hidden.

**Table 3:** Ambiguity metrics of the multi-start fits by specification (median [2.5th, 97.5th percentile] across recordings; for the two proportions of fits, one-process collapse and fits within the prior intervals, the mean across recordings [2.5th, 97.5th percentile of the per-recording values]). Large upper percentiles of the dispersion of ln *τ* come from recordings in which near-equivalent fits end at vanishingly small or very large time constants, a symptom of the likelihood ridge; they indicate that the fits are not pinned down, not their magnitude.

| Metric | *M*<sub>0</sub> | *M*<sub>1</sub> | *M*<sub>2</sub> | *M*<sub>3</sub> |
|---|---|---|---|---|
| Distinct solutions | 23.50 [6.72, 56.38] | 2.00 [1.00, 4.00] | 20.50 [6.00, 51.10] | 1.00 [1.00, 2.00] |
| Near-equivalent fits | 60.00 [1.00, 100.00] | 96.00 [15.87, 100.00] | 91.00 [1.00, 100.00] | 100.00 [44.90, 100.00] |
| SD of ln *τ*<sub>*p*</sub> (near-equiv.) | 0.00 [0.00, 18.60] | 0.00 [0.00, 0.08] | 0.05 [0.00, 19.63] | 0.00 [0.00, 0.00] |
| SD of ln *τ*<sub>*s*</sub> (near-equiv.) | 0.01 [0.00, 3.13] | 0.00 [0.00, 0.03] | 2.83 [0.00, 14.61] | 0.00 [0.00, 0.00] |
| Range of *x*<sub>*p*</sub> variance share | 0.00 [0.00, 1.00] | 0.00 [0.00, 0.04] | 0.29 [0.00, 0.99] | 0.00 [0.00, 0.00] |
| One-process collapse | 75% [15%, 99%] | 83% [0%, 100%] | 23% [0%, 100%] | 4% [0%, 59%] |
| Within prior intervals | 4% [0%, 50%] | 77% [0%, 100%] | 1% [0%, 1%] | 18% [0%, 100%] |
| Best RMSE (ms) | 1.00 [0.00, 65.56] | 3.10 [0.00, 65.58] | 3.16 [0.00, 66.98] | 3.39 [0.02, 65.75] |

The median SD of ln *τ*_*s*_ among near-equivalent fits was 0.01 for *M*_0_, 0.00 for *M*_1_, 2.83 for *M*_2_, 0.00 for *M*_3_ (lowest under *M*_3_). The best RR error differed by at most 2.40 ms (240%) between specifications, and the share of fits with both time constants inside the 95% prior intervals was 4% for *M*_0_, 77% for *M*_1_, 1% for *M*_2_ and 18% for *M*_3_. One process absorbed essentially all variability in 75% under *M*_0_, 83% under *M*_1_, 23% under *M*_2_ and 4% under *M*_3_ of the fits. Time-scale priors alone (*M*_1_) therefore did not prevent one-process solutions, whereas the specifications that include the spectral anchor (*M*_2_, *M*_3_) largely did; the lower ambiguity under *M*_3_ thus comes largely from fixing the variance ratio of the two drives through the anchor, and it should not be read as evidence that the apportionment between the drives is correct. The *M*_3_ estimates lay inside the 95% prior interval in 19% (*τ*_*p*_) and 99% (*τ*_*s*_) of recordings, with a median posterior contraction of 0.59 and 0.57 for ln *τ*_*p*_ and ln *τ*_*s*_, respectively. For the vagal-like drive, most estimates therefore lay outside the 95% prior interval, so the data moved *τ*_*p*_ away from the central region of its prior; this may reflect slower fluctuations being absorbed by the faster drive, and it limits the reading of *τ*_*p*_ as a strictly vagal time constant. In simulations with known truth, when the true time constants lay outside the prior intervals, the median absolute log-error of *τ*_*s*_ was 1.32 for *M*_0_ and 0.98 for *M*_3_ (the *M*_3_ error was lower than the *M*_0_ error); when they lay inside, 1.40 and 0.19. These values delimit the claim: constraints that help when the truth is compatible with them can bias the estimates when it is not. The 95% intervals of the filtered *M*_3_ states, which are conditional on the MAP parameters, contained the true state at 70% to 100% of the time points across the four scenarios; the lowest coverage was that of *x*_*s*_ in the sympathetic-dominant variance scenario (70%).

For *M*_3_, optimization converged for 313 of 313 recordings (100.0%), and the following summaries refer to the converged fits (exercise (n = 263), fantasia (n = 40), prcp (n = 10)). The filtered reconstruction of RR had a mean absolute percentage error (MAPE) of 0.222% [0.003, 5.544] and an RMSE of 2.611 ms [0.025, 89.252] (median [2.5th, 97.5th percentile] across recordings); because the implied RR at each beat uses states updated with that beat, these values describe in-sample filtered fit and not predictive accuracy. The innovation diagnostics indicate that the assumptions of serially independent, Gaussian innovations are met only partly in the recordings: the KS distance had a median of *D* = 0.056 [0.023, 0.177] against a mean recording-specific critical value of 0.0376 (approximate, because the parameters are estimated from the same data), *D* was below its critical value in 21.7% of recordings, compared with 0% exceeding it in well-specified simulations, and the innovation PACF departs from white noise (13.0 lags outside the approximate bounds on average; Supplementary Figure S10). Residual structure of this kind plausibly reflects components that the model does not represent, for example respiratory and baroreflex oscillations, and the reconstructed states should be read as conditional on this misspecification.

The estimated clearance rates were 0.07 s^−1^ [0.02, 0.38] for the vagal drive, corresponding to an e-folding time of 15.05 s [2.61, 44.17], and 0.007 s^−1^ [0.005, 0.009] for the sympathetic drive, corresponding to 150.0 s [112.6, 220.9] (Supplementary Figure S10, panel D). These estimates should be interpreted as MAP estimates under informative physiological priors and an enforced ordering *k*_*p*_ > *k*_*s*_. The separation of the two time scales is therefore a property of the identification structure and the data jointly, and does not by itself constitute independent evidence that the two drives have been segregated “blindly”. The extent to which the data move the estimates away from the prior centers is quantified by the contraction and prior-contrast values reported above.

### 3.3 Latent states during orthostatic perturbation

Having examined whether physiological information reduces ambiguity, we next asked whether the resulting states exhibit expected responses to an independently characterized perturbation. For the PRCP recordings, the tilts were taken from the tilt-angle signal where available (10 of 10 recordings) and from the protocol annotations otherwise. Each recording contained several maneuvers (40 tilts and 20 stand-ups analyzed in total). The filtered states of one recording are shown in Figure 2 (panel A) together with heart rate, arterial pressure and tilt angle, and the average response to a tilt in panel B; whether the components follow the expected pattern across recordings is summarized below as a test of physiological coherence; it should not be treated as proof that they correspond to the activity of the respective autonomic branches.

**Figure 2:**
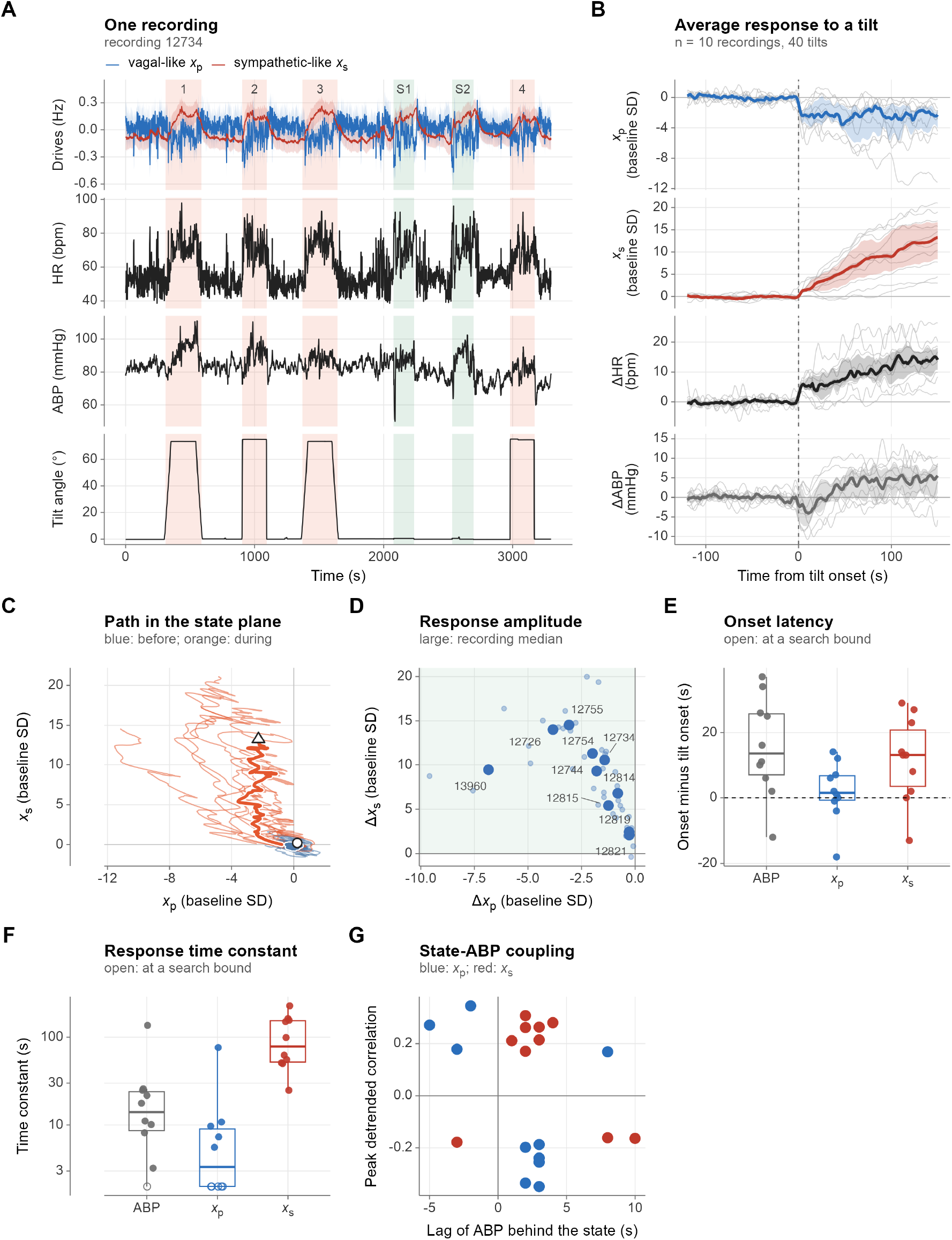
Latent states during repeated orthostatic perturbation. (A) One recording (recording 12734, median standardized response): filtered drives (blue, vagal-like *x*_*p*_; red, sympathetic-like *x*_*s*_) with 95% intervals conditional on the MAP parameters, heart rate, mean arterial pressure (ABP) and tilt angle; each tilt is shaded and numbered, and each stand-up (green) is labeled S. (B) All tilts aligned on their onset, each relative to its own baseline (states in baseline SDs; heart rate and ABP as changes); the line is the median over recordings, the band the quartiles and the grey lines the individual recordings. (C) Median path of the two states in the (*x*_*p*_, *x*_*s*_) plane in baseline SDs. (D) Standardized change of the two states between baseline and tilt (large points: recordings; small points: individual tilts); changes of *x*_*s*_ reach tens of baseline SDs because the baseline SD of *x*_*s*_ is small. (E) Onset of the response of each state, and of ABP when available, relative to the tilt onset (open symbols: at a search bound). (F) Step-response time constants. (G) Peak lagged correlation between each state and ABP after detrending, with its lag (the sign differs between recordings, so it is not a consistent coupling). When recordings with several tilt-angle levels are available, a further panel shows the mean states, heart rate and ABP by angle. The accuracy of the heart-rate-based events is shown in Supplementary Figure S5.

In 10 recordings with a defined perturbation (40 tilts, each analyzed against its own baseline; a recording contributes the median over its tilts), the vagal-like state decreased and the sympathetic-like state increased in 10 of 10 (exact 95% interval 69% to 100%), which is consistent with the expected pattern of vagal withdrawal with sympathetic engagement. The same pattern was present in 98% of the individual tilts. As a negative control, the same contrast applied to the two halves of each tilt’s resting baseline gave the expected pattern in 13 of 40 baselines (32%; exact 95% interval 19% to 49%), against 39 of 40 tilts (98%; 87% to 100%), so the pattern is clearly less frequent without a perturbation; the median absolute standardized change was 7.3 times larger for *x*_*p*_ and 8.2 times larger for *x*_*s*_ during the tilt (exact paired signed-rank test over 10 recordings, one-sided: *p* < 0.001 for both). The 25% expected for two independent random signs lies within this interval, so slow drift within the baseline is possible but is not demonstrated. Because heart rate rises during tilt, the net change of the two states is largely fixed by RR; what the spectral anchor and the priors determine is how that change is apportioned between the two drives. The sympathetic-like response began a median of 11.0 s after the vagal-like response, and its time constant was higher than that of the vagal-like response (80.7 versus 3.8 s); 70% of the *x*_*p*_ and 50% of the *x*_*s*_ step-response fits reached a search bound, so these constants are approximate. With 10 recordings and with events partly defined from the same RR series, this is a test of physiological coherence and not evidence that the components correspond to the activity of the respective autonomic branches.

The reconstructed dynamics of one recording, the average response and the group-level summaries are shown in Figure 2. Responses by kind of maneuver (slow tilt, rapid tilt, stand-up), the arterial-pressure response and its lagged correlation with the states, the dose–response over tilt angle and the comparison with a CWT moving spectrum are reported in the supplementary material (sections “Secondary analyses of the perturbation” and “Responses by type of maneuver”).

### 3.4 The latent representation across healthy aging

Next, we asked whether the representation is stable across individuals and whether it differs between young and older adults. The aging analysis addresses which dimensions of the model-based description differ between young and older adults in the Fantasia database, and not simply whether a given parameter differs. Only group membership was available as a subject characteristic.

Among 20 young and 20 older adults, 8 of 11 latent variables reached |*δ*| ≥ 0.33: *τ*_*p*_ (higher in older adults, *δ* = 0.72; adjusted *p* < 0.05), *λ*_*P*_ (lower in older adults, *δ* = −0.53; adjusted *p* < 0.05), 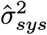 (lower in older adults, *δ* = −0.71; adjusted *p* < 0.05), the SD of *x*_*p*_ (lower in older adults, *δ* = −0.81; adjusted *p* < 0.05), the SD of *x*_*s*_ (lower in older adults, *δ* = −0.61; adjusted *p* < 0.05), the SD ratio *x*_*p*_/*x*_*s*_ (lower in older adults, *δ* = −0.54; adjusted *p* < 0.05), ln(*k*_*p*_/*k*_*s*_) (lower in older adults, *δ* = −0.69; adjusted *p* < 0.05) and the share of *x*_*p*_ variance (lower in older adults, *δ* = −0.54; adjusted *p* < 0.05); 2 of 4 HRV variables reached |*δ*| ≥ 0.33: SDNN (lower in older adults, *δ* = −0.71; adjusted *p* < 0.05) and RMSSD (lower in older adults, *δ* = −0.81; adjusted *p* < 0.05). These latent summaries are not independent: the share of *x*_*p*_ variance is a monotone function of the SD ratio of the two states, both depend strongly on the spectral anchor *λ*_*P*_, ln(*k*_*p*_/*k*_*s*_) combines the two time constants, and 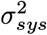 and the SDs of the states scale with the overall amplitude of variability, so that the count above overstates the number of distinct dimensions in which the groups differ. Agreement between two non-overlapping windows of the same recording (ICC of the log values) had a median of 0.88 for the latent summaries and 0.93 for the conventional HRV metrics; the lowest agreement was that of *τ*_*s*_ (0.31). With these group sizes, all comparisons are exploratory, and no continuous age, sex or other covariates were available. Group contrasts are shown in Figure 3 and listed in the “Additional analyses of the Fantasia recordings” section of the supplementary material, together with the respiration and blood-pressure measures.

**Figure 3:**
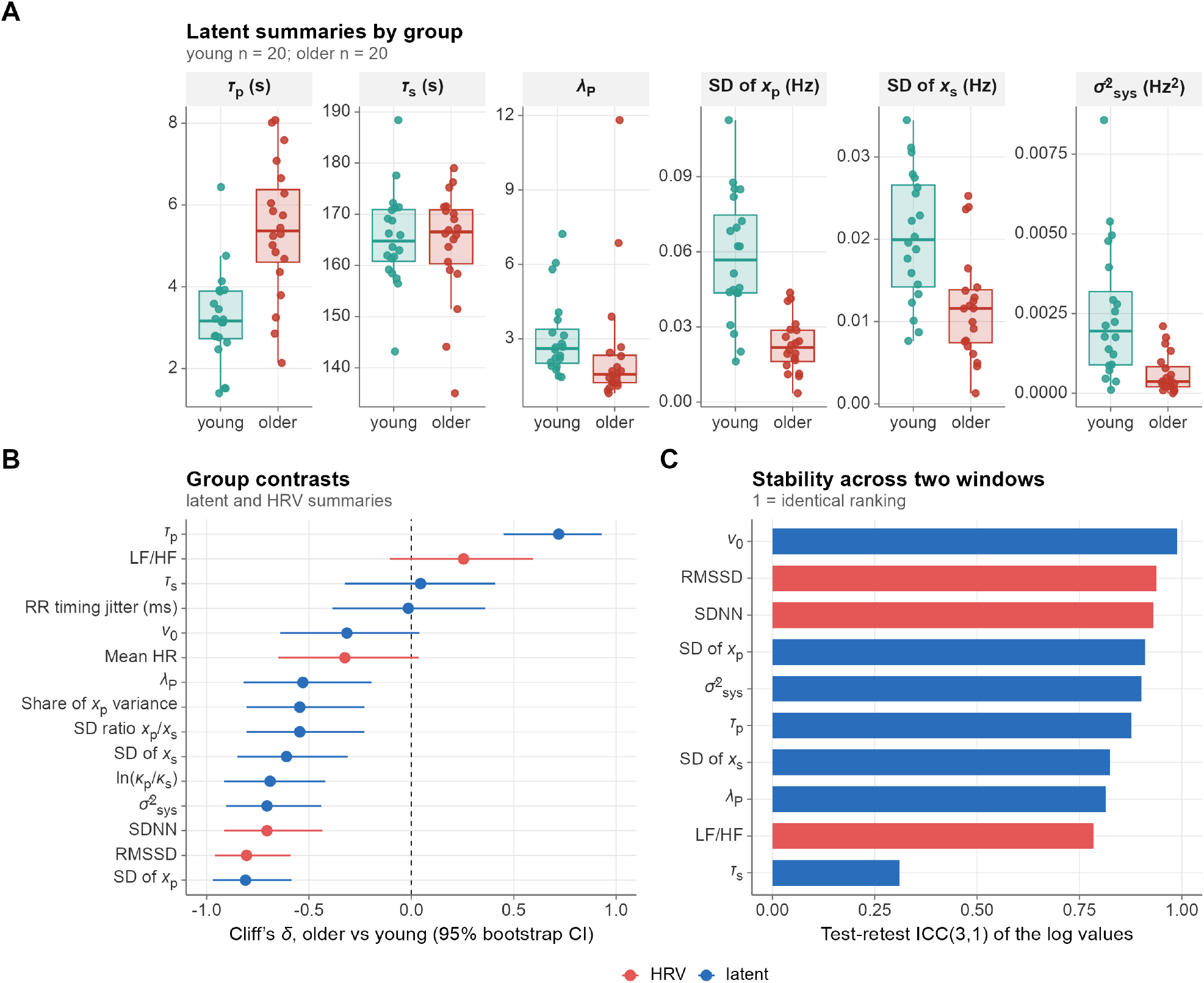
Latent representation in young and older adults. (A) Distributions of latent summaries by group. (B) Effect sizes (Cliff’s *δ*, older versus young) for latent and conventional HRV variables. (C) Test-retest agreement (ICC(3,1) of log values) between two non-overlapping windows.

### 3.5 Latent dynamics during exercise and recovery

Finally, we asked whether the reconstructed trajectories characterize adaptation and recovery during exercise. In the two-minute step-test recordings, the reconstruction tracked the changes in RR across rest, exercise and recovery (Figure 4, panel A). The rest, exercise and recovery phases were identified from heart rate. The recovery definition (90% recovery of the excursion) and the end-of-exercise rule (offset time of the fitted exercise profile) were those selected by the simulation benchmark (Supplementary Figure S3).

**Figure 4:**
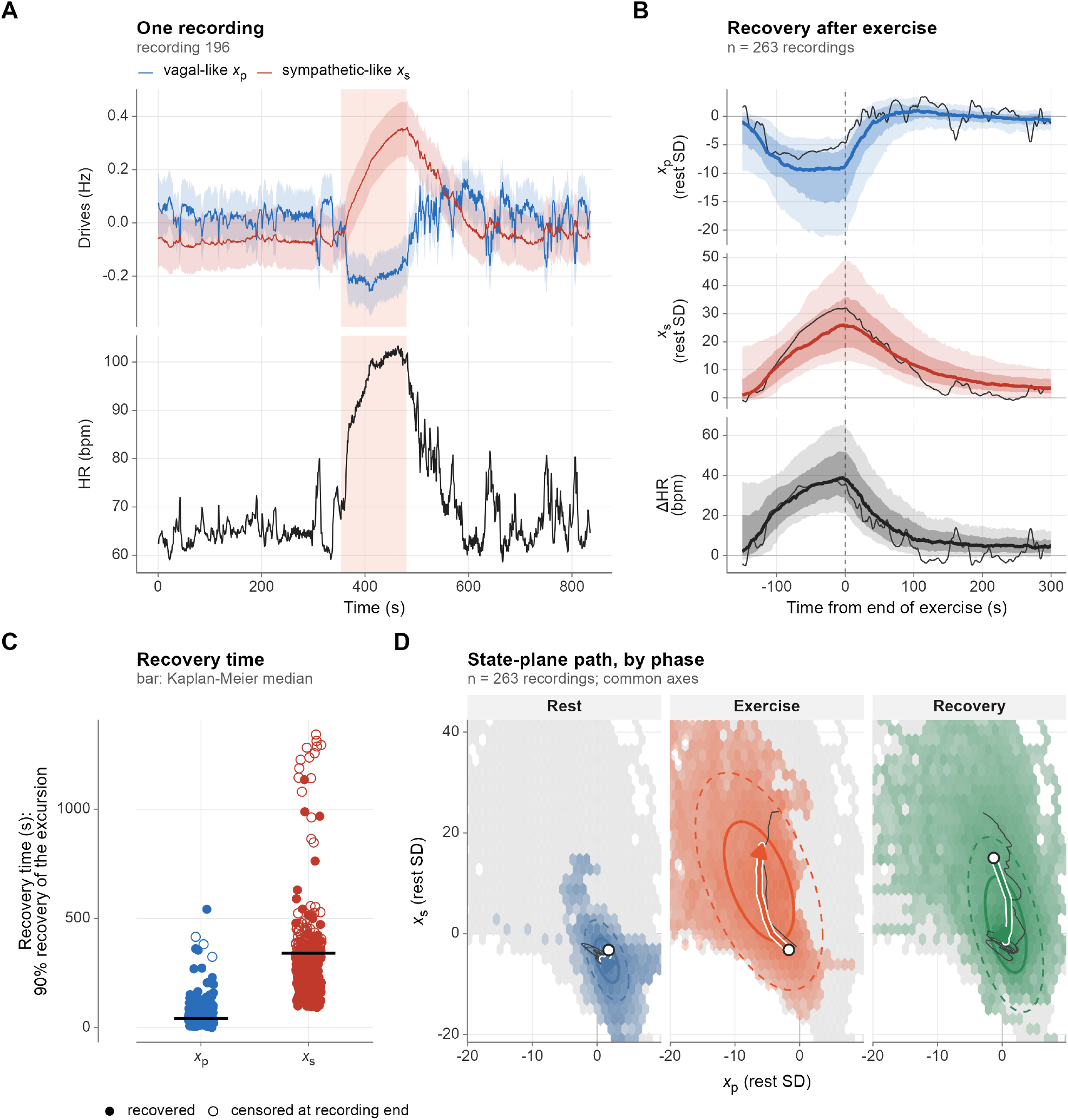
Adaptation and recovery in older adults. (A) One recording (recording 196, median standardized response): filtered states and heart rate, with the detected exercise shaded. (B) All recordings aligned on the end of exercise, each relative to its rest level (median, bands of 25–75% and 10–90% of recordings, and the recording of A). (C) Recovery times of each state (open symbols: right-censored at the end of the recording; bars: Kaplan–Meier medians). (D) Filtered states in the (*x*_*p*_, *x*_*s*_) plane in rest-SD units, one panel per phase on common axes (central 99% of the pooled points): density of the points of the phase (colored hexagons) over that of all phases (grey), ellipses holding 50% (solid) and 90% (dashed) of them, and the median path through the phase (arrow; open circle: start).

In 263 of 263 exercise recordings the rest, exercise and recovery phases could be defined from heart rate. From rest to the end of exercise, the vagal-like state changed by −7.9 [−26.9, −1.4] and the sympathetic-like state by 24.9 [5.5, 59.5] rest SDs, with a heart-rate increase of 40 [19, 117] bpm (median [2.5th, 97.5th percentile] across recordings). Recovery times (90% recovery of the excursion; Kaplan-Meier medians) were 42 s for the vagal-like state (1% of recordings censored) and 341 s for the sympathetic-like state (37% of recordings censored). For a first-order relaxation, recovery to 90% of the excursion takes about 2.3 *τ*, so these times are of the order implied by the estimated time constants, which are themselves shaped by their priors; they describe the fitted kinetics and are not independent measurements of recovery. The path in the state plane had a median length of 214 [95, 590], an enclosed area of 215 [38, 1051] and a maximum energy *E* of 253 [36, 1525] in rest-SD units. In the state plane, the strongest basin of the occupied region (the mode of the density of the filtered states) was displaced from its position in the rest phase by 9.3 rest-SD units to the exercise phase and 2.3 rest-SD units to the recovery phase (Supplementary Figure S6); the number of basins detected was 1 in the rest phase, 3 in the exercise phase, 1 in the recovery phase. The landscape is a descriptive map of where the states lie and is not a fitted potential of the model. Conventional HRV metrics predicted the latent summaries of this cohort with a median leave-one-out *R*^2^ of 0.39. The strongest rank correlation between a latent response and an HRV change was −0.65 (standardized change of the vagal-like state with the change in SDNN, n = 263). No subject characteristics or clinical and functional outcomes were available for this cohort, so the analysis is descriptive and no association with such variables is reported. The corresponding example recording is shown in Figure 4. The occupation of the (*x*_*p*_, *x*_*s*_) plane in each phase is summarized as a density in Supplementary Figure S6; this map is descriptive and is not a potential function estimated from the model.

## 4 Discussion

### 4.1 Principal observations

We examined whether a physiology-informed latent state-space model can reconstruct model-conditional autonomic dynamics from RR intervals when the latent decomposition is not guaranteed to be identified by the observations alone. The premise of the study is that a close fit to RR does not by itself guarantee a physiologically meaningful decomposition, so that independent physiological information is better treated as part of the identification structure than as an auxiliary refinement. In the recordings analyzed, adding the physiological information reduced the number of distinct latent solutions and the occurrence of one-process solutions relative to RR alone at a cost in RR error. In the datasets analyzed, the innovation diagnostics indicated residual serial structure or a departure from the assumed innovation distribution in most recordings, so that the reconstructions are conditional on a noise model that holds only approximately; with this caveat, in the tilt-table, aging and exercise analyses, the vagal-like state decreased and the sympathetic-like state increased in most tilt-table recordings; most of the partly redundant latent summaries differed between young and older adults; and the exercise recordings showed a rest-exercise-recovery trajectory whose recovery times are closely tied to the fitted time constants. These findings concern the behavior of a model-based representation under the stated assumptions and should not be taken as direct evidence about neural activity. The latent states have not been validated against an external measure of autonomic activity; the evidence is of internal consistency and of agreement with expected qualitative responses.

### 4.2 Physiological meaning of the latent trajectories

The latent trajectories are physiologically constrained reconstructions. They describe the drives that the model would need, given the kinetic and spectral properties assumed in the identification structure, to account for the observed interval timing. This is the sense in which the components are called vagal-like and sympathetic-like: their identity rests on the imposed kinetic ordering and spectral allocation and not on independent measurement. During orthostatic tilt, the reconstructed vagal-like state decreased and the sympathetic-like state increased in 10 of 10 recordings (100%), a pattern consistent with the expected autonomic adaptation to orthostatic stress. The same contrast was clearly less frequent in the resting baselines (negative control), where it reflects drift and not a perturbation. Representing adaptation as a trajectory in the (*x*_*p*_, *x*_*s*_) plane, with summaries such as path length or enclosed area, offers a compact description of response and recovery. The quadratic energy *E* used to summarize the path is computed from the states expressed in rest SDs and is a descriptive device, not an energy of the physiological system. Interpreting trajectory loops in terms of dissipation, entropy production or allostatic load would require independent validation and is not supported by the present analysis.

### 4.3 Relationship with conventional HRV

Conventional HRV metrics summarize the variability of the RR series and remain robust and widely understood descriptors of the output of the system. SPARK addresses a complementary question: rather than asking how variable RR is, it asks what latent dynamical state could account for how RR changes, and how that state responds, adapts and recovers. The two approaches are therefore descriptive and generative, respectively, and neither is intended to replace the other. Window-based spectral estimates trade temporal against frequency resolution under non-stationary conditions [5,11], whereas the filter updates beat by beat. A comparison with a CWT moving spectrum in the tilt recordings is given in the supplementary material (section “Secondary analyses of the perturbation”); the temporal behavior of the filter depends on its own assumptions, including independent drives and time-invariant kinetics within a recording, and does not establish general superiority. In the Fantasia recordings, HRV metrics predicted the latent summaries with a median leave-one-out *R*^2^ of 0.42, so that a substantial part of their variation was not linearly recoverable from SDNN, RMSSD and mean heart rate. Incomplete prediction from a limited set of conventional HRV metrics does not establish additional physiological information or incremental clinical value; such claims require comparison against broader benchmarks and independent outcomes.

### 4.4 Identification and physiological priors

In an underdetermined inverse problem, physiological knowledge is not merely an auxiliary source of prior information; it forms part of the identification structure of the model (Table 1): positivity and the ordering *k*_*p*_ > *k*_*s*_ are structural, the time-scale and separation priors restrict where kinetic solutions can lie, the spectral anchor allocates variance between the two drives, and the jitter prior limits the extent to which the noise term can absorb autonomic variability.

Two consequences follow from this design. First, the kinetic estimates are posterior modes under informative priors that are fixed across recordings. The estimates are regularized toward the prior distributions but are not bounded by their central 95% intervals, and differences between groups or conditions in *k* should therefore be interpreted together with the contrast between prior and posterior. The separation of fast and slow time scales is built into the identification structure and is not, by itself, an empirical finding. Second, the spectral anchor assigns branch energy by frequency and is computed from the same recording. Conditions in which the two branches overlap in frequency, such as strong orthostatic stress or exercise, may violate the assumed allocation and lead to misattribution between the drives [8,36]. The pharmacological evidence on which the spectral weights are based was drawn from the literature and was not generated here. These assumptions are explicit, and their influence is evaluated through the identifiability and sensitivity analyses. Varying the physiological hyperparameters one at a time changed the time constants by a median absolute log-ratio of at most 0.64 and left the reconstructed trajectories correlated at least 0.99 with the reference. The direction and time course of the reconstructed states were therefore robust to these settings. Correlations are insensitive to a change of scale, however, and do not address how variance is apportioned between the two drives: the share of latent variance attributed to the vagal-like process changed by a median absolute 0.23 at most, for the vagal floor (0.5), and by 0.37 when the spectral anchor was removed, so statements about the split between the two drives are conditional on the anchor settings (see the limitations).

### 4.5 Phase-domain formulation and relation to existing approaches

Point-process models of heartbeat intervals based on generalized linear structures are largely phenomenological with respect to the sinoatrial node [15,37], whereas state-space formulations of the integral pulse frequency modulation framework generally solve the nonlinear first-passage observation model in the time domain and rely on extended or unscented filtering [14,17,18]. By conditioning on the observed interval, SPARK obtains a Gaussian measurement equation that is linear in the integrated drives, together with exact propagation of the Ornstein–Uhlenbeck dynamics across irregular intervals, which avoids discretization error in the transitions. Here the transitions are the transition kernels of the latent drives from one observed beat to the next: because the drives are Ornstein–Uhlenbeck processes, the distribution of the state at the next beat given the current one is Gaussian with a mean and covariance known in closed form (the matrix exponential Φ_*k*_ and its process covariance), so no Euler–Maruyama or other time-stepping approximation of the dynamics is needed, however irregular the intervals are. The price is the terminal-threshold approximation: the neglected boundary-reflection term is bounded in proportion to the ratio of diffusion to the square root of the drift (see the supplementary material). The bound is small in the high-drift, low-noise regime but is not zero, and it should be examined in recordings with large threshold noise; in the fitted recordings, *ε* = *s*_*RR*_/Δ*t* had a median of 0.0063 (maximum 0.0579), corresponding to a bound of at most 0.0116 on the CDF discrepancy.

Under the strict first-passage formulation, the variance of the interval scales with the inverse cube of the drift, in qualitative agreement with the usual observation that respiratory sinus arrhythmia is larger when vagal influence is high, although its amplitude is not a pure index of vagal tone (“Interval-variance scaling” section of the supplementary material). SPARK conditions on the observed interval and uses a Gaussian measurement equation whose variance does not depend on the vagal level, so this dependence is not represented and the stationary variances of the two drives are treated as constant within a recording.

### 4.6 Relation to nonlinear stochastic dynamics

Seen as a dynamical system, SPARK couples a fast and a slow mean-reverting stochastic process to a threshold-crossing phase oscillator that is reset at every event, and the data are the crossing times. Three aspects connect it to general questions of nonlinear stochastic dynamics. First, the observation is a first-passage problem: conditioning on the observed interval replaces it by a terminal-threshold condition whose error is bounded and small in the fitted regime (see *Phase-domain formulation*), so that an intrinsically nonlinear observation becomes tractable by exact linear-Gaussian filtering. The continuous-time drives evolve without interruption, and only the integral states, which accumulate phase, are reset to zero at each beat; the resulting system mixes continuous dynamics with discrete events at the beats. Second, in this context identifiability is a question about time-scale separation: the fitted rates of the two drives differ by a median factor of 10 in these recordings, and when the likelihood is left unconstrained it shows ridges along which solutions with very different time scales and variance partitions fit the intervals almost equally well (Section 3.1); external information on the time scales of the two drives and on how variance is distributed across frequency is what removes this degeneracy (neither source alone does: Table 3), and it does so by assumption and not by the data. Third, because the filter keeps the continuous-time trajectories of both drives, the response of a perturbed system relaxing towards its baseline can be described geometrically in the (*x*_*p*_, *x*_*s*_) plane (path length, enclosed area, occupied basins); we present these as descriptive summaries of the reconstructed trajectories, not as a fitted potential or an estimate of dissipation. Whether richer dynamics, such as state-dependent noise, time-varying kinetics or coupling between the drives, could be detected from RR intervals alone is an open question that requires a more complex model construction and an identifiability analysis similar to the one used here.

### 4.7 Limitations

Several limitations delimit the claims. The latent states are model-based reconstructions and are not direct measurements of autonomic neural activity; they have not been compared with microneurographic recordings or tested under pharmacological blockade, and the atropine-informed spectral evidence is external to the present data. Identification remains incomplete even with physiological information. The innovation diagnostics indicated residual serial structure or a departure from the assumed innovation distribution in most recordings (the KS distance was below its critical value in 22% of them), so the reported intervals are conditional on a noise model that holds only approximately. The vagal-like time constant lay inside its 95% prior interval in 19% of recordings, so it should not be read as a strictly vagal time constant: the faster drive may absorb slower fluctuations, such as baroreflex-related oscillations. The recovery times and the time constants of the two drives are shaped by their priors and describe the fitted kinetics; in recordings whose duration is comparable to or shorter than the sympathetic time scale, *τ*_*s*_ is likely to be informed mainly by its prior.

The hyperparameters are fixed across recordings, the spectral anchor is an approximation, and kinetics and stationary variances are assumed constant within a recording, which may be restrictive during strongly non-stationary exercise. The state intervals are conditional on the MAP parameters, the states are filtered and not smoothed, the optimizer is started from three fixed points and keeps the best run (a better local optimum cannot be excluded for an individual recording), and the simulation studies use data generated by the model itself. Finally, the apportioning of a perturbation-induced excursion between the two drives is not determined by the data alone: it depends on the spectral anchor, and in the sensitivity analyses the share of latent variance attributed to the vagal-like process changed by a median absolute 0.10 or more with the vagal floor, the spectral crossover and the *τ*_*p*_ prior interval, so statements about the relative contribution of the two drives are conditional on those settings. For the same reason the tonic level of each drive is not identified: *v*_0_ and any constant offset of *x*_*p*_ and *x*_*s*_ enter the likelihood only through their combination with the mean rate. The representation therefore supports comparisons of variability, kinetics and mean rate between people and times (summarized for the Fantasia recordings in the Supplementary Material), but not of the absolute vagal or sympathetic tone, which would need an external anchor such as autonomic blockade.

The data impose additional limits. Sex and other participant characteristics were not analyzed as covariates. The responses by type of maneuver rest on 20 slow tilts, 20 rapid tilts, 20 stand-ups (at most 2 events of a kind per recording) and are descriptive. The tilt-table dataset comprises a limited number of recordings (10 recordings), each with several tilts that are not independent of one another; events for the recordings without a tilt angle and for the exercise cohort were detected from the same RR series, so timing relative to them is not independent evidence about the states. The Fantasia database includes only group membership as a subject characteristic, and the exercise cohort consists of older adults with a single protocol and no subject-level characteristics or clinical outcomes, so the relevance of the latent trajectories to such outcomes was not evaluated and the behavior of the framework in populations with overt cardiac autonomic pathology remains untested [38,39]. The single-stream formulation does not include respiratory or blood-pressure inputs: respiratory sinus arrhythmia is captured only as an oscillatory component of the vagal state, and baroreflex feedback is implicit [8,40,41], so central autonomic commands cannot be distinguished from peripheral mechanical modulation.

### 4.8 Future directions

The most direct extensions address these limitations. Concurrent respiratory and arterial-pressure signals could be incorporated through explicit coupling terms [42–44], which may reduce reliance on the spectral anchor; validation against microneurography and graded pharmacological blockade [45,46] would test the interpretation of the latent components directly; and hierarchical formulations could learn the hyperparameters across recordings instead of fixing them. Because the continuous-time trajectories are preserved, quantities such as the geometry of the phase-plane path or the behavior of the kinetic parameters near critical slowing down could be explored [47,48], each requiring dedicated validation before a physiological interpretation. Datasets with clinical and functional outcomes and subject covariates would be required to determine whether the latent trajectories carry information that is relevant in those settings.

## 5 Conclusion

SPARK provides a physiology-informed framework for reconstructing latent cardiac dynamics from RR intervals. In the datasets analyzed, regularization and spectral anchoring improved solution stability and reduced one-process collapse at a cost in RR error, while the reconstructed states showed responses consistent with orthostatic and exercise-related changes. These findings support its use as an exploratory dynamical representation, but do not establish branch-specific physiological validity. Substantial residual structure, conditional uncertainty intervals, and sensitivity of variance allocation to the spectral anchor remain important limitations. Independent physiological validation is required before interpreting the components as quantitative measures of parasympathetic and sympathetic activity.

## CRediT authorship contribution statement

**Matías Castillo-Aguilar:** Conceptualization; Methodology; Software; Validation; Formal analysis; Investigation; Resources; Data curation; Writing – original draft; Writing – review & editing; Visualization. **Noah Beelders:** Conceptualization; Validation; Writing – review & editing. **Cristian Núñez-Espinosa:** Writing – review & editing; Supervision; Project administration; Funding acquisition.

## Ethics statement

The exercise cohort is a secondary analysis of recordings from a previously published study [33], which was approved by the Ethics Committee of the University of Chile (ACTA No. 029 − 18/05/2022) and the Ethics Committee of the University of Magallanes (No. 008/SH/2022). No new data were collected for the present work, and only the RR-interval series, identified by a recording number, were used, so no additional ethical approval was required. The PRCP and Fantasia recordings are publicly available, de-identified datasets from PhysioNet, analyzed secondarily under their open license (Open Data Commons Attribution 1.0).

## Code availability

The model is implemented in R (engine deposited with this article as model_code.R). An equivalent implementation runs entirely in the browser, without uploading data, at https://matcasti.github.io/SPARK-app/. In the five reference series tested, its likelihood, states and parameter estimates agree with those of the R engine (fitted parameters to within 10^-5^, relative; states, in Hz, to within 10^-6^). The application is intended for research use and exploration, not for clinical decisions.

## Data availability

The PRCP recordings (Physiologic Response to Changes in Posture; https://doi.org/10.13026/C2PC7R) and the Fantasia database (https://doi.org/10.13026/C2RG61) are publicly available from PhysioNet [49] under the Open Data Commons Attribution license; the repository downloads them with checksum verification (`make download`). The RR-interval series of the exercise cohort (263 recordings, without personal identifiers) are in the repository’s `data-raw/tmst` folder. The analysis code and its configuration are at https://github.com/matcasti/SPARK-model/, and the figures, tables and numbers of this article are regenerated from them (`make data`, `make run`, `make render`). The repository is public and under version control; the version used for this article is the release v1.0.0 (https://github.com/matcasti/SPARK-model/releases/tag/v1.0.0), which fixes the code and configuration at a dated commit and carries the result files of the article, whose checksums are listed in the repository (`results/MANIFEST.csv`). The release is not deposited in a repository that issues a DOI.

## Declaration of generative AI and AI-assisted technologies in the writing process

During the preparation of this work the authors used Claude (Anthropic) to assist with drafting and language editing of the text. After using this tool, the authors reviewed and edited the content as needed and take full responsibility for the content of the publication.

---

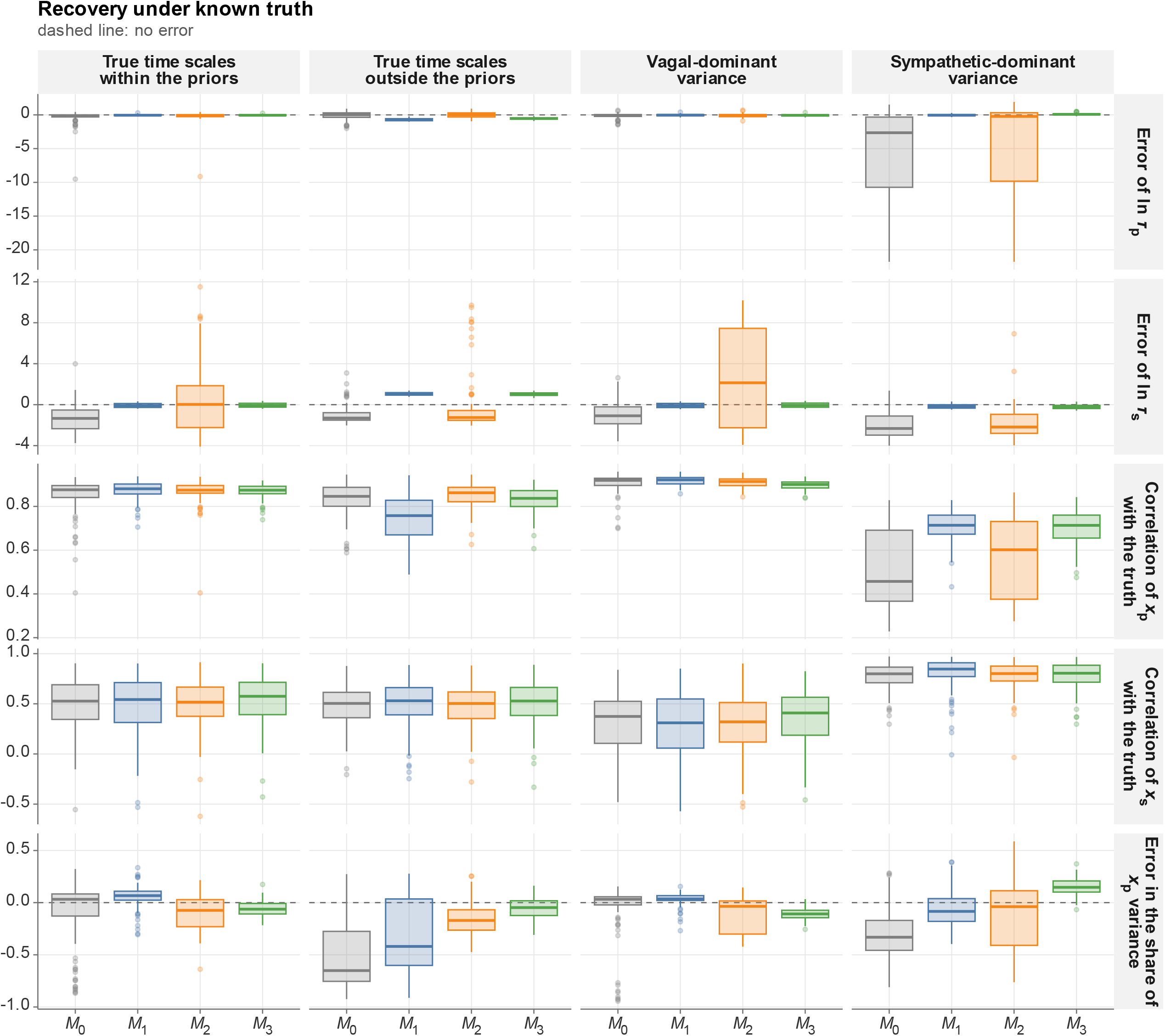

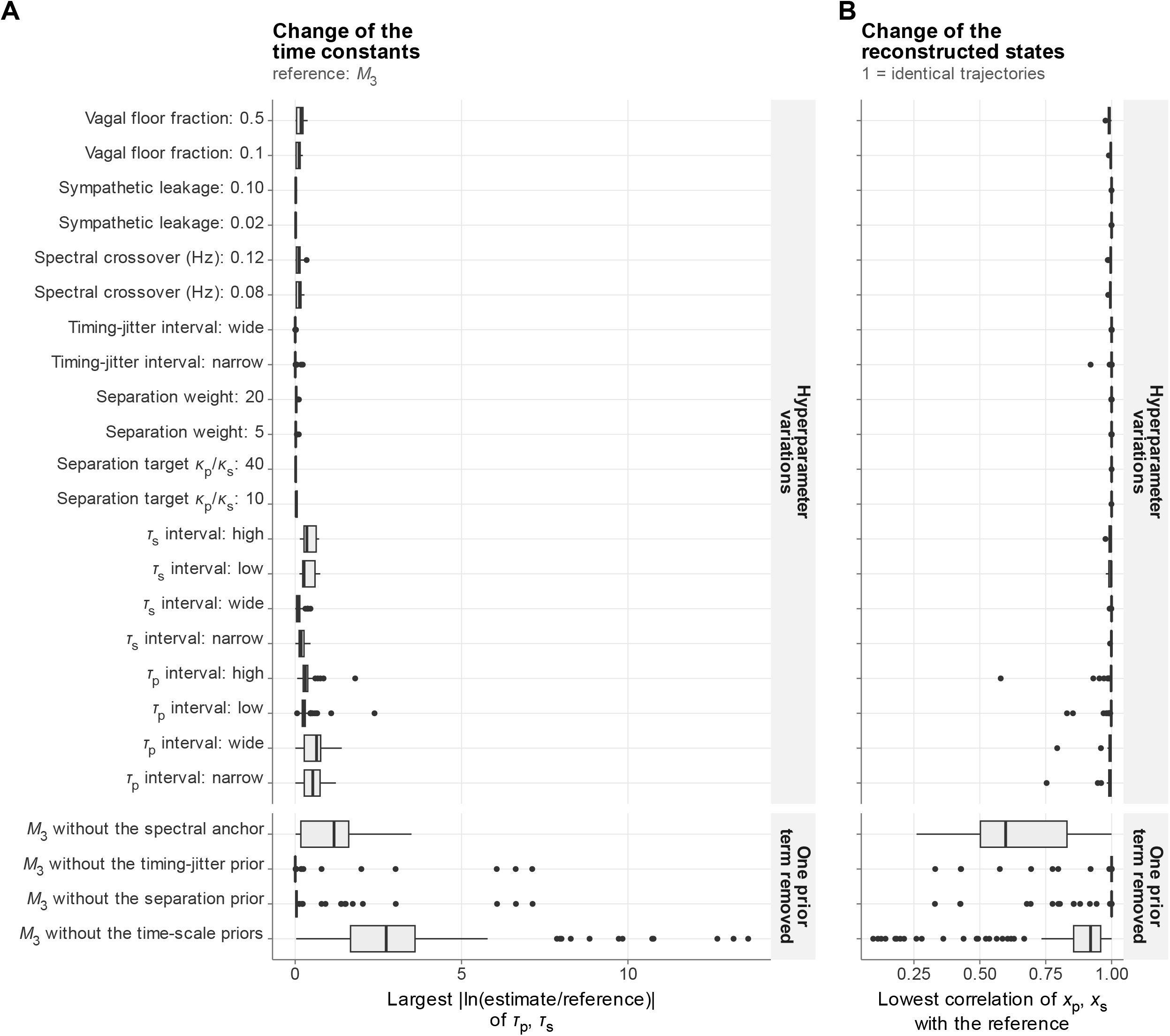

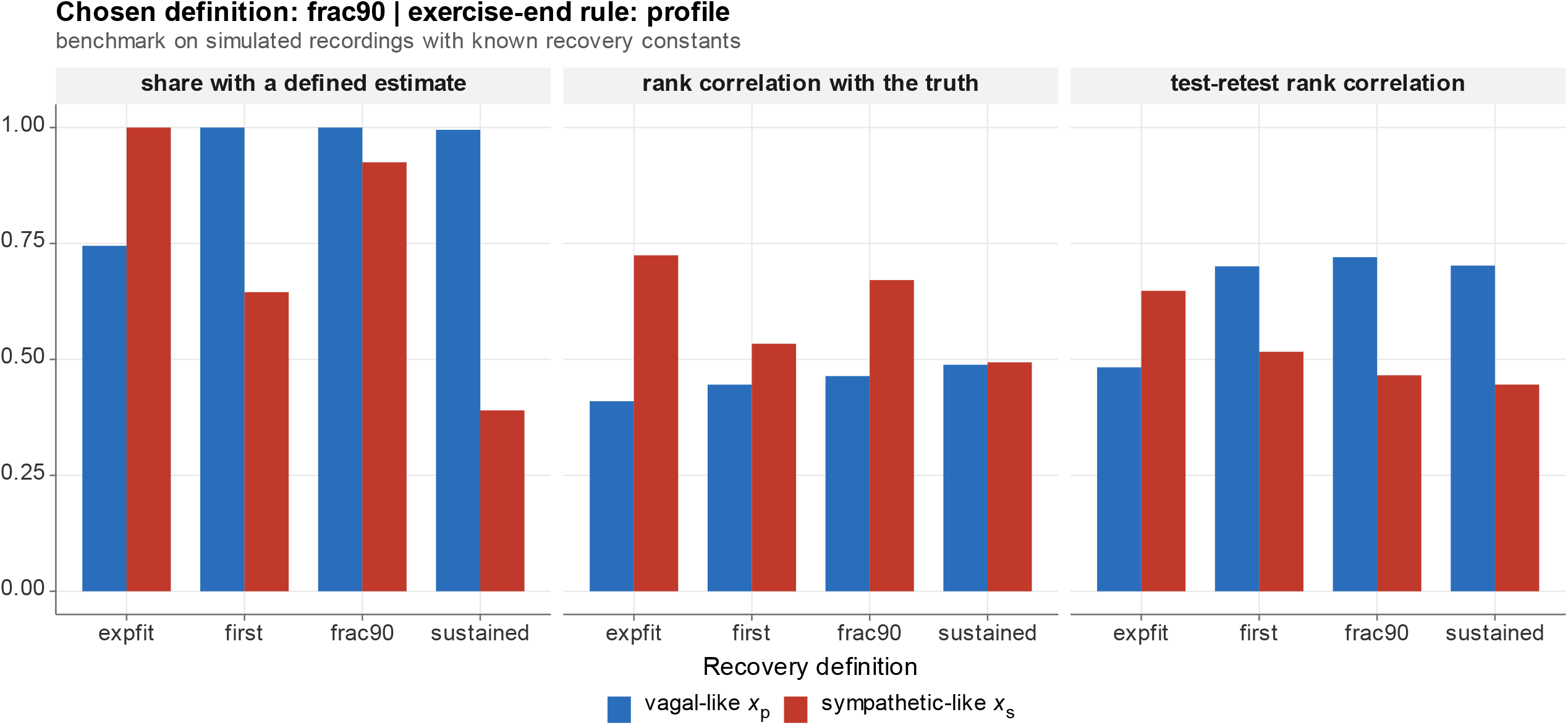

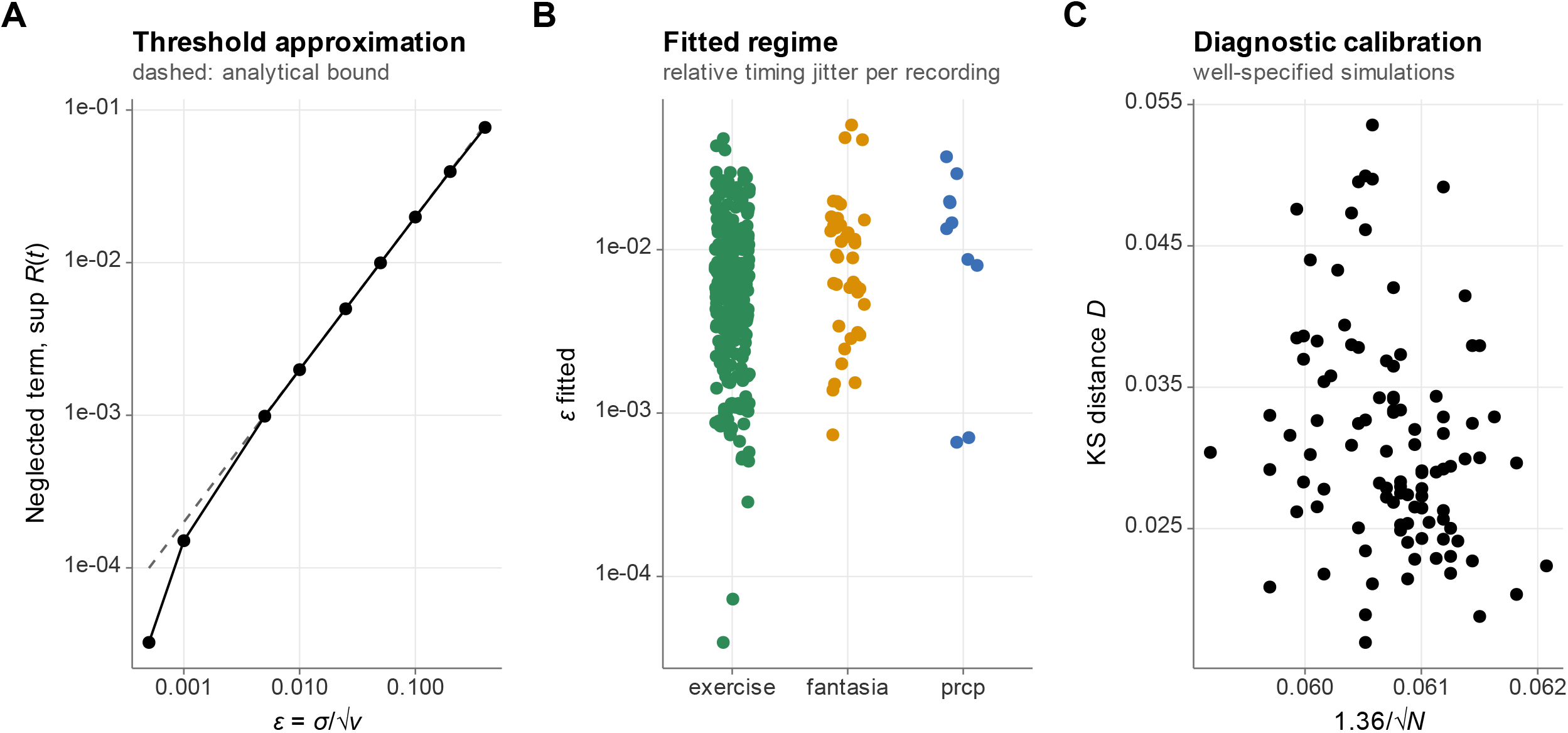

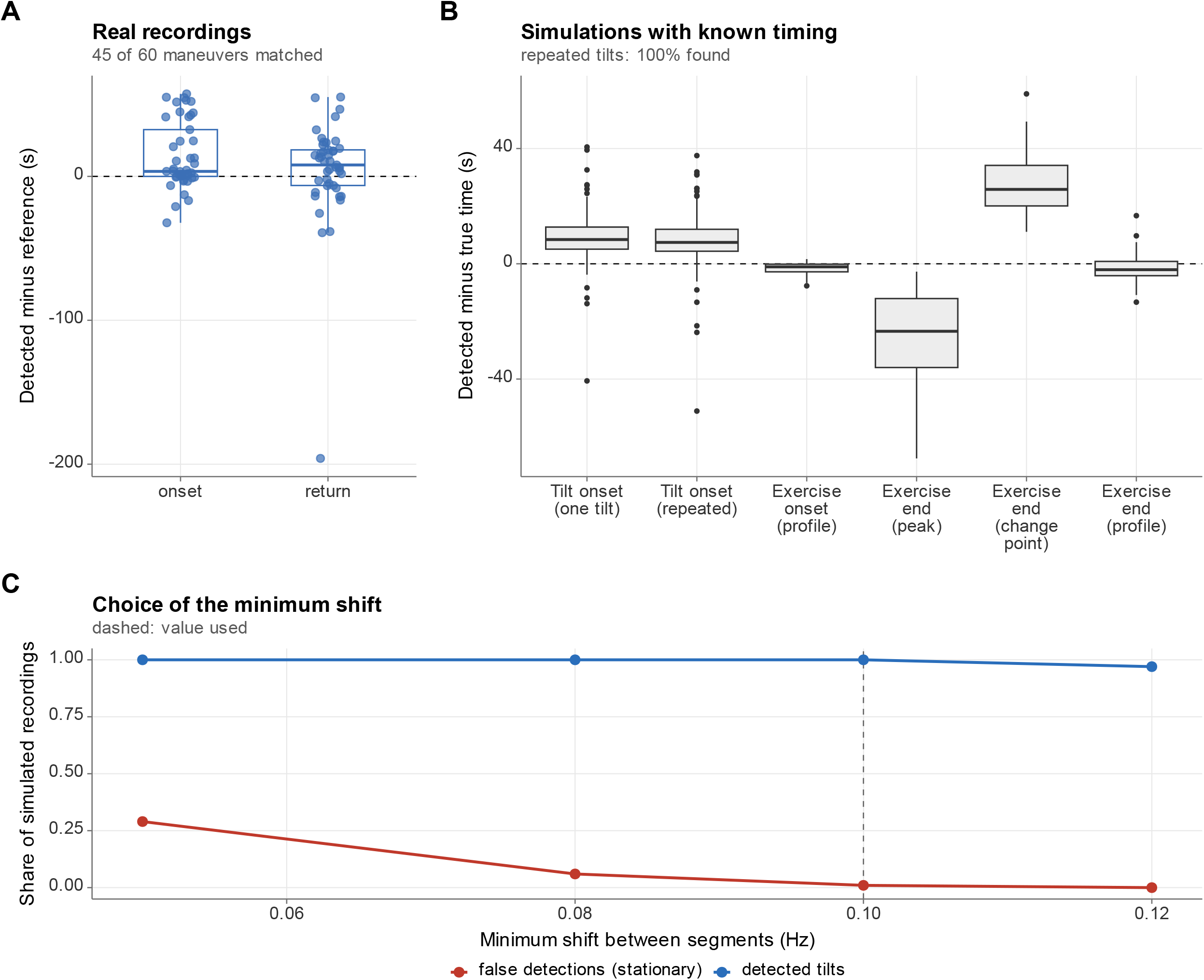

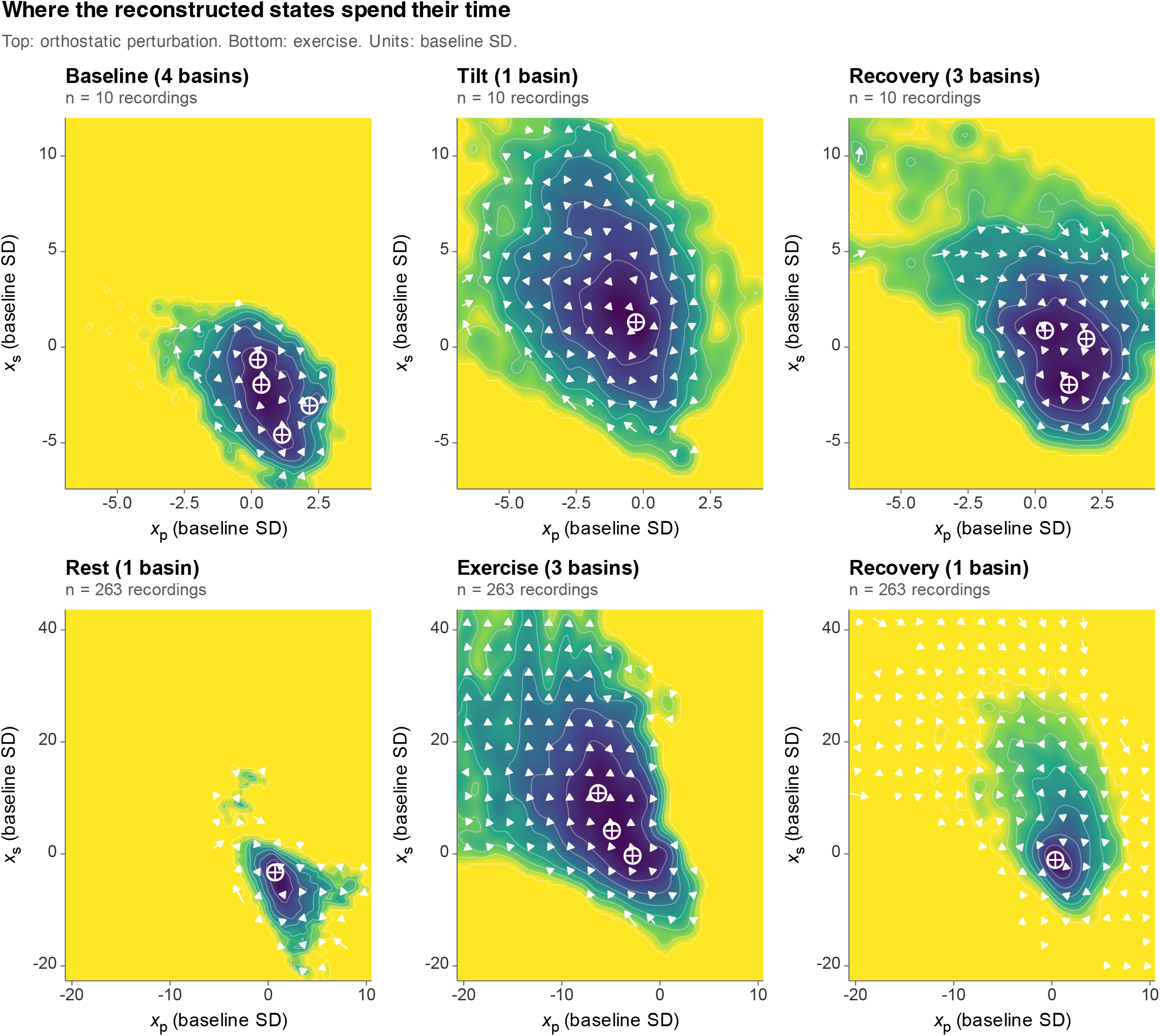

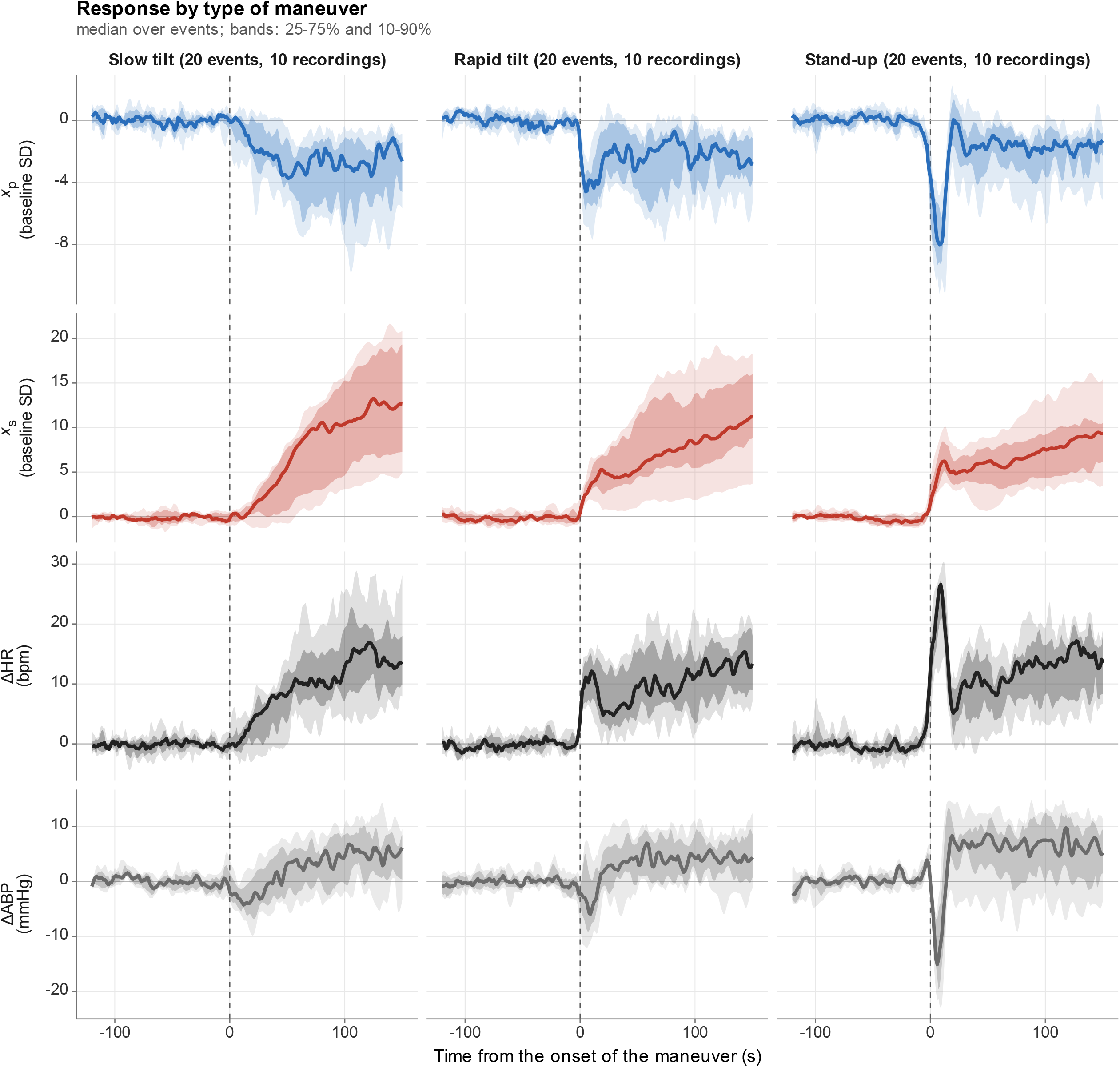

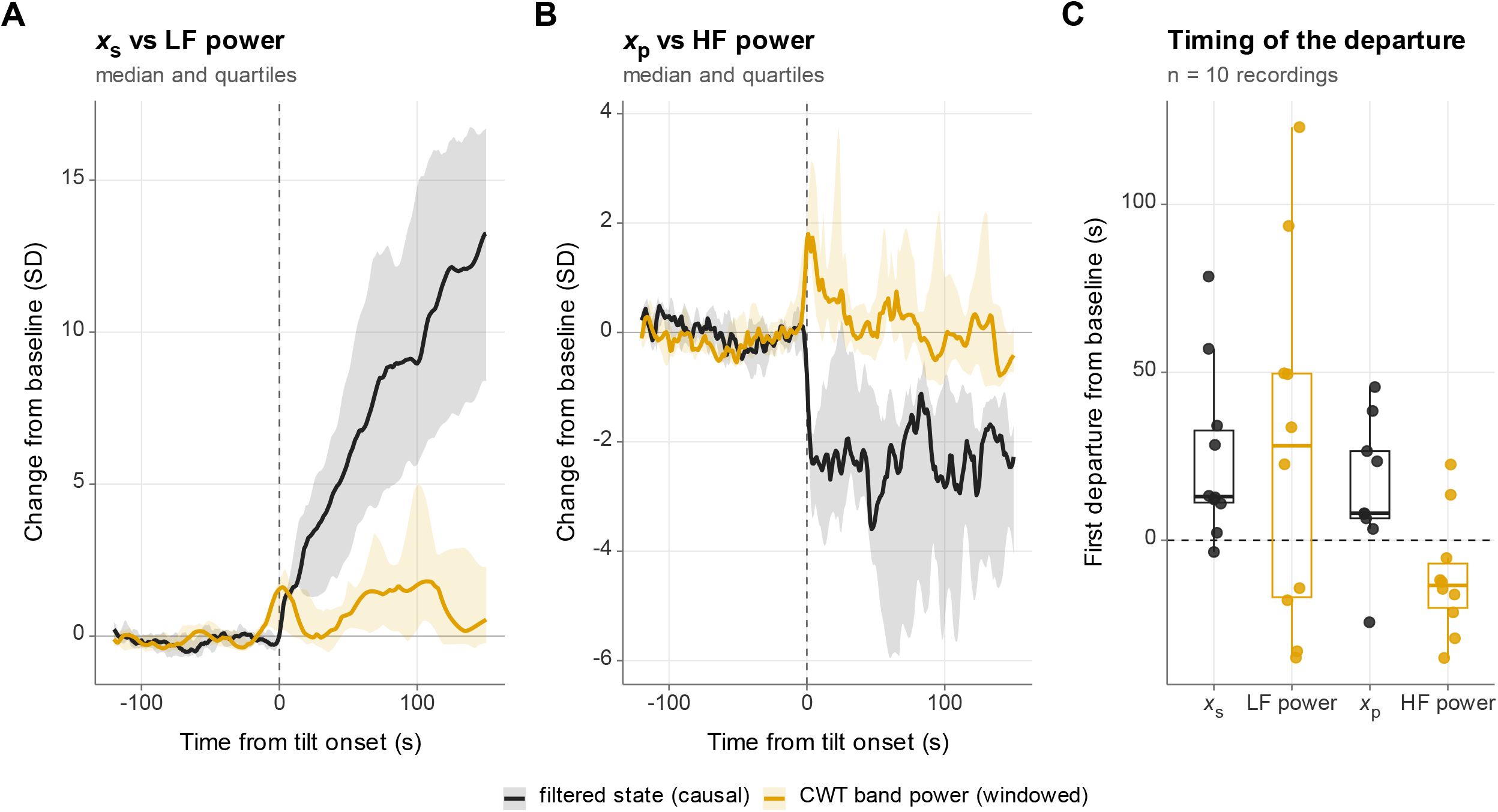

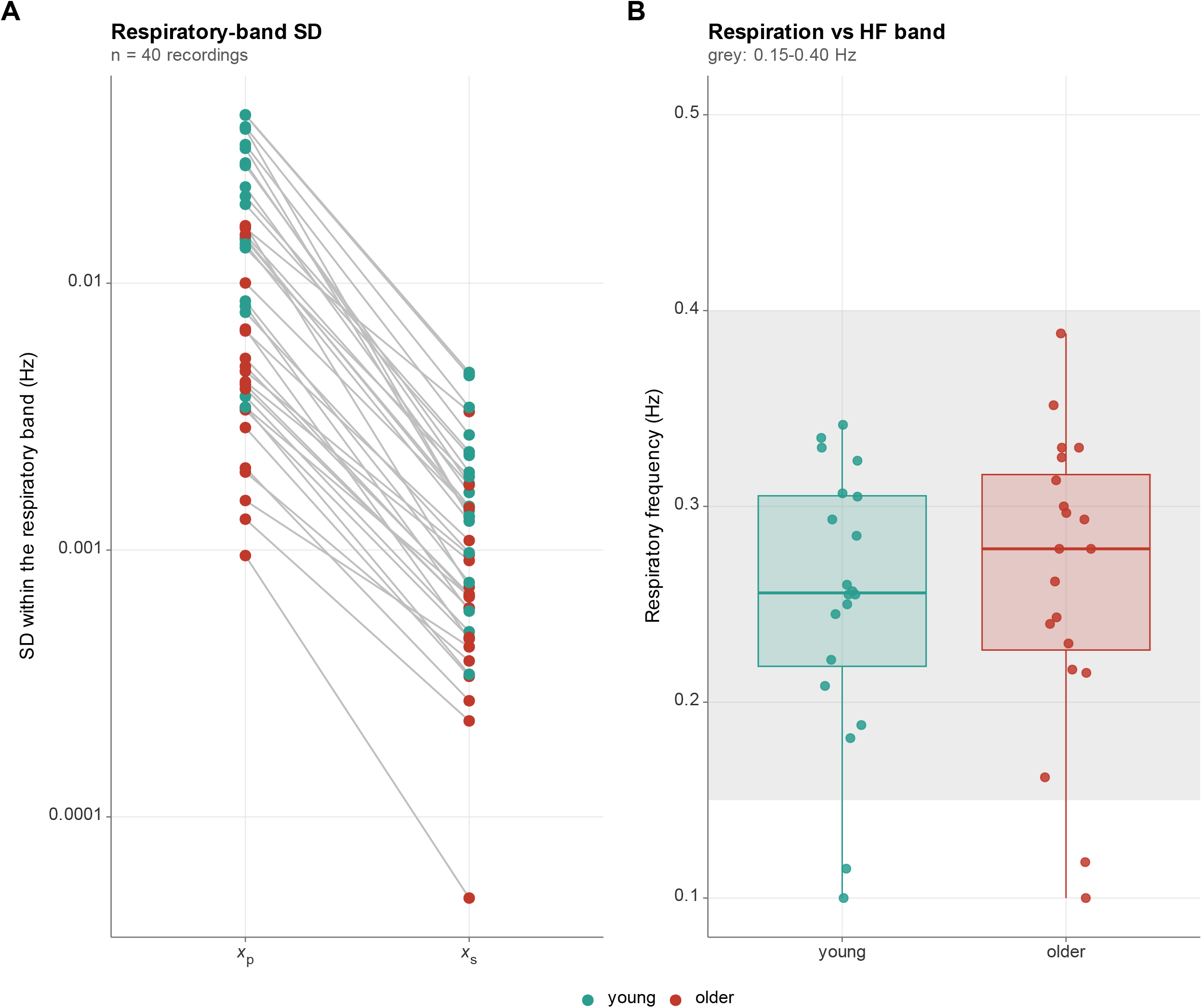

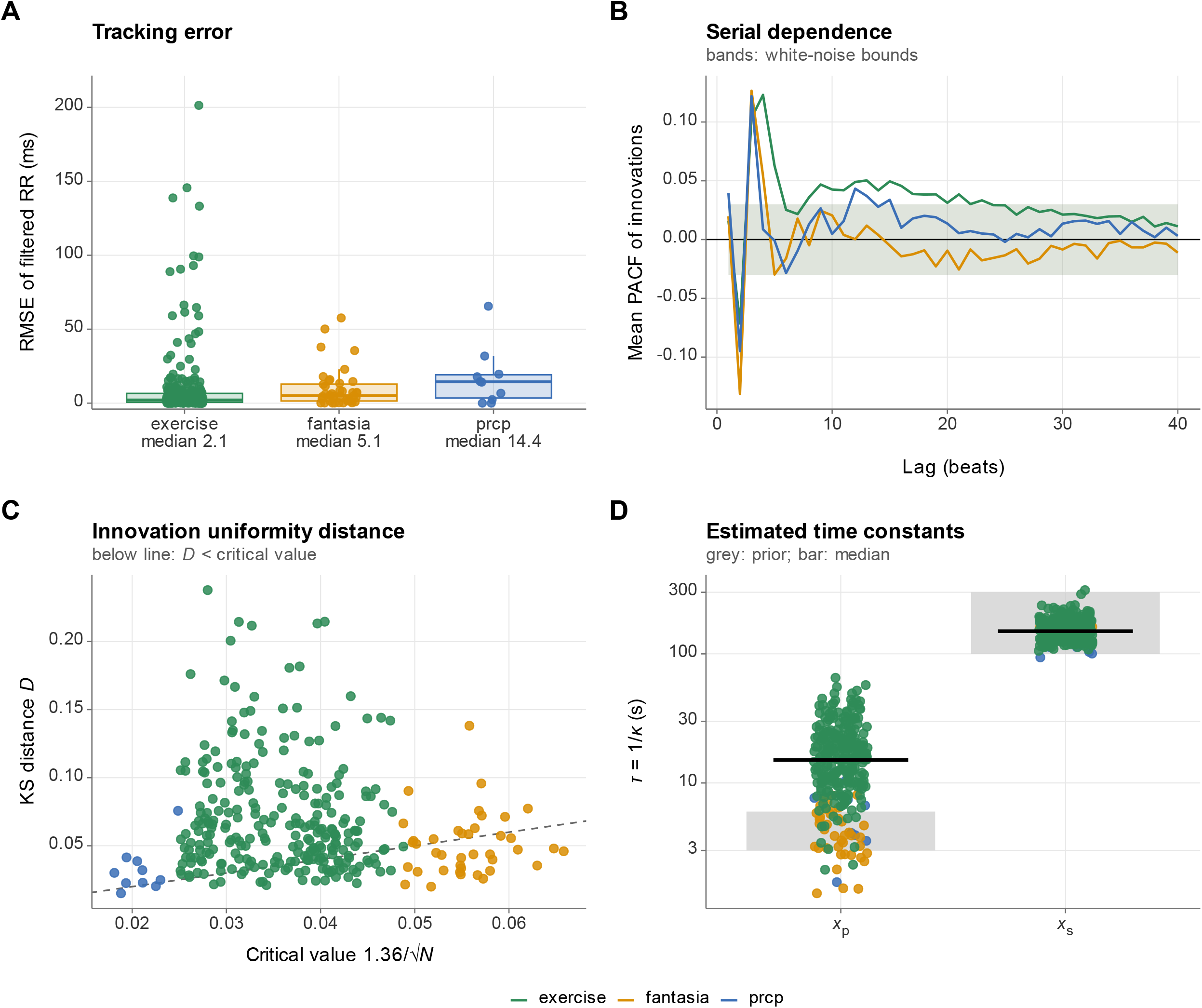

