## Supplementary material for "SPARK: Physiology-informed inference of latent cardiac autonomic dynamics from RR intervals"

### 1 Asymptotic convergence of the phase-domain mapping

The phase-domain mapping evaluates the accumulated phase over a deterministic temporal window $\Delta t_k$ and thereby relaxes the path-survival condition of a strict first-passage time (FPT) formulation. We show here that the discrepancy between the terminal-threshold and first-passage formulations is explicitly bounded and vanishes as the threshold noise becomes small relative to the drift, which is the regime expected for sinoatrial pacing.

Consider the closed-form cumulative distribution function (CDF) of the first-passage time $\tau$ of a one-dimensional drift–diffusion process with constant net drift $\nu > 0$, diffusion coefficient $\sigma$ and an absorbing boundary at $a = 1$:

$$F_\tau(t) = \Phi\left(\frac{\nu t - 1}{\sigma\sqrt{t}}\right) + \exp\left(\frac{2\nu}{\sigma^2}\right)\Phi\left(\frac{-\nu t - 1}{\sigma\sqrt{t}}\right),$$

where $\Phi(\cdot)$ is the standard normal CDF. By the reflection principle, the first term is the probability that the unconstrained process lies above the threshold at time $t$, which corresponds to the terminal-threshold observation equation used in the phase-domain inversion. The second term, which we denote $R(t)$, accounts for sample paths that crossed the boundary before $t$ and subsequently diffused back below it.

The magnitude of $R(t)$ can be bounded explicitly. Let $x = (\nu t + 1)/\left(\sigma\sqrt{t}\right) > 0$ and use the Gaussian tail inequality $\Phi(-x) \leq \varphi(x)/x$, where $\varphi$ is the standard normal density. Then

$$R(t) \leq \frac{1}{\sqrt{2\pi}}\frac{\sigma\sqrt{t}}{\nu t + 1}\exp\left(\frac{2\nu}{\sigma^2} - \frac{(\nu t + 1)^2}{2\sigma^2 t}\right) = \frac{1}{\sqrt{2\pi}}\frac{\sigma\sqrt{t}}{\nu t + 1}\exp\left(-\frac{(\nu t - 1)^2}{2\sigma^2 t}\right).$$

Because the exponential factor does not exceed one and $\nu t + 1 \geq 2\sqrt{\nu t}$, it follows that, uniformly in $t > 0$,

$$\sup_{t>0}\left|F_\tau(t) - \Phi\left(\frac{\nu t - 1}{\sigma\sqrt{t}}\right)\right| \leq \frac{\sigma}{2\sqrt{2\pi}\,\nu} = \frac{\varepsilon}{2\sqrt{2\pi}} \approx 0.2\,\varepsilon, \qquad \varepsilon = \frac{\sigma}{\sqrt{\nu}}.$$

The multiplier $\exp(2\nu/\sigma^2)$ offsets the Gaussian tail decay of $\Phi(-x)$ near the mean interval $t \approx 1/\nu$, so the neglected term is small because $\varepsilon$ is small and not because it is exponentially negligible; it decays rapidly only away from $t \approx 1/\nu$. The bound therefore vanishes as $\varepsilon \to 0$, that is, as the drift becomes large relative to the diffusion.

For a locally constant drift with unit threshold, the mean interval is $\Delta t \approx 1/\nu$ and the standard deviation of the first-passage time is $\sigma/\nu^{3/2} = \varepsilon\,\Delta t$. The parameter $\varepsilon$ is therefore approximately the relative timing jitter of the threshold crossing, $\varepsilon \approx s_{RR}/\Delta t$, where $s_{RR} = \sqrt{\hat{\sigma}^2_{sys}\lambda_R\,\Delta t}/\hat{\nu}_0$ is the equivalent RR timing jitter reported by the model. For jitter within the prior range of 0.5–20 ms and RR intervals near 0.8 s, $\varepsilon$ ranges from approximately $6 \times 10^{-4}$ to $2.5 \times 10^{-2}$, and the bound ranges from approximately $1 \times 10^{-4}$ to $5 \times 10^{-3}$ in CDF units. The terminal-threshold formulation is thus a close approximation of the first-passage distribution in this regime, but it remains a modeling assumption that should be verified when the estimated jitter is large.

#### 1.1 Numerical verification

The bound was verified numerically (step 11 of the analysis pipeline, Table S 12). Evaluating the neglected term $R(t)$ on a fine grid of $t$ for $\varepsilon$ between $5 \times 10^{-4}$ and 0.4, the ratio of its supremum to the analytical bound was at most 0.999 (a value of 1 or less confirms the bound, which is tight near $t \approx 1/\nu$). A Monte Carlo simulation of the drift–diffusion process (Euler steps of 2 ms, 4000 paths per setting) gave maximal differences between the empirical first-passage CDF and the closed form of 0.017 and 0.020 for $\varepsilon$ = 0.1 and 0.3 (Monte Carlo standard error at most 0.008); discretization of the simulated paths makes the empirical crossing time slightly later than the continuous-time one, so this comparison is a coarse check of the closed form and not of the bound itself.

In the fitted recordings, the relative timing jitter $\varepsilon \approx s_{RR}/\Delta t$ was 0.0063 [0.0007, 0.0297] (median [2.5th, 97.5th percentile] across recordings; maximum 0.0579), which corresponds to a bound on the CDF discrepancy of at most 0.0116. The engine reports the equivalent jitter in ms as `rr_jitter_ms`.

### 2 Estimation algorithm

This section details the algorithm implemented in the accompanying R engine. The filter evaluates the augmented state $\tilde{X}(t) = \left[x_p(t), x_s(t), I_p(t), I_s(t)\right]^\top$ over the observed intervals $\Delta t_k$, $k = 1, ..., N$. All variances in the recursion are expressed in units of the global scale $\sigma^2_{sys}$, which is restored after profiling. In these units the stationary variances are $V_s = 1$ and $V_p = \lambda_P \, g_k$, where $g_k = 1$ for the ungated model.

The quantities of the augmented state-space model and their estimation status are summarized in Table S 1.

Table S 1: Quantities of the augmented state-space model

| Symbol | Description | Estimation status |
|---|---|---|
| $x_p(t), x_s(t)$ | Latent vagal and sympathetic drives (Hz, deviations from baseline) | Reconstructed (filtered) |
| $I_p(t), I_s(t)$ | Running time-integrals of the drives since the last beat | Reconstructed (filtered) |
| $\nu_0$ | Baseline pacing rate (Hz): the mean rate at baseline | Profiled (GLS) |
| $\sigma^2_{sys}$ | Global variance scale | Profiled (concentrated ML) |
| $\kappa_p, \kappa_s$ $(\tau_p, \tau_s)$ | Clearance rates (e-folding times) of the vagal and sympathetic drives | Estimated (MAP) |
| $\lambda_P = V_p/V_s$ | Relative vagal-to-sympathetic stationary variance | Fixed from wavelet spectral partition |
| $\lambda_R$ | Threshold-jitter variance rate, relative to $\sigma^2_{sys}$ | Estimated (MAP) |

#### 2.1 Continuous-time transitions and process covariance

The transition matrix is $\Phi_k = \exp(A \, \Delta t_k)$ with entries derived in the matrix exponential section below. For each branch $b \in \{p, s\}$, the process covariance of $(x_b, I_b)$ over an interval of length $\Delta t_k$ has the block form $\begin{bmatrix} q^b_{xx} & q^b_{xI} \\ q^b_{xI} & q^b_{II} \end{bmatrix}$. Writing $x = \kappa_b \, \Delta t_k$ and using $\sigma^2_b = 2\kappa_b V_b$, the entries are

$$
\begin{aligned}
q^b_{xx} &= V_b(1 - e^{-2x}), \\
q^b_{xI} &= \frac{V_b}{\kappa_b}(1 - e^{-x})^2, \\
q^b_{II} &= \frac{V_b}{\kappa_b^2}(2x - 3 + 4e^{-x} - e^{-2x}).
\end{aligned}
$$

The covariance is block-diagonal across branches because the drives are independent. When $x < 10^{-3}$, the exponential differences suffer catastrophic cancellation, and truncated Taylor expansions in $\Delta t_k$ are used instead:

$$
\begin{aligned}
q^b_{xx} &\approx 2\kappa_b V_b \left[\Delta t_k - \kappa_b \Delta t_k^2 + \frac{2}{3}\kappa_b^2 \Delta t_k^3\right], \\
q^b_{xI} &\approx 2\kappa_b V_b \left[\frac{1}{2}\Delta t_k^2 - \frac{1}{2}\kappa_b \Delta t_k^3 + \frac{7}{24}\kappa_b^2 \Delta t_k^4\right], \\
q^b_{II} &\approx 2\kappa_b V_b \left[\frac{1}{3}\Delta t_k^3 - \frac{1}{4}\kappa_b \Delta t_k^4 + \frac{7}{60}\kappa_b^2 \Delta t_k^5\right].
\end{aligned}
$$

For the vagal branch, $V_p = \lambda_P$ (constant within a recording).

#### 2.2 Dual-filter recursion, Joseph update and phase reset

Two filters are run in parallel and share one covariance recursion and Kalman gain. The stochastic filter $\tilde{X}^{(0)}$ treats the observation as $y_k = 1$ (that is, $\nu_0 = 0$), and the deterministic filter $\tilde{X}^{(\nu)}$ treats it as $y_k = \Delta t_k$ (the unit-$\nu_0$ response). Both start from zero, with initial covariance $P_0 = \mathrm{diag}(\lambda_P, 1, 0, 0)$, which is the stationary covariance of the drives and zero for the phase integrals. At each beat:

1. **Prediction.** $\tilde{X}_{k \mid k-1} = \Phi_k \tilde{X}_{k-1 \mid k-1}$ for both filters, and $P_{k \mid k-1} = \Phi_k P_{k-1 \mid k-1} \Phi_k^\top + Q_k$.
2. **Innovation variance and gain.** With the threshold-noise variance $R_k = \lambda_R\, \mu_k\, \Delta t_k$, where $\mu_k$ is the artifact multiplier defined below, $S_k = H P_{k \mid k-1} H^\top + R_k$ (floored at $10^{-12}$) and $K_k = P_{k \mid k-1} H^\top / S_k$.
3. **Innovations.** $v_k^{(0)} = 1 - H\tilde{X}^{(0)}_{k \mid k-1}$ and $v_k^{(\nu)} = \Delta t_k - H\tilde{X}^{(\nu)}_{k \mid k-1}$.
4. **Update.** $\tilde{X}_{k \mid k} = \tilde{X}_{k \mid k-1} + K_k v_k$ for each filter, and the Joseph form
$$P_{k \mid k} = (I - K_k H)\, P_{k \mid k-1}\, (I - K_k H)^\top + K_k R_k K_k^\top,$$
followed by symmetrization, $P_{k \mid k} \leftarrow \left(P_{k \mid k} + P_{k \mid k}^\top\right)/2$.
5. **Storage and phase reset.** The posterior states and covariance are stored and the integral components ($I_p$, $I_s$) of both state vectors and the corresponding rows and columns of $P_{k \mid k}$ are set to zero, whereas $x_p$, $x_s$ and their covariances are propagated unchanged to the next interval.

The measurement-noise term $R_k$ contains only the threshold-level jitter. The uncertainty in the integrated drives is carried by $HP_{k \mid k-1}H^\top$ and enters $S_k$ but not the Joseph noise term. $R_k$ represents the timing jitter of the depolarization threshold and the timing error of the acquisition system. The Joseph form preserves symmetry and positive semidefiniteness of $P_{k \mid k}$ under finite-precision arithmetic.

**Artifact multiplier.** Let $J_k = \max\big(|\Delta t_k - \Delta t_{k-1}|/\Delta t_{k-1}, |\Delta t_{k+1} - \Delta t_k|/\Delta t_k\big)$ be the largest relative change between adjacent intervals at beat $k$ (one-sided at the ends of the series). The multiplier is $\mu_k = \min\{1 + (J_k/\theta_J)^{p_J}, 10^{10}\}$ with $\theta_J = 0.25$ and $p_J = 10$, and the inflated jitter variance is capped at $10^{12}$. As $\mu_k$ increases, $K_k \to 0$ and the observation has a negligible effect on the states. The multiplier references both adjacent intervals and is therefore not strictly causal; it affects only the observation-noise weighting and not the dynamics of the latent drives. Beats with $\mu_k \geq 100$ are excluded from $\hat{\sigma}^2_{sys}$ and the likelihood.

#### 2.3 GLS profiling and reconstruction of the states

Because $\nu_0$ enters the observation linearly, the generalized least squares (GLS) estimate follows from the two innovation sequences,

$$\hat{\nu}_0 = \frac{\sum_{k=1}^N v_k^{(0)} v_k^{(\nu)} / S_k}{\sum_{k=1}^N \left(v_k^{(\nu)}\right)^2 / S_k},$$

where the artifact-inflated $S_k$ down-weights aberrant beats. With baseline-corrected innovations $v_k = v_k^{(0)} - \hat{\nu}_0 v_k^{(\nu)}$ and $\mathcal{K} = \{k : \mu_k < 100\}$ (all beats if fewer than two satisfy this), the concentrated estimate of the global scale and the profiled log-likelihood are

$$\hat{\sigma}^2_{sys} = \frac{1}{n_{\text{eff}}} \sum_{k \in \mathcal{K}} \frac{v_k^2}{S_k}, \qquad \mathcal{L}^* = -\frac{1}{2} \sum_{k \in \mathcal{K}} \ln S_k - \frac{n_{\text{eff}}}{2} \ln \hat{\sigma}^2_{sys}, \qquad n_{\text{eff}} = |\mathcal{K}|.$$

If $\hat{\nu}_0$ is not finite and positive, the implied RR would be undefined, and the fit is assigned $\mathcal{L}^* = -\infty$.

The reconstructed (forward-filtered) states are $\hat{X}_{k \mid k} = \tilde{X}^{(0)}_{k \mid k} - \hat{\nu}_0\, \tilde{X}^{(\nu)}_{k \mid k}$. The variances of the drives include the uncertainty in $\hat{\nu}_0$, whose variance is $\hat{\sigma}^2_{sys}/D$ with $D = \sum_k \left(v_k^{(\nu)}\right)^2 / S_k$:

$$\mathrm{Var}\big(\hat{x}_{b,k}\big) = \hat{\sigma}^2_{sys} \left[ P^{(bb)}_{k \mid k} + \frac{\left(\tilde{X}^{(\nu)}_{b,k \mid k}\right)^2}{D} \right], \quad b \in \{p, s\}.$$

The filtered RR at beat $k$ is $\widehat{RR}_k = \left(1 + \hat{I}_{p,k} - \hat{I}_{s,k}\right)/\hat{\nu}_0$ with variance $\hat{\sigma}^2_{sys}\left(P^{(33)}_{k\mid k} + P^{(44)}_{k\mid k} - 2P^{(34)}_{k\mid k}\right)/\hat{\nu}_0^2$. Because these quantities use the update at beat $k$, they are in-sample filtered values. The stated 95% intervals are conditional on the MAP kinetic parameters.

#### 2.4 Spectral anchoring

The relative vagal variance $\lambda_P = V_p/V_s$ is computed once per recording as follows.

1. **Resampling.** The instantaneous heart rate (Hz), $1/\Delta t_k$, is interpolated onto a uniform 4 Hz grid with a monotone (Fritsch–Carlson) Hermite spline, which avoids overshoot between beats, and the mean is subtracted.
2. **Global wavelet spectrum.** A Morlet continuous wavelet transform is computed [R package `biwavelet`]. Power at each scale is masked outside the cone of influence and averaged over time to give the global spectrum $G(f)$, and frequencies are sorted in ascending order.
3. **Frequency weights.** A logistic transition $L(f) = [1 + \exp\{-s\,(f - f_c)\}]^{-1}$ with crossover $f_c = 0.10$ Hz defines the weights
$$w_v(f) = \phi + (1-\phi)\,L(f), \qquad w_s(f) = (1-\phi)\,(1 - L(f)),$$
with $w_v + w_s = 1$, where $\phi = 0.25$ is the floor fraction of the vagal weight at low frequencies. The steepness $s = -\ln\{\epsilon/(1-\epsilon)\}/(f_u - f_c)$ is set so that $1 - L(f_u) = \epsilon = 0.05$ at the LF/HF boundary $f_u = 0.15$ Hz, which defines the "sympathetic leakage" into the high-frequency band.
4. **Energy ratio.** Energies are integrated by the trapezoidal rule over the log-spaced scales on $0.04 \leq f \leq 0.40$ Hz,
$$E_v = \int w_v(f)\,G(f)\,df, \qquad E_s = \int w_s(f)\,G(f)\,df, \qquad \lambda_P = E_v/E_s.$$

The unweighted low- and high-frequency band powers (LF: 0.04–0.15 Hz; HF: 0.15–0.40 Hz) and the mean heart rate are also stored as diagnostics and are not used in the fit.

The anchor rests on two assumptions that merit explicit statement. First, the spectral weights encode external evidence about the distribution of vagal and sympathetic modulation across frequency. Pharmacological blockade paradigms indicate that parasympathetic tone forms a persistent baseline contribution to resting LF power [1,2]; because the size of this contribution varies across individuals and conditions, the floor fraction is set to a deliberately conservative $\phi$ = 0.25 (a quarter of the low-frequency weight is attributed to the vagal branch) and is varied between 0.10 and 0.50 in the sensitivity analyses. In addition, sympathetic oscillations peak heavily around the 0.10 Hz baroreflex resonance frequency ($f_c$ = 0.10 Hz) [2]. Furthermore, due to the sluggish intracellular signaling kinetics of beta-adrenergic cascades, sympathetic transfer is structurally limited to a negligible leakage fraction of less than 5% ($\epsilon$ = 0.05) above the standard 0.15 Hz high-frequency boundary [3–5]. Second, the OU drives have Lorentzian power spectra whose integral equals the stationary variance (Parseval relation), so that the ratio of partitioned energies in the observed heart-rate spectrum is taken as the ratio of latent stationary variances. This ignores differences in the transfer from latent drive to observed heart rate (for example, the low-pass effect of integration over each interval) and is therefore an approximation.

#### 2.5 Priors, MAP objective and optimization

The parameters $\theta = (\theta_1, \theta_2, \theta_3)$ are mapped to the physical scale by $\kappa_s = e^{\theta_1}$, $\kappa_p = \kappa_s + e^{\theta_2}$ and $\lambda_R = e^{\theta_3}$, which ensures positivity and $\kappa_p > \kappa_s$ without bound constraints. The log-prior terms, defined up to additive constants, are as follows.

*Time-scale priors.* For $b \in \{p, s\}$ with default central $c$-coverage interval $\left[\tau_b^{lo}, \tau_b^{hi}\right]$ (default $c = 0.95$),
$$\ln p(\tau_b) = -\frac{1}{2}\left(\frac{\ln \tau_b - \mu_b}{\varsigma_b}\right)^2, \quad \mu_b = \frac{1}{2}\left(\ln \tau_b^{lo} + \ln \tau_b^{hi}\right), \quad \varsigma_b = \frac{\ln \tau_b^{hi} - \ln \tau_b^{lo}}{2\, z_{(1+c)/2}},$$

that is, a log-normal prior whose central interval is the stated range, with $z_{(1+c)/2}$ the standard normal quantile.

*Separation prior.* With $U = \left(\kappa_p - \kappa_s\right)/\left(\kappa_p + \kappa_s\right) \in (0,1)$ and $1 - U = 2\kappa_s/\left(\kappa_p + \kappa_s\right)$,

$$\ln p(U) = (\alpha - 1)\ln U + (\beta - 1)\ln(1 - U),$$
$$\alpha - 1 = w\,U^*, \quad \beta - 1 = w\,(1 - U^*), \quad U^* = \frac{r-1}{r+1},$$

which is a Beta density with mode $U^*$ corresponding to $\kappa_p/\kappa_s = r$ and strength $w$. The defaults are $r = 20$ and $w = 10$.

*Timing-jitter prior.* The equivalent RR timing jitter at the median interval $\widetilde{\Delta t}$ is $s_{RR} = \sqrt{\hat{\sigma}^2_{sys}\,\lambda_R\,\widetilde{\Delta t}/\hat{\nu}_0}$, and a log-normal prior is placed on it, constructed as above from the central interval $[0.5, 20]$ ms. The hyperparameters are fixed and independent of the data, but $s_{RR}$ depends on the profiled quantities.

The objective minimized is

$$\mathrm{Cost}(\theta) = -\left[\mathcal{L}^* + \ln p(\tau_s) + \ln p(\tau_p) + \ln p(U) + \ln p(s_{RR})\right],$$

and non-finite values are replaced by $10^{12}$ so that the optimizer backs off. Optimization uses the BFGS algorithm (R `optim`, finite-difference gradients, at most 1000 iterations) in coordinates scaled by the spread of the priors, $z_1 = (\ln\tau_p - \mu_p)/s_p$, $z_2 = (\ln d - \ln d_0)/s_d$ with $d = \ln(\tau_s/\tau_p) = \ln(\kappa_p/\kappa_s) > 0$, $d_0 = \mu_s - \mu_p$, $s_d = \sqrt{s_p^2 + s_s^2}/d_0$, and $z_3 = \ln\lambda_R$ ($\mu$ and $s$ are the parameters of the log-normal priors). The map keeps $\kappa_p > \kappa_s$ and changes only the geometry seen by the optimizer, not the objective. Three starts are run: (i) the default, with $\kappa_s$ and $\kappa_p$ at the inverse of the geometric centers of the time-scale priors (with $\kappa_p - \kappa_s$ floored at 0.05 s $^{-1}$) and $\lambda_{R,0} = 10^{-3}$; (ii) a noise-free start ($\tau_p$ = 12 s, $\tau_s$ = 160 s, $\lambda_R = 10^{-6}$); and (iii) a noisy start ($\tau_p$ = 25 s, $\tau_s$ = 150 s, $\lambda_R = 0.3$). The last two cover the two ends of the noise-to-signal ratio, which spans several orders of magnitude and where narrow secondary optima were found in preliminary checks with a single start. The run with the lowest cost is kept, and among runs within 0.01 of the lowest the earliest, because runs that reach the same optimum differ by the optimizer tolerance. The default start reached the lowest objective (within 0.01) in 306 of 313 recordings; in the other 7 (2%), 4 from the noise-free start and 3 from the noisy start, the lowest objective was below that of the default start by a median of 46.6 (range 10.2 to 662.7). Three starts do not guarantee a global optimum.

#### 2.6 Parameter uncertainty

Standard errors are computed from the numerical Hessian $\mathcal{H}$ of $\mathrm{Cost}(\theta)$ at the optimum. The covariance of $\theta$ is approximated by the Moore–Penrose pseudo-inverse $\mathcal{H}^+$, which tolerates near-singularity, and mapped to the physical parameters through the Jacobian $\mathcal{J}$ of $\theta \mapsto (\kappa_s, \kappa_p, \lambda_R)$, giving $\mathrm{Cov} = \mathcal{J}\,\mathcal{H}^+\,\mathcal{J}^\top$. For the time constants, the delta method gives $\mathrm{Var}(\tau) \approx \mathrm{Var}(\kappa)/\kappa^4$. Wald 95% intervals are estimate $\pm$ 1.96 SE. The profiled ($\hat{\nu}_0$, $\hat{\sigma}^2_{sys}$), fixed ($\lambda_P$, hyperparameters) and derived quantities carry no standard error, and the intervals therefore do not reflect uncertainty in $\lambda_P$ or in the hyperparameters.

#### 2.7 Default settings

Table S 2 lists the default configuration of the engine.

Table S 2: Default configuration of the estimation engine

| Group | Setting | Default |
|---|---|---|
| Time-scale priors | Central 95% interval of $\tau_p$ / $\tau_s$ | [3, 6] s / [100, 300] s |
| Separation prior | Target $\kappa_p/\kappa_s$; weight | 20; 10 |
| Timing-jitter prior | Central 95% interval | [0.5, 20] ms |
| Spectral anchor | Integration range; crossover $f_c$; leakage $\epsilon$ at 0.15 Hz; vagal floor $\phi$; CWT sampling rate | 0.04–0.40 Hz; 0.10 Hz; 0.05; 0.25; 4 Hz |
| Artifact multiplier | $\theta_J$; $p_J$; likelihood-exclusion multiplier | 0.25; 10; 100 |
| Optimizer | Method; maximum iterations; starts ($\tau_p$, $\tau_s$, $\lambda_R$); floor on $\kappa_p - \kappa_s$ at the default start | BFGS; 1000; prior centers and $10^{-3}$, 12 s, 160 s and $10^{-6}$, 25 s, 150 s and 0.3; 0.05 s$^{-1}$ |
| Reporting | Diagnostic multiplier limit; PACF lags | 1.5; 40 |

All reported analyses use these defaults unless stated in the main text; the configuration file (`config.R`) accompanying the analysis code records every setting, including the analysis windows, the number of multi-start fits and the simulation sizes.

### 3 Goodness-of-fit diagnostic in diffusion-based phase-domain architectures

Point-process models are commonly assessed with the Kolmogorov–Smirnov (KS) test applied to time-rescaled intervals. The classical time-rescaling theorem assumes probabilistic event arrivals, whereas SPARK models a deterministic threshold crossing of a stochastic diffusion. This section describes how the same principle, which is the probability integral transform of model-standardized quantities to a uniform distribution, is applied to the phase-domain innovations. The two settings differ in the generative noise distribution but share this mapping.

#### 3.1 The classical Poisson arrival formulation

In classical point-process statistics, a heartbeat is a stochastic arrival governed by a conditional intensity function $\lambda(t)$. The time-rescaling theorem states that integrating the intensity over the observed inter-beat interval $[t_{k-1}, t_k]$,

$$\Lambda_k = \int_{t_{k-1}}^{t_k} \lambda(t)\, dt,$$

transforms the sequence of intervals into unit-rate exponential variates, $\Lambda_k \sim \text{Exp}\ (1)$. The exponential CDF maps these to a uniform distribution:

$$u_k = 1 - \exp(-\Lambda_k) \sim \mathcal{U}(0, 1),$$

which is evaluated with the KS test.

#### 3.2 The phase-domain diffusion formulation

In SPARK, the event is triggered when the accumulated phase reaches a threshold of 1, and the stochasticity arises from the diffusion within the interval together with uncertainty about the latent drives. Over the observed interval, the filter predicts the phase from the past data and the baseline, and the innovation is

$$v_k = v_k^{(0)} - \hat{\nu}_0\, v_k^{(\nu)} = 1 - \hat{\nu}_0\, \Delta t_k - H\, \hat{X}_{k \mid k-1},$$

the difference between the observed unit phase and its one-step-ahead prediction. If the model and the parameters are correct, $v_k$ is conditionally Gaussian with zero mean and variance $\sigma^2_{sys} S_k$, where $S_k = H P_{k \mid k-1} H^\top + R_k$ combines the prediction uncertainty of the integrated drives and the threshold noise. In contrast to the exponential variates of the Poisson formulation, the standardized innovation is therefore normal.

#### 3.3 Standardization and the probability integral transform

The standardized innovation is

$$z_k = \frac{v_k}{\sqrt{\hat{\sigma}^2_{sys}\, S_k}} \approx \mathcal{N}(0, 1),$$

and the probability integral transform with the standard normal CDF gives

$$u_k = \Phi(z_k) \approx \mathcal{U}(0, 1).$$

Uniformity of $u_k$ is assessed with the KS distance $D$, and the approximate 5% critical value is $1.36/\sqrt{N}$. Because $\hat{\nu}_0$, $\hat{\sigma}^2_{sys}$ and the kinetic parameters are estimated from the same series, the nominal critical value is approximate, and the test is conservative in this respect. Serial independence is assessed with the partial autocorrelation function (PACF) of $z_k$ over 40 lags, with bounds at $\pm 1.96/\sqrt{N}$. Diagnostics retain beats with finite innovations and artifact multiplier $\mu_k < 1.5$, and the innovation variance used for standardization excludes the artifact inflation. The plug-in values used for $\hat{\nu}_0$ and $\hat{\sigma}^2_{sys}$ make the standardized innovations approximate.

The diagnostic is the probability integral transform of the model-standardized innovations: $u_k = \Phi(z_k)$ with $\Phi$ the standard normal distribution function, and not $1 - e^{-\Lambda_k}$ as in the time-rescaling theorem. It shares with time-rescaling the idea of isolating the model-standardized noise and mapping it to a uniform distribution, and it is not an application of the Poisson-specific theorem. Agreement with uniformity and serial independence is necessary for adequacy but does not establish it, since the diagnostics may have limited power against specific misspecifications, including the structure of the latent decomposition.

### 4 Interval-variance scaling under first-passage

Respiratory sinus arrhythmia is generally larger when vagal tone is high, so that the variance of the observed intervals depends on the vagal state. This section examines how such dependence arises under a strict first-passage formulation, and why it is not carried automatically by the phase-domain filter.

For a Brownian motion with constant drift $\nu$, diffusion coefficient $\sigma_z$ and threshold $a = 1$, the first-passage time $\tau$ follows an inverse Gaussian distribution with

$$\mathbb{E}[\tau] = \frac{1}{\nu}, \qquad \mathrm{Var}(\tau) = \frac{\sigma_z^2}{\nu^3}.$$

The variance is inversely proportional to the cube of the drift. For a fixed diffusion coefficient, a decrease in the net drift, such as that produced by greater vagal influence, therefore increases the variance of the interval by a power law (and not exponentially), whereas an increase in the drift decreases it. This is a qualitative agreement with the dependence of interval variability on vagal tone, and it holds for the strict first-passage formulation with locally constant drift.

In the phase-domain filter, the measurement equation conditions on the observed interval, and its variance, $\sigma_{sys}^2 S_k$, depends on the kinetic parameters, the time elapsed and the artifact multiplier, but not on the level of $x_p$. State-dependent interval variance is therefore not represented by the filter, and the stationary variances of the two drives are treated as constant within a recording. This is an assumption that may be restrictive when the vagal level changes strongly, for example during exercise or orthostatic perturbation (see Limitations in the main text).

### 5 Continuous-discrete state-space formulation via matrix exponential

The framework evaluates continuous-time kinetics across irregular sampling intervals without Euler–Maruyama approximation, by augmenting the state with the running integrals of the drives. Let $\tilde{X}(t) = \left[x_p(t), x_s(t), I_p(t), I_s(t)\right]^\top$, where $x_p$ and $x_s$ are the Ornstein–Uhlenbeck drives and $I_p$, $I_s$ their running integrals. The dynamics are the linear SDE

$$d\tilde{X}(t) = A\,\tilde{X}(t)\,dt + G\,dW(t),$$

with

$$A = \begin{bmatrix} -\kappa_p & 0 & 0 & 0 \\ 0 & -\kappa_s & 0 & 0 \\ 1 & 0 & 0 & 0 \\ 0 & 1 & 0 & 0 \end{bmatrix}, \qquad G = \begin{bmatrix} \sigma_p & 0 \\ 0 & \sigma_s \\ 0 & 0 \\ 0 & 0 \end{bmatrix}.$$

Because the system is linear and time-invariant between observations, the transition over any observed interval $\Delta t_k$ is given by the matrix exponential:

$$\Phi_k = \exp(A\,\Delta t_k) = \begin{bmatrix} e^{-\kappa_p \Delta t_k} & 0 & 0 & 0 \\ 0 & e^{-\kappa_s \Delta t_k} & 0 & 0 \\ \frac{1-e^{-\kappa_p \Delta t_k}}{\kappa_p} & 0 & 1 & 0 \\ 0 & \frac{1-e^{-\kappa_s \Delta t_k}}{\kappa_s} & 0 & 1 \end{bmatrix}.$$

This transition allows the Kalman recursion to propagate the state across irregular intervals without discretization error in the deterministic part of the dynamics; the process covariance is computed from the closed-form integrals given in the estimation algorithm. After each measurement update, the integral states are reset to zero, which represents the restart of phase accumulation at each beat.

#### 5.1 Explicit derivation of the transition matrix

The matrix exponential is evaluated from its Taylor series,

$$\exp(A\,\Delta t_k) = \sum_{n=0}^{\infty} \frac{(A\,\Delta t_k)^n}{n!} = I + A\,\Delta t_k + \frac{1}{2!} A^2 \Delta t_k^2 + \dots$$

Given the block structure of $A$, its powers follow a regular pattern. For any integer $n \geq 1$,

$$A^n = \begin{bmatrix} (-\kappa_p)^n & 0 & 0 & 0 \\ 0 & (-\kappa_s)^n & 0 & 0 \\ (-\kappa_p)^{n-1} & 0 & 0 & 0 \\ 0 & (-\kappa_s)^{n-1} & 0 & 0 \end{bmatrix}.$$

For the top-left element, which governs $x_p$, the series reduces to the scalar exponential:

$$\sum_{n=0}^{\infty} \frac{(-\kappa_p \Delta t_k)^n}{n!} = e^{-\kappa_p \Delta t_k}.$$

For the element in position $(3,1)$, which governs $I_p$, the series starts at $n = 1$, because the identity matrix has a zero in this position. Extracting a factor of $-1/\kappa_p$ gives

$$\sum_{n=1}^{\infty} \frac{\Delta t_k^n}{n!} (-\kappa_p)^{n-1} = -\frac{1}{\kappa_p} \sum_{n=1}^{\infty} \frac{(-\kappa_p \Delta t_k)^n}{n!}.$$

The right-hand sum is the Taylor series of $e^{-\kappa_p \Delta t_k}$ minus its $n = 0$ term, which equals 1, so that

$$-\frac{1}{\kappa_p}\left(e^{-\kappa_p \Delta t_k} - 1\right) = \frac{1 - e^{-\kappa_p \Delta t_k}}{\kappa_p}.$$

Because the vagal and sympathetic pathways are decoupled in $A$, the same reduction applies to the sympathetic parameters. The element in position $(2,2)$ is

$$\sum_{n=0}^{\infty} \frac{(-\kappa_s \Delta t_k)^n}{n!} = e^{-\kappa_s \Delta t_k},$$

and the element in position $(4,2)$ is

$$-\frac{1}{\kappa_s}\left(e^{-\kappa_s \Delta t_k} - 1\right) = \frac{1 - e^{-\kappa_s \Delta t_k}}{\kappa_s}.$$

Finally, the right half of $A^n$ is zero for all $n \geq 1$, so the elements in positions $(3,3)$ and $(4,4)$ come only from the identity matrix and equal 1. Assembling these scalar solutions gives the closed-form transition matrix $\Phi_k$, in which the accumulation of the integral states is evaluated analytically and without numerical integration over each interval.

### 6 Simulation model and recovery under known truth

Synthetic recordings were generated from the model so that the latent drives are known. On a fine grid of step $h = 0.01$ s, the drives were simulated by their exact Ornstein–Uhlenbeck transition, $x(t+h) = e^{-\kappa h}x(t) + \sqrt{V(1-e^{-2\kappa h})}\,\varepsilon$, with stationary variances $V_s = \sigma^2_{sys}$ and $V_p = \lambda_P \sigma^2_{sys}$; the phase increments were $\{\nu_0 - x_p + x_s\}\,h + \sqrt{\sigma^2_{sys}\lambda_R h}\,\varepsilon'$, a beat was registered at the interpolated threshold crossing, and the phase was reset at each beat. Deterministic profiles added to the drives (exponential approach to a target amplitude, followed by exponential decay for exercise) were used to mimic a perturbation or an exercise bout; these departures from the stationary OU model make the benchmarks conservative with respect to model misspecification. Default generating values were $\nu_0 \approx 1.2$ Hz, $\sigma^2_{sys} = 9 \times 10^{-4}$ Hz$^2$ and $\lambda_R = 0.02$.

The recovery study used four scenarios: true time scales within the prior ranges ($\tau_p$ 3.5–5.5 s, $\tau_s$ 120–250 s), outside them ($\tau_p$ 8–12 s, $\tau_s$ 40–70 s), and with vagal-dominant ($\lambda_P$ 6–12) or sympathetic-dominant ($\lambda_P$ 0.2–0.5) variance. Each specification was fitted to each recording and compared with the generating values (supplementary figure S1).

Table S 3: Recovery under known truth: median absolute log-error of the time constants, median correlation between reconstructed and true states, median absolute error of the share of vagal variance, and share of time points at which the 95% interval of the filtered state contained the true state (conditional on the MAP parameters), by scenario and specification.

| Scenario | Spec. | n | Abs. error of ln $\tau_p$ | Abs. error of ln $\tau_s$ | Corr. of $x_p$ | Corr. of $x_s$ | Abs. error of the share of $x_p$ variance | Coverage of $x_p$ | Coverage of $x_s$ |
|---|---|---|---|---|---|---|---|---|---|
| True time scales within the priors | $M_0$ | 100 | 0.18 | 1.40 | 0.88 | 0.53 | 0.09 | 88% | 58% |
| True time scales within the priors | $M_1$ | 100 | 0.08 | 0.20 | 0.88 | 0.54 | 0.08 | 90% | 51% |
| True time scales within the priors | $M_2$ | 100 | 0.17 | 2.21 | 0.87 | 0.52 | 0.11 | 94% | 94% |
| True time scales within the priors | $M_3$ | 100 | 0.08 | 0.19 | 0.87 | 0.57 | 0.07 | 96% | 97% |
| True time scales outside the priors | $M_0$ | 100 | 0.32 | 1.32 | 0.85 | 0.50 | 0.65 | 68% | 71% |
| True time scales outside the priors | $M_1$ | 100 | 0.70 | 1.01 | 0.76 | 0.53 | 0.42 | 78% | 79% |
| True time scales outside the priors | $M_2$ | 100 | 0.30 | 1.36 | 0.86 | 0.50 | 0.18 | 92% | 94% |
| True time scales outside the priors | $M_3$ | 100 | 0.55 | 0.98 | 0.84 | 0.53 | 0.08 | 89% | 95% |
| Sympathetic-dominant variance | $M_0$ | 100 | 2.65 | 2.33 | 0.46 | 0.80 | 0.33 | 83% | 64% |
| Sympathetic-dominant variance | $M_1$ | 100 | 0.11 | 0.22 | 0.71 | 0.85 | 0.14 | 91% | 81% |
| Sympathetic-dominant variance | $M_2$ | 100 | 0.93 | 2.22 | 0.60 | 0.80 | 0.25 | 96% | 63% |
| Sympathetic-dominant variance | $M_3$ | 100 | 0.09 | 0.28 | 0.71 | 0.81 | 0.15 | 90% | 70% |
| Vagal-dominant variance | $M_0$ | 100 | 0.17 | 1.34 | 0.92 | 0.37 | 0.05 | 90% | 49% |
| Vagal-dominant variance | $M_1$ | 100 | 0.09 | 0.19 | 0.92 | 0.31 | 0.04 | 91% | 30% |
| Vagal-dominant variance | $M_2$ | 100 | 0.19 | 3.17 | 0.91 | 0.32 | 0.07 | 94% | 99% |
| Vagal-dominant variance | $M_3$ | 100 | 0.08 | 0.19 | 0.90 | 0.41 | 0.11 | 97% | 100% |

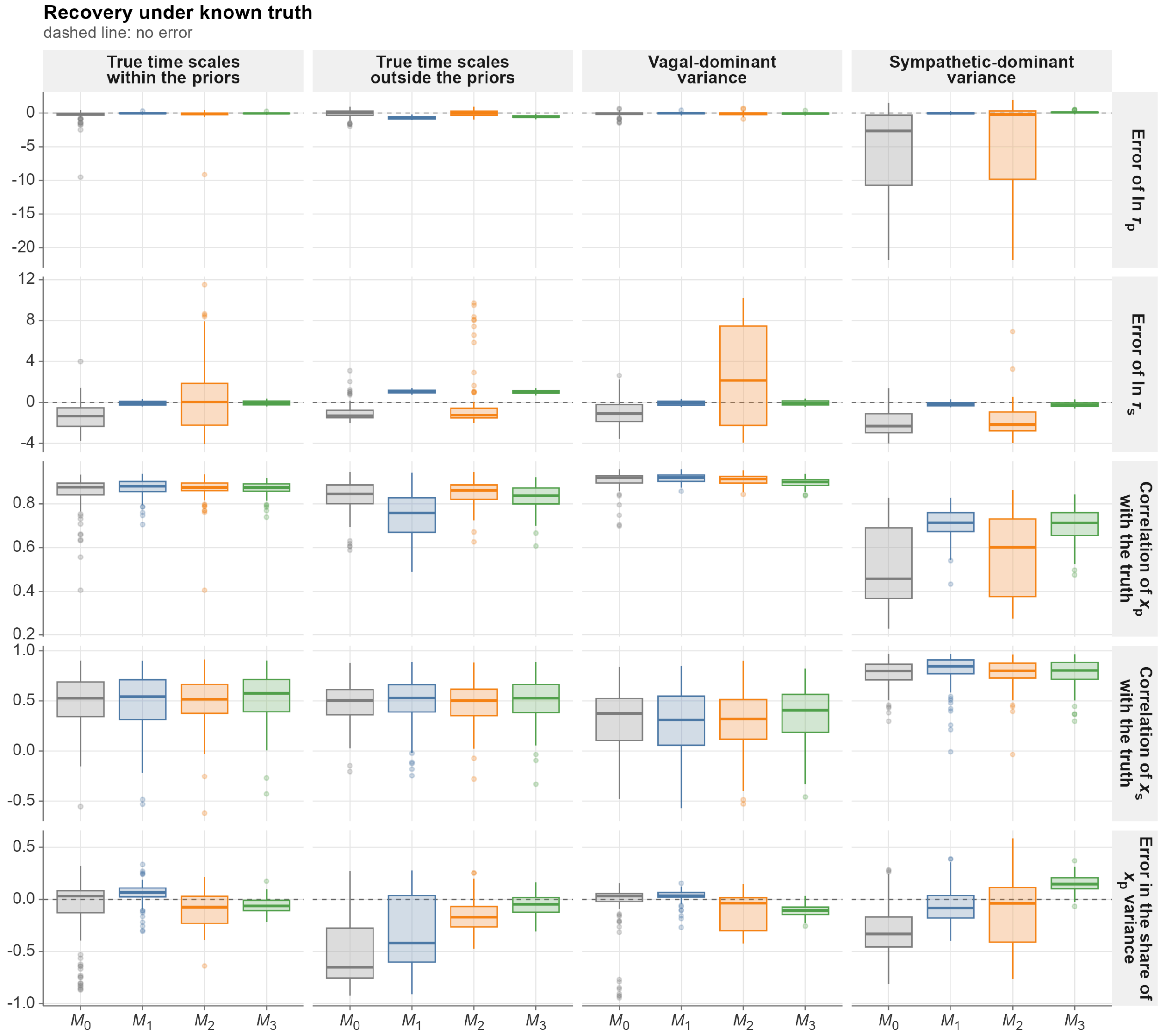


Figure S 1: Recovery under known truth. Errors in the time constants and in the share of vagal variance, and correlation of the reconstructed states with the true drives, by scenario and specification.

### 7 Identifiability and sensitivity analyses: details

*Starting points.* The first start of each recording is the engine default (centers of the time-scale priors, $\lambda_{R,0} = 10^{-3}$). The remaining starts are drawn log-uniformly with the same random numbers for all specifications of a recording, seeded by the recording identifier. The vagal rate is required to exceed the sympathetic rate by construction ($\kappa_p = \kappa_s + e^{\theta_2}$).

*Distinct solutions.* Starts are visited in order, and a start opens a new solution if its distance in $\left(\ln \tau_p, \ln \tau_s\right)$ to every existing solution center exceeds the clustering radius (0.10, approximately a 10% difference in each time constant).

*Metrics of the multi-start fits.* Starts are drawn log-uniformly ($\tau_s$ 20–600 s; $\tau_p$ 0.5–30 s with $\tau_p < 0.8\,\tau_s$; $\lambda_R$ $10^{-4}$ –$10^{-1}$; $\lambda_P$ 0.05–20 for specifications in which it is free). For each recording and specification, the following quantities were computed: (i) the number of *distinct solutions*, defined by greedy clustering of the estimates in $\left(\ln \tau_p, \ln \tau_s\right)$ with radius 0.10; (ii) the *near-equivalent fits*, defined as fits whose profiled log-likelihood lies within 2 units of the best fit, together with the dispersion of $\ln \tau_p$ and $\ln \tau_s$ and the number of distinct solutions among them; (iii) the dispersion of the reconstructed trajectories among near-equivalent fits (root-mean-square across-start SD of $x_p(t)$ and $x_s(t)$ relative to their within-trajectory SD) and their median pairwise correlation; (iv) the *share of $x_p$ variance*, $\mathrm{Var}(\hat{x}_p)/\left[\mathrm{Var}(\hat{x}_p) + \mathrm{Var}(\hat{x}_s)\right]$, and the proportion of fits in which one process absorbed essentially all variability (share below 0.05 or above 0.95), referred to as one-process collapse; (v) the proportion of fits whose $\tau_p$ and $\tau_s$ both lie within the 95% intervals of the time-scale priors; and (vi) the retained fit to RR (RMSE and profiled log-likelihood). Because the priors alter the objective, specifications are compared on the profiled log-likelihood and RR error, and not on the posterior objective.

*Quantities recorded in the sensitivity analyses.* Using $M_3$ as the reference, each hyperparameter of the identification structure was varied one at a time (time-scale intervals, separation target and weight, timing-jitter interval, spectral crossover, sympathetic leakage and vagal floor), and each prior component was removed in turn (leave-one-out). For each variation, the log-ratio of $\tau_p$ and $\tau_s$ to the reference, the correlation and the normalized root-mean-square difference of the reconstructed states, and the change in RR error were recorded. The data-informativeness of the kinetic estimates was assessed by the position of the estimates relative to the prior (standardized distance from the prior center and the proportion within the 95% prior interval), by the posterior contraction of $\ln \tau$ (one minus the ratio of posterior to prior variance, from the Laplace approximation), and by the log-ratio of the $M_3$ and $M_0$ estimates.

*Laplace standard deviations.* With $\mathcal{H}$ the numerical Hessian of the objective in $\theta = (\ln \kappa_s, \ln(\kappa_p - \kappa_s), \ldots)$ and $\mathcal{H}^+$ its pseudo-inverse, the covariance of $(\ln \tau_s, \ln \tau_p)$ is $J\mathcal{H}^+J^\top$ with $\partial \ln \tau_s / \partial \theta_1 = -1$, $\partial \ln \tau_p / \partial \theta_1 = -\kappa_s/\kappa_p$ and $\partial \ln \tau_p / \partial \theta_2 = -(\kappa_p - \kappa_s)/\kappa_p$. Posterior contraction relative to the log-normal prior is $1 - \mathrm{Var}_{post}/\mathrm{Var}_{prior}$, with $\mathrm{Var}_{prior} = \varsigma^2$ as defined in the estimation algorithm. A negative value indicates a posterior wider than the prior; the quantity is local and approximate (Laplace) and is used descriptively.

*Profile surfaces.* On a log-spaced grid ($\tau_p$ 0.5–30 s, $\tau_s$ 20–600 s), $\tau_p$ and $\tau_s$ were fixed at each grid point with $\tau_p < \tau_s$, and $\lambda_R$ (and $\lambda_P$ in $M_0$, $M_1$) were optimized (one-dimensional Brent search or Nelder–Mead). The surface is the resulting objective minus its minimum.

*Hyperparameter variations.* Table S 4 lists the variations. Each was applied to $M_3$ with all other settings at their defaults, and the leave-one-out specifications remove the time-scale priors, the separation prior, the timing-jitter prior or the spectral anchor (in which case $\lambda_P$ becomes free).

Table S 4: Hyperparameter variations used in the sensitivity analysis

| Hyperparameter | Default | Variations |
|---|---|---|
| $\tau_p$ interval (s) | [3, 6] | [3.5, 5], [2, 9], [1.5, 3], [6, 12] |
| $\tau_s$ interval (s) | [100, 300] | [150, 250], [60, 500], [50, 150], [200, 600] |
| Separation target $\kappa_p/\kappa_s$ | 20 | 10, 40 |
| Separation weight | 10 | 5, 20 |
| Timing-jitter interval (ms) | [0.5, 20] | [1, 10], [0.2, 50] |
| Spectral crossover (Hz) | 0.10 | 0.08, 0.12 |
| Sympathetic leakage at 0.15 Hz | 0.05 | 0.02, 0.10 |
| Vagal floor fraction | 0.25 | 0.1, 0.5 |

Table S 5: Sensitivity of the $M_3$ reconstructions to the hyperparameters (median across recordings; the last column is the absolute change in the share of latent variance attributed to the vagal-like process). Varied: one hyperparameter changed at a time; Removed: one prior term left out.

| Kind | Setting | n | Abs. change in ln $\tau_p$ | Abs. change in ln $\tau_s$ | Corr. of $x_p$ | Corr. of $x_s$ | Rel. RMSD of $x_p$ | Rel. RMSD of $x_s$ | Abs. change in the share of $x_p$ variance |
|---|---|---|---|---|---|---|---|---|---|
| Varied | Spectral crossover (Hz): 0.08 | 150 | 0.13 | 0.05 | 0.996 | 0.996 | 0.22 | 0.21 | 0.17 |
| Varied | Spectral crossover (Hz): 0.12 | 150 | 0.11 | 0.04 | 0.997 | 0.998 | 0.15 | 0.16 | 0.10 |
| Varied | Timing-jitter interval: narrow | 150 | 0.00 | 0.00 | 1.000 | 1.000 | 0.00 | 0.00 | 0.00 |
| Varied | Timing-jitter interval: wide | 150 | 0.00 | 0.00 | 1.000 | 1.000 | 0.00 | 0.00 | 0.00 |
| Varied | Vagal floor fraction: 0.1 | 150 | 0.12 | 0.03 | 0.997 | 0.998 | 0.16 | 0.16 | 0.11 |
| Varied | Vagal floor fraction: 0.5 | 150 | 0.18 | 0.07 | 0.992 | 0.993 | 0.29 | 0.29 | 0.23 |
| Varied | Sympathetic leakage: 0.02 | 150 | 0.01 | 0.00 | 1.000 | 1.000 | 0.02 | 0.02 | 0.02 |
| Varied | Sympathetic leakage: 0.10 | 150 | 0.02 | 0.01 | 1.000 | 1.000 | 0.03 | 0.03 | 0.02 |
| Varied | $\tau_p$ interval: high | 150 | 0.31 | 0.09 | 0.998 | 0.999 | 0.08 | 0.09 | 0.05 |
| Varied | $\tau_p$ interval: low | 150 | 0.27 | 0.09 | 0.999 | 0.999 | 0.07 | 0.08 | 0.04 |
| Varied | $\tau_p$ interval: narrow | 150 | 0.53 | 0.18 | 0.995 | 0.997 | 0.17 | 0.14 | 0.09 |
| Varied | $\tau_p$ interval: wide | 150 | 0.64 | 0.19 | 0.994 | 0.997 | 0.17 | 0.16 | 0.10 |
| Varied | $\tau_s$ interval: high | 150 | 0.04 | 0.35 | 0.999 | 0.997 | 0.07 | 0.09 | 0.03 |
| Varied | $\tau_s$ interval: low | 150 | 0.03 | 0.27 | 0.999 | 0.998 | 0.06 | 0.06 | 0.01 |
| Varied | $\tau_s$ interval: narrow | 150 | 0.02 | 0.18 | 1.000 | 0.999 | 0.03 | 0.05 | 0.01 |
| Varied | $\tau_s$ interval: wide | 150 | 0.01 | 0.09 | 1.000 | 1.000 | 0.02 | 0.02 | 0.00 |
| Varied | Separation weight: 20 | 150 | 0.01 | 0.03 | 1.000 | 1.000 | 0.00 | 0.01 | 0.00 |
| Varied | Separation weight: 5 | 150 | 0.00 | 0.02 | 1.000 | 1.000 | 0.00 | 0.00 | 0.00 |
| Varied | Separation target $\kappa_p/\kappa_s$: 10 | 150 | 0.01 | 0.03 | 1.000 | 1.000 | 0.01 | 0.01 | 0.00 |
| Varied | Separation target $\kappa_p/\kappa_s$: 40 | 150 | 0.00 | 0.01 | 1.000 | 1.000 | 0.00 | 0.00 | 0.00 |
| Removed | $M_3$ without the spectral anchor | 150 | 1.17 | 0.20 | 0.598 | 0.939 | 0.94 | 0.88 | 0.37 |
| Removed | $M_3$ without the timing-jitter prior | 150 | 0.00 | 0.00 | 1.000 | 1.000 | 0.00 | 0.00 | 0.00 |
| Removed | $M_3$ without the timescale priors | 150 | 2.08 | 2.40 | 0.950 | 0.932 | 0.68 | 0.65 | 0.40 |
| Removed | $M_3$ without the separation prior | 150 | 0.01 | 0.04 | 1.000 | 1.000 | 0.01 | 0.01 | 0.00 |

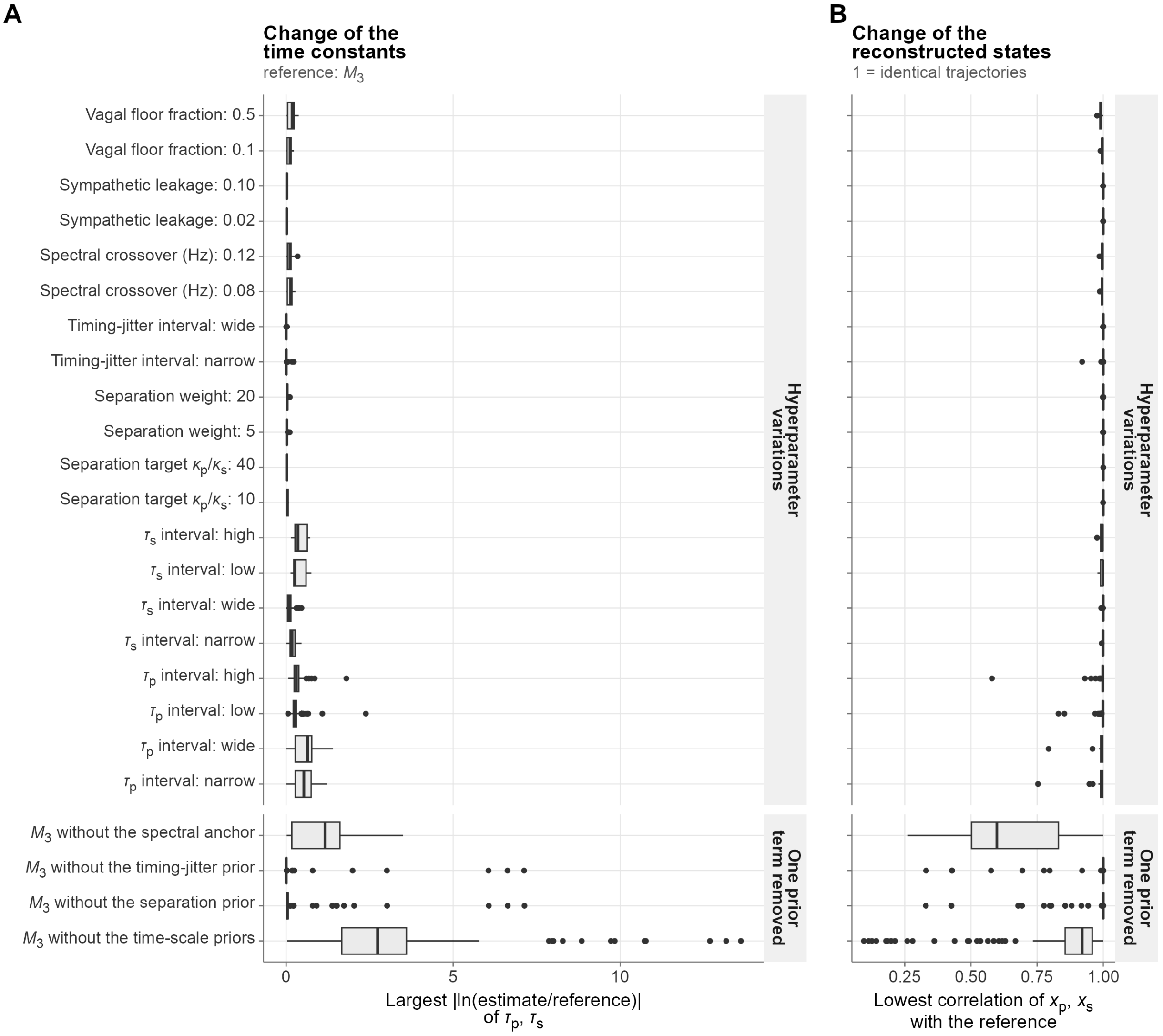


Figure S 2: Sensitivity to the hyperparameters. (A) Largest absolute log-ratio of $\tau_p$ and $\tau_s$ relative to the reference. (B) Lowest correlation of the reconstructed states with the reference trajectories.

*Results of the sensitivity analyses.* Across 150 recordings, varying one hyperparameter at a time changed the time constants by a median absolute log-ratio of at most 0.64 (largest for the $\tau_p$ prior interval (wide)) and the lowest median correlation with the reference trajectories was 0.99. Removing one prior component at a time, the largest change of the time constants occurred without the time-scale priors (median absolute log-ratio 2.40). By dataset, $\tau_p$ lay inside its 95% prior interval in 30% (prcp), 55% (fantasia) and 3% (exercise) of the recordings and $\tau_s$ in 90% (prcp), 100% (fantasia) and 99% (exercise). Supplementary figure S2 shows the changes of the time constants and of the reconstructed states for each variation.

### 8 Recovery-time benchmark

*Definitions.* For each state, smoothed with a 10-s moving average, with the excursion measured from the rest level, four definitions of recovery were compared: (a) *sustained return*, the first time after the end of exercise at which the excursion remains within 1 rest SD for 20 s; (b) *first return*, the first time within 1 rest SD; (c) *90% recovery*, the first time at which the excursion falls below 10% of its end-of-exercise value; and (d) *exponential constant*, the time constant of an exponential decay fitted to the excursion after the end of exercise. Recordings that did not meet the criterion within the observation window were treated as right-censored at the end of the recording (Kaplan–Meier medians are reported), and recordings without an excursion larger than 1 rest SD were flagged.

*Benchmark.* Simulated rest–exercise–recovery recordings were generated with known recovery constants ($\tau_{off,p}$ 12–60 s, $\tau_{off,s}$ 25–100 s), each profile realized twice with independent noise. Each realization was fitted ($M_3$), events were detected, and the four recovery definitions (main text) were computed for each state under each end-of-exercise rule. The score of a definition was the success proportion multiplied by the mean of the two correlations, and the end-of-exercise rule with the smallest median absolute error was retained. Table S 6 reports, for the retained end-of-exercise rule, the proportion of recordings with a defined estimate, the Spearman correlation with the true constant and the test-retest Spearman correlation between the two realizations.

Table S 6: Benchmark of recovery-time definitions on simulated recordings (mean over the two states).

| Definition | Defined estimate | Rank correlation with truth | Test-retest rank correlation | Score |
|---|---|---|---|---|
| 90% recovery of the excursion | 96% | 0.57 | 0.59 | 0.56 |
| exponential time constant | 87% | 0.57 | 0.57 | 0.49 |
| first return to within 1 rest SD | 82% | 0.49 | 0.61 | 0.45 |
| sustained return to within 1 rest SD | 69% | 0.49 | 0.57 | 0.37 |

The definition with the highest score was 90% recovery of the excursion, and the end-of-exercise rule with the smallest median absolute error was offset time of the fitted exercise profile (100 simulated profiles; supplementary figure S3). The ranking depends on the simulated profiles, on the 10-s smoothing and on the tolerance of 1 rest SD, and should be revisited if the real recovery curves differ qualitatively (for example, non-monotonic recovery).

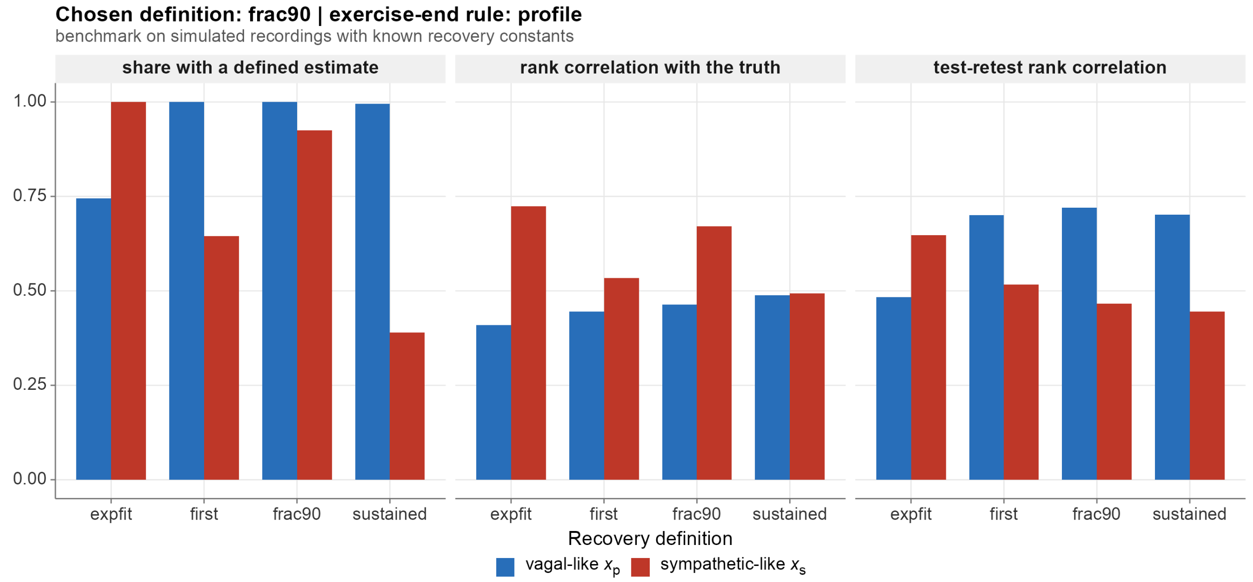


Figure S 3: Benchmark of recovery-time definitions: proportion of defined estimates, rank correlation with the true constant, and test-retest rank correlation, for each state.

### 9 Calibration of the innovation diagnostics

To examine the behavior of the Kolmogorov–Smirnov and PACF diagnostics when the model is correctly specified and its parameters are estimated from the same data, recordings were simulated from the model (time scales within the prior ranges) and fitted with $M_3$. Across 100 simulated recordings, the KS distance exceeded the approximate critical value $1.36/\sqrt{N}$ in 0% (nominal level 5% if parameters were known), the mean number of PACF lags outside the bounds was 1.58 of 40 (nominal 2.0), and the SD of the standardized innovations had a mean of 1.001 (nominal 1). Supplementary figure S4 summarizes these checks together with the numerical verification of the bound.

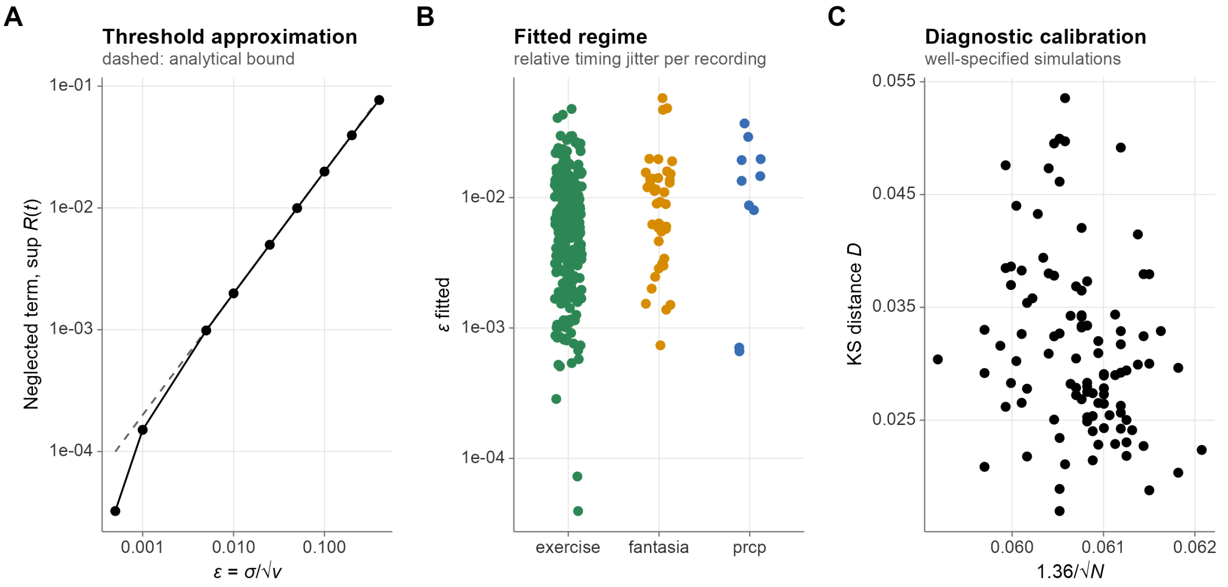


Figure S 4: Numerical checks. (A) Neglected reflection term and its bound as a function of $\varepsilon$. (B) Fitted $\varepsilon$ in the recordings. (C) KS distance versus critical value in well-specified simulations.

The closed-form process covariance of the Ornstein–Uhlenbeck integrals (including the Taylor branch used for $\kappa\Delta t < 10^{-3}$) was compared with numerical integration by the Van Loan method over a grid of rates, intervals and variances; the maximum relative error was 2.5e-09 for the closed form and 7.2e-11 for the Taylor branch. The compiled filter used to accelerate the multi-start analyses reproduced the engine's R filter (likelihood, filtered states and implied RR) with a maximum discrepancy of 2e-13.

### 10 Data preparation and its checks

*Data files.* Each recording was stored as a CSV file with one row per ECG-derived RR interval: the time of the end of the interval (`time`, s; record clock for PRCP and Fantasia, cumulative RR clock for the exercise cohort), the RR interval (`rri`, s; `rri` equals the difference of successive `time` values) and, when available, systolic and diastolic blood pressure and ABP of the cardiac cycle of the interval, respiration (Fantasia; mean over the interval) or the tilt-table angle (PRCP; mean over the interval). Beat annotations gave the RR intervals; for PRCP, the protocol annotations of each record (start and end of every tilt, the stand-up maneuvers and the returns to supine) were also read and stored as a table of maneuvers. Blood pressure was read from the arterial waveform in a window of one pulse per beat, shifted by the ECG-to-pulse delay estimated for each record from the ensemble-averaged waveform, and implausible values were set to missing. RR intervals, blood pressure and respiration were cleaned with an adaptive running-median/median-absolute-deviation gate (window of 50 beats; values beyond 3.5 median absolute deviations replaced by linear interpolation; two passes), a setting chosen on simulated series with injected artifacts because it altered few artifact-free beats (about 0.2%) while correcting most artifacts (about 99.7%; supplementary material). In PRCP, RR intervals between 0.3 and 2 s that fell within 5 s before to 30 s after a maneuver boundary were left unchanged, because genuine fast changes at these transitions (for instance a return to supine) resemble outliers. For PRCP, arterial pressure between an annotated loss of the Finapres signal and its switch-on was set to missing. Not all recordings contained all signals: blood pressure, respiration or the tilt angle was missing for some subjects, and the analyses that use them were restricted to the recordings with at least 80% valid values in the analysis window, with the number of recordings reported for each analysis. The latent state model itself uses only the RR intervals. The `rri` column was used as written in all datasets. All time-based analyses use the file's `time` column as the common clock for the latent states and for the covariates. For PRCP, arterial pressure between an annotated loss of the Finapres signal and its switch-on was set to missing.

*Preparation script.* The recordings were prepared from the WFDB files (Fantasia, PRCP) and the text files of RR intervals (exercise cohort). The beat table is built once from the ECG annotations (beat symbols only), so that `time[k] - time[k-1] = rri[k]` exactly and the covariates of a row refer to the same cardiac cycle. For blood pressure, the delay between the R peak and the foot of the arterial pulse is estimated per record (minimum of the ensemble-averaged waveform), systolic and diastolic pressure are the maximum and minimum of the waveform in the pulse window of each beat, values outside plausible ranges (systolic 50–300 mmHg, diastolic 20–200 mmHg, pulse pressure of at least 10 mmHg) are set to missing, and a signal with fewer than 80% valid values is treated as absent. For PRCP, the RR intervals come from the ECG beats and the arterial pressure from the ABP waveform in the pulse window of each ECG beat. For PRCP, the protocol annotations (`.anI`: start and end of every slow and rapid tilt, stand-up from supine, back to supine) are parsed into `prcp_events.csv`, one row per maneuver; labels that are not recognized are reported by the script and are not used. A quality table (`prep_qc.csv`) records, per recording, the number of beats, the proportion of RR values changed by cleaning, gaps, the estimated delay and the signals present. The script was verified by running it on synthetic WFDB records with known truth (`tests/suites/test_prep.R`): the RR intervals, beat times, pressure values, respiration and angle written to the CSV files matched the generating values; the delay was recovered; injected pressure artifacts and dropouts were cleaned; absent channels produced missing columns; and the resulting files were read by the analysis loader and fitted by the model.

*Cleaning gate.* The gate replaces values beyond a multiple of the local median absolute deviation (running median and MAD over 50 beats) by linear interpolation, and is repeated. Settings were compared on simulated resting series with respiratory modulation and about 3% injected artifacts (ectopic pairs, long and short intervals, isolated spikes; `tools/checks/benchmark_cleaning_gate.R`). The gating strategy (setting 3.5 median absolute deviations with two passes) altered about 0.2% of artifact-free beats, corrected about 99.7% of the artifacts and left RMSSD within about 2% of the artifact-free value. In PRCP, RR intervals between 0.3 and 2 s lying within 5 s before to 30 s after a maneuver boundary are not replaced: in a record with six maneuvers, the gate would otherwise have altered four to five beats of up to 0.4 s at each return to supine, a genuine fast change, and this protection left the proportion of altered beats of that record at 0.9% instead of 1.1%.

### 11 Event detection

*Maneuvers, labels and baselines.* Each tilt was labeled slow or rapid from the annotated tilt whose ascent contained the angle-derived onset (within 5 s); the wording of the annotations differs between records and is parsed accordingly. For recordings without a usable angle signal the annotated tilt times were used (onset: start of the ascent; return: end of the descent). Every maneuver, not only tilts, spaces the baseline of the next one. For tilt recordings, which can contain several perturbations, the heart-rate segmentation allowed up to thirteen segments (six perturbations) with a minimum segment length of 30 s, and for the exercise recordings one to three segments. The onset was defined as an upward change point, and recovery as the next downward change point that brought heart rate back at least halfway to its pre-perturbation level (a dip inside a perturbation did not end it); for the exercise recordings only the first upward change point was used, and the parametric profile was fitted over exercise durations of 40–360 s.

*Tilt angle.* The (beat-wise) angle is median-filtered (5 beats) and the baseline taken as its median over the first 60 s. The onset is the first time at which the angle exceeds the baseline by more than 10° for at least 5 s, and the return the first later time at which it lies within 10° of baseline for 5 s. Plateaus are runs of at least 20 s of the same angle rounded to 5°, and the state means over the second half of a plateau are used for the dose-response analysis.

*Heart rate.* Heart rate (Hz) is interpolated at 1 Hz and averaged in bins of $b = 5$ s. For $K = 1, 2, 3$ segments, the optimal piecewise-constant partition with segments of at least 30 s is found by exact dynamic programming (minimizing the residual sum of squares). The number of segments is selected by the Bayesian information criterion $n \ln(\mathrm{SSE}/n) + (2K - 1) \ln n$ among partitions in which adjacent segment means differ by at least $\Delta_{min}$ (default 0.10 Hz, approximately 6 bpm); the best-scoring $K$ satisfying the rule is retained, and $K = 1$ (no event) otherwise. Change points are placed midway between adjacent bins. The BIC treats the binned values as independent, which is approximate for a smooth heart-rate series, and the minimum-shift rule is the operative safeguard against false detections.

For exercise recordings, the profile $HR(t) = h_0 + A\,g(t)$ is also fitted, with $g(t) = 0$ before the onset $t_{on}$, $g(t) = 1 - e^{-(t-t_{on})/\tau_{on}}$ until the end of exercise $t_{off}$, and $g(t) = g(t_{off})\,e^{-(t-t_{off})/\tau_{off}}$ afterwards. For each combination of the nonlinear parameters ($t_{on}$, $t_{off}$, $\tau_{on}$, $\tau_{off}$) on a coarse grid (about 80 points over the recording; exercise duration 40–360 s), $h_0$ and $A$ are obtained by linear least squares, and the best grid point is refined by Nelder–Mead. Three candidate rules for the end of exercise are compared: the heart-rate peak after the onset (smoothed over three bins), the second change point, and $t_{off}$.

Table S 7: Heart-rate-based event detection on simulated recordings with known timing: detection rate for tilt-like recordings and false-detection rate in resting recordings as a function of the minimum-shift rule.

| Minimum shift (Hz) | Detection rate (tilt-like) | False-detection rate (rest) |
|---|---|---|
| 0.05 | 100% | 29% |
| 0.08 | 100% | 6% |
| 0.10 | 100% | 1% |
| 0.12 | 97% | 0% |

In these simulations, the median onset error was 8.4 s for tilt-like recordings, and the median absolute error of the end of exercise was 23.4 s, 25.8 s, 2.7 s for the peak, second-change-point and profile rules, respectively. These values characterize the detection procedure under the simulated profiles, which are idealized. For recordings with repeated tilts, simulated with three tilts of 180 s separated by 240 s, 100% of the tilts were found (median onset error 7.4 s), with 0.0 additional events per recording. In the PRCP recordings, the accuracy against the tilt angle is shown in Figure S 5: each angle-derived onset was matched to the nearest heart-rate-detected onset within 60 s, and heart-rate events matching no tilt (other maneuvers that raise heart rate) were counted.

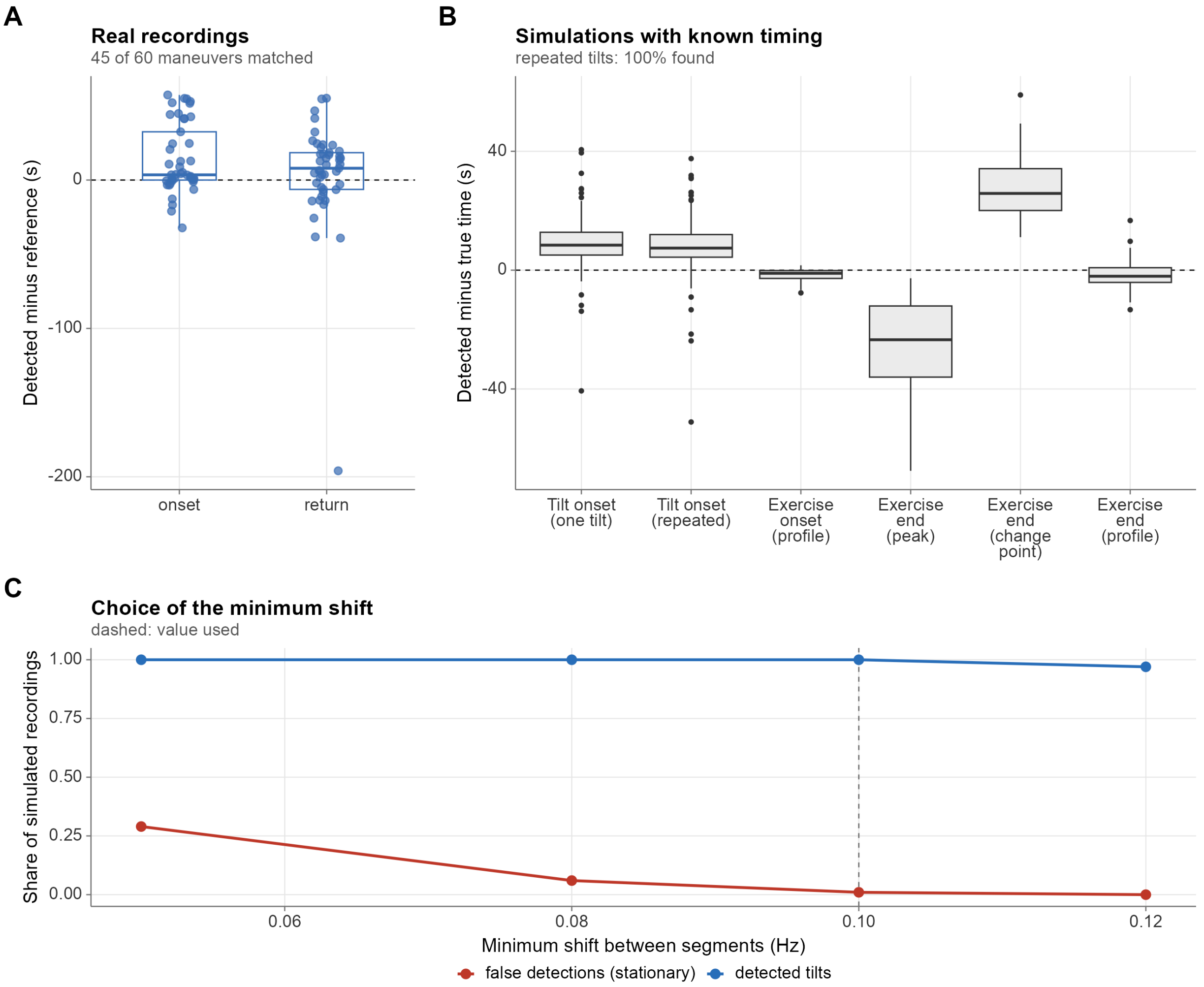


Figure S 5: Event detection. (A) Heart-rate-detected minus angle- or annotation-derived onset and return of each maneuver (tilts and stand-ups) in the PRCP recordings. (B) Detected minus true times in simulated recordings with known timing. (C) False-detection rate in stationary simulated recordings and detection rate of simulated tilts as a function of the minimum shift between segments (dashed: value used).

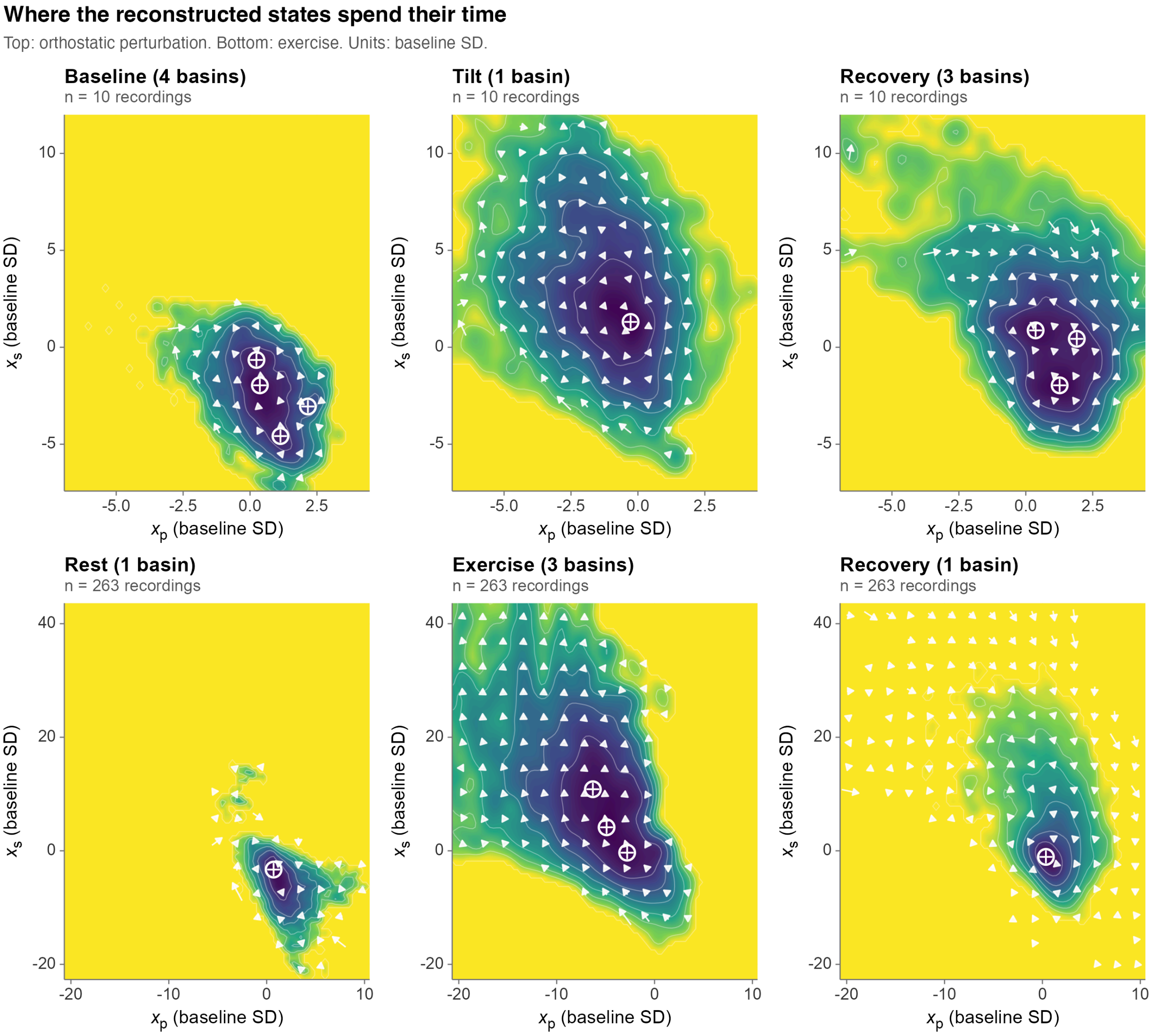


Figure S 6: State-space landscapes of the perturbation (top) and exercise (bottom) conditions. Each panel shows the pooled density of the filtered states in one phase (every recording contributes equally), with its basins (rings) and the mean flow of the states (arrows). Coordinates are in units of each recording's baseline SD; dark = high occupancy. The map is descriptive and is not a potential function estimated from the model.

### 12 Responses by type of maneuver (PRCP)

The PRCP protocol comprises slow tilts, rapid tilts and stand-ups from supine, which the annotations of each record identify. The pooled analysis in the main text uses the tilts; Figure S 7 shows the event-locked response of the two states, heart rate and ABP for each kind of maneuver, every event relative to its own baseline, and Table S 9 the paired within-recording contrasts. The stand-ups are active changes of posture during which the table angle does not change, so they could not be identified from the angle. The contrasts are exploratory: recording medians, Wilcoxon signed-rank test, Benjamini-Hochberg across the table, and few recordings.

The protocol annotations distinguish slow tilts, rapid tilts and stand-ups (Table S 8). Both states changed in the expected direction in 20 of 20 slow tilts (100%), 19 of 20 rapid tilts (95%) and 20 of 20 stand-ups (100%). The median standardized change of the vagal-like state was −1.5, −1.6 and −1.4 baseline SDs and that of the sympathetic-like state 9.4, 7.2 and 7.2, in the same order. Paired within-recording contrasts (rapid minus slow tilt, and stand-up minus tilts; 8 exploratory contrasts on 10 pairs of recording medians) had $p < 0.05$ in 4 (3 after Benjamini-Hochberg adjustment): rapid - slow tilt for the onset latency of $x_p$ (median difference −21.5, $p$ = 0.006), rapid - slow tilt for the onset latency of $x_s$ (median difference −30.5, $p$ = 0.002), stand-up - tilts for the onset latency of $x_p$ (median difference −11.0, $p$ = 0.041) and stand-up - tilts for the onset latency of $x_s$ (median difference −21.5, $p = 0.004$). Onset latencies are measured from the annotated time of the maneuver, which marks the start of the posture change and not a calibrated onset of the stimulus (a negative latency means that the response starts before the annotated time), so contrasts in latency between kinds are not interpreted physiologically.

Table S 8: Responses by type of maneuver in the PRCP recordings (median [25th, 75th percentile] over events; descriptive).

| Maneuver | Events (rec.) | Both states as expected | $\Delta x_p$ (SD) | $\Delta x_s$ (SD) | Onset $x_p$ (s) | Onset $x_s$ (s) | ΔABP (mmHg) |
|---|---|---|---|---|---|---|---|
| slow tilt | 20 (10) | 100% | −1.5 [−2.9, −0.8] | 9.4 [4.4, 12.2] | 14 [4, 24] | 29 [14, 40] | 4.9 [−0.5, 7.9] |
| rapid tilt | 20 (10) | 95% | −1.6 [−2.9, −0.8] | 7.2 [5.5, 11.7] | −4 [−16, −2] | −4 [−20, 5] | 3.9 [−0.8, 8.1] |
| stand-up | 20 (10) | 100% | −1.4 [−2.8, −0.7] | 7.2 [3.8, 9.3] | −7 [−14, −4] | −8 [−14, −4] | 8.8 [2.8, 10.9] |

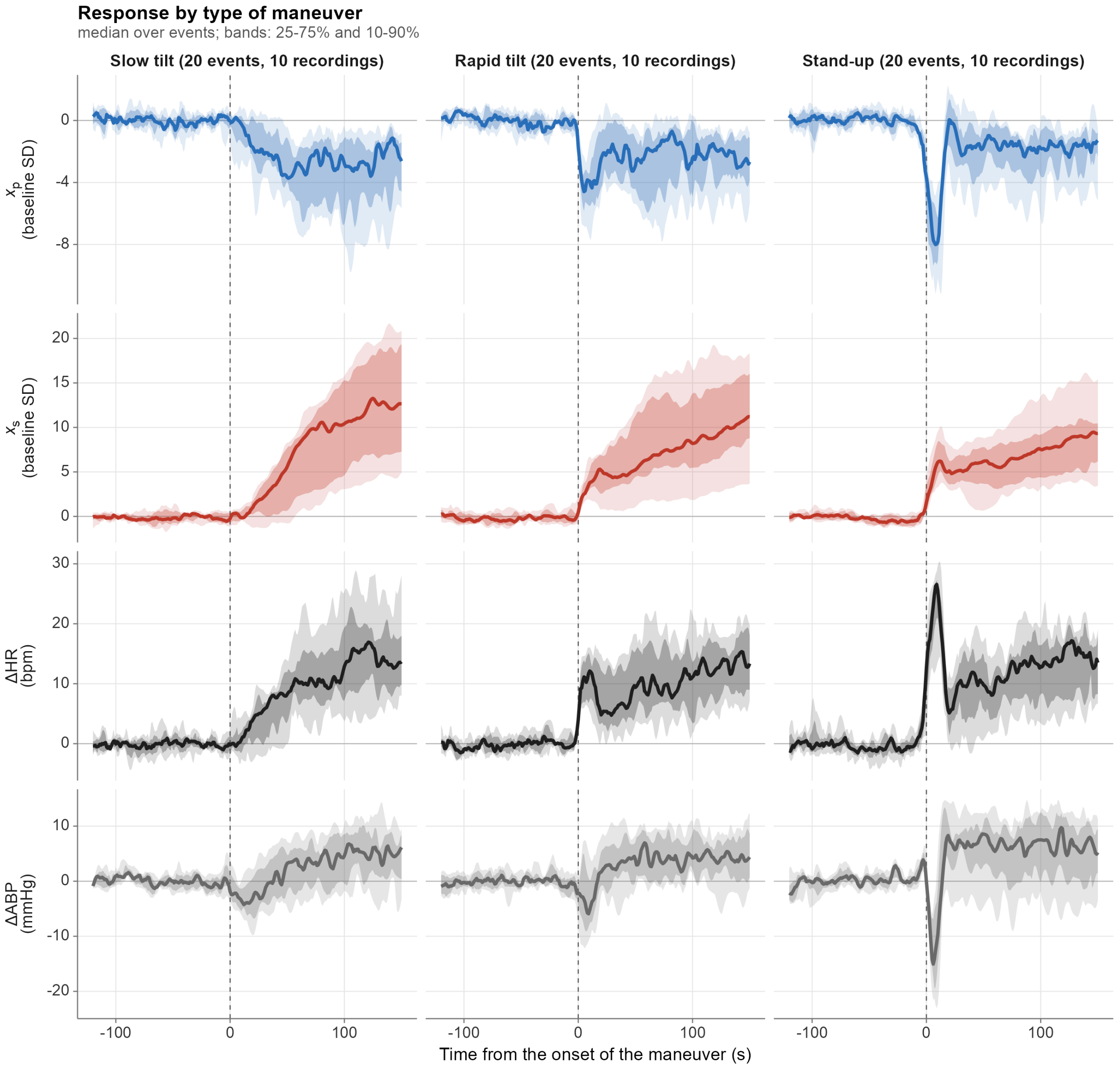


Figure S 7: Responses by type of maneuver. Rows: vagal-like state, sympathetic-like state (in baseline SDs), heart rate and ABP (changes); columns: one per kind of maneuver (slow tilt, rapid tilt and stand-up; a tilt that the annotations do not label appears as "tilt"). Line: median over events; bands: 25-75% and 10-90%; each event is relative to its own baseline, and the scale of a row is shared across columns.

Table S 9: Paired within-recording contrasts between kinds of maneuver in the PRCP recordings (exploratory).

| Contrast | Variable | Pairs | Median difference | $p$ | $p$ (BH) |
|---|---|---|---|---|---|
| rapid - slow tilt | Change of $x_p$ (SD) | 10 | −0.12 | 0.131 | 0.209 |
| rapid - slow tilt | Change of $x_s$ (SD) | 10 | −1.47 | 0.432 | 0.493 |
| rapid - slow tilt | Onset of $x_p$ (s) | 10 | −21.50 | 0.006 | 0.016 |
| rapid - slow tilt | Onset of $x_s$ (s) | 10 | −30.50 | 0.002 | 0.016 |
| stand-up - tilts | Change of $x_p$ (SD) | 10 | −0.14 | 0.846 | 0.846 |
| stand-up - tilts | Change of $x_s$ (SD) | 10 | −2.14 | 0.232 | 0.310 |
| stand-up - tilts | Onset of $x_p$ (s) | 10 | −11.00 | 0.041 | 0.082 |
| stand-up - tilts | Onset of $x_s$ (s) | 10 | −21.50 | 0.004 | 0.016 |

### 13 Respiration, blood-pressure and lagged-correlation estimators

All series are beat-wise and are resampled to 4 Hz by natural cubic splines before spectral estimation. Spectra are estimated with a tapered (10%), detrended periodogram smoothed over about 0.01 Hz, and power is reported one-sided (twice R's two-sided density).

*Respiratory frequency* is the frequency of the spectral maximum of the respiration signal between 0.10 and 0.50 Hz. *Respiratory-band SD and variance share* of a state are the square root and the fraction of the one-sided power of that state within the respiratory frequency $\pm 0.04$ Hz (relative to its total power). *Coherence* is the magnitude-squared coherence between the state and respiration averaged over the same band and weighted by the respiratory power in each frequency bin, so that bins without respiratory power, in which coherence is noise, do not dominate. Coherence does not depend on amplitude and cannot by itself distinguish a state that carries a respiratory oscillation from one that carries it at low amplitude.

*Baroreflex alpha index.* With $P_{RR}$ and $P_{SBP}$ the spectra of the RR intervals (ms) and the systolic pressure (mmHg) and $\gamma^2$ their squared coherence, $\alpha_B = \sqrt{P_{RR}/P_{SBP}}$ over the frequencies of band $B$ (LF 0.04–0.15 Hz; HF 0.15–0.40 Hz) at which $\gamma^2 \geq 0.5$ (ms/mmHg); it is undefined when no frequency in the band meets the criterion. In a numerical check in which RR was an 8 ms/mmHg gain of systolic pressure delayed by 2 s with added noise, the estimator returned the generating gain within 15%.

*Lagged correlation.* Both series are placed on a 1-Hz grid and the centered 60-s moving mean is removed from each (the first and last 30 s are discarded). The cross-correlation is computed for lags of $\pm 30$ s, the peak absolute value and its lag are reported (a positive lag means that the second series lags the first), and a surrogate $p$-value is obtained from the peak absolute correlations of 200 circular shifts of the second series (shifts of at least 60 s), which preserve its autocorrelation. In a numerical check with a known 5-s delay, the lag was recovered and the surrogate $p$-value was below 0.05.

### 14 Secondary analyses of the perturbation

*Negative control for the phase contrast.* The resting baseline of each analyzed tilt (60 to 180 s long, ending at the onset) was split into two halves, and the second half was contrasted with the first as the tilt is contrasted with the baseline in the main analysis: difference of the mean of each state, in SDs of the first half, with the expected signs (decrease of $x_p$, increase of $x_s$). No maneuver separates the two halves, so any expected pattern arises from drift, noise or the construction of the contrast. Halves with fewer than 20 beats were skipped; only tilts that have both contrasts were compared. The share of events with both signs as expected is given with exact (Clopper–Pearson) 95% intervals, which are descriptive because the tilts of a recording are not independent, and the absolute standardized changes were compared between the tilt and the placebo contrast with an exact one-sided paired signed-rank test on recording medians. The baseline begins 120 s after the previous maneuver returned, so it may still contain part of the recovery from it, and the placebo share can therefore exceed the 25% expected for independent random signs.

*Comparison with a CWT moving spectrum.* The temporal resolution of the reconstruction was compared with the continuous-wavelet-transform (CWT) moving band power of the heart rate (LF 0.04–0.15 Hz, HF 0.15–0.40 Hz, cone of influence masked), quantified as the time at which each series first leaves its baseline band (baseline mean $\pm 2$ SD, sustained for 5 s) relative to the tilt onset.

*Arterial pressure.* For the recordings with arterial pressure, the same phase contrast (excluding the first 30 s after the onset) and step-response fit were applied to systolic, diastolic and mean arterial pressure (ABP) and to pulse pressure, as independently measured responses, and the temporal relation between each state and ABP was examined by the lagged correlation of the two detrended series, with a surrogate $p$-value from circular shifts of ABP that preserve its autocorrelation (section "Respiration, blood-pressure and lagged-correlation estimators").

*Dose–response.* For recordings with several tilt levels, the mean of each state, of heart rate and of ABP over the second half of each plateau was related to the angle by the within-recording Spearman correlation (at least four levels).

*Kinds of maneuver.* The pooled analyses use the tilts (slow and rapid together); because the original description of the dataset reports that the initial heart-rate increase of the rapid tilt and the stand-up is absent in the slow tilt [6], the same quantities were also computed by kind of maneuver and compared within recordings (rapid versus slow tilts, and stand-ups versus tilts; Wilcoxon signed-rank test on recording medians, Benjamini–Hochberg across contrasts; exploratory), as reported in section "Responses by type of maneuver (PRCP)".

*Results.* In the 10 recordings with arterial pressure, ABP changed by a median of 4.3 mmHg (pulse pressure −7.2 mmHg) with an onset 13.5 s after the tilt onset; after removing slow trends, the peak lagged correlation of ABP with the states was −0.19 for $x_p$ (lag 2 s; surrogate $p < 0.05$ in 100% of recordings) and 0.21 for $x_s$ (lag 2 s; 100%). These correlations are weak in magnitude. The sign differed between recordings: the correlation with $x_p$ was negative in 6 and positive in 4 of 10 recordings; that with $x_s$ was negative in 3 and positive in 7 of 10 recordings. The sympathetic-like state left its baseline band 13 s and the LF band power 28 s after the tilt onset (median), and for the vagal-like state and the HF band power 8 and −13 s, respectively. The sympathetic-like state therefore left its baseline band earlier than the LF band power, and the vagal-like state not earlier than the HF band power (a negative value for the band power means that it departed from its baseline before the tilt onset, which is consistent with the spreading of the moving window across the transition and not with an anticipatory response); the comparison with the moving spectrum is not uniformly in favor of the filter. In the state plane, the strongest basin of the occupied region (the mode of the density of the filtered states) was displaced from its position in the baseline phase by 3.3 baseline-SD units to the tilt phase and 0.9 baseline-SD units to the recovery phase (Supplementary Figure S6); the number of basins detected was 4 in the baseline phase, 1 in the tilt phase, 3 in the recovery phase. The landscape is a descriptive map of where the states lie and is not a fitted potential of the model. The comparison with the CWT moving spectrum is shown in Figure S 8.

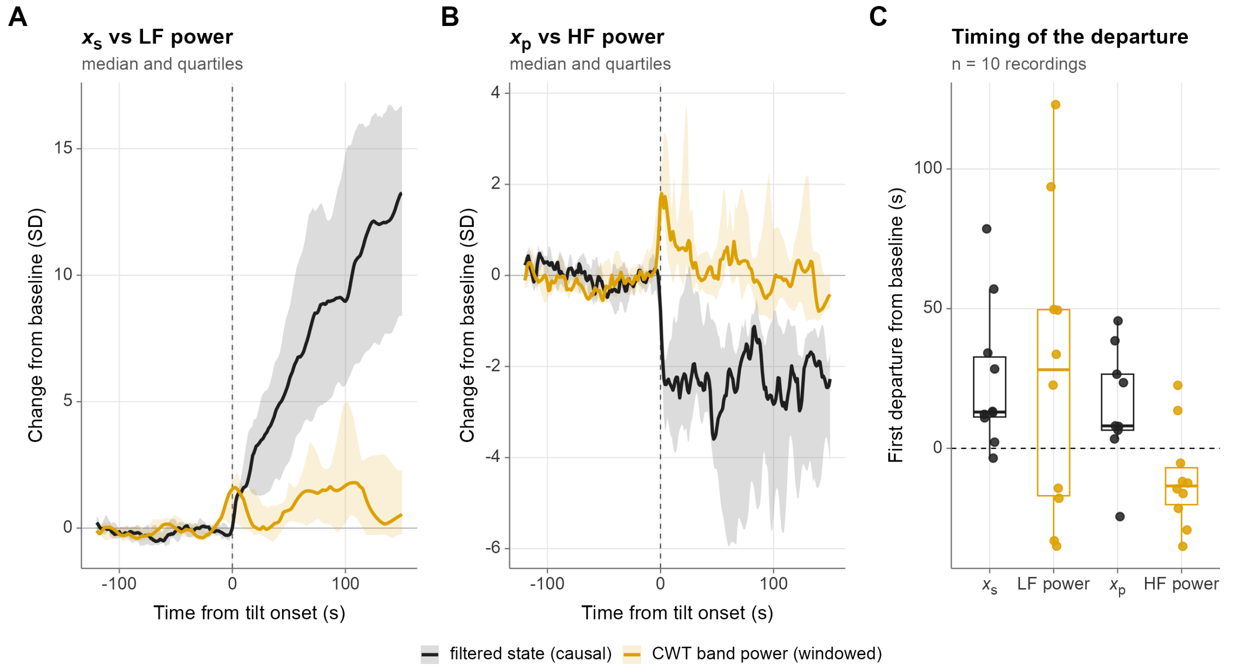


Figure S 8: Temporal resolution of the reconstruction relative to a CWT moving spectrum during the perturbation. (A) Median change of the sympathetic-like state (forward filter) and of the CWT LF band power around the tilt onset, in baseline SDs (bands: quartiles over recordings). (B) The same for the vagal-like state and the HF band power; the window required for frequency resolution spreads the CWT power around the transition. (C) Time of first departure from the baseline band, relative to the tilt onset, across recordings.

### 15 Additional analyses of the Fantasia recordings

*Group contrasts.* The contrasts between young and older adults in the latent summaries and in the conventional HRV metrics are listed in Table S 10 (shown in Figure 3 of the main text).

Table S 10: Group contrasts in the Fantasia database (older minus young): medians, Hodges–Lehmann shift, Cliff's delta with bootstrap 95% interval, and Benjamini–Hochberg adjusted p-values within family.

| Variable | Family | Median (young) | Median (older) | HL shift | Cliff's $\delta$ [95% CI] | $p$ (BH) |
|---|---|---|---|---|---|---|
| $\nu_0$ | latent | 1.03 | 0.967 | −0.0793 | −0.31 [−0.64, 0.04] | 0.111 |
| $\tau_p$ | latent | 3.16 | 5.37 | 2.22 | 0.72 [0.45, 0.93] | <0.001 |
| $\tau_s$ | latent | 165 | 167 | 0.638 | 0.04 [−0.32, 0.41] | 0.900 |
| $\lambda_P$ | latent | 2.61 | 1.57 | −0.873 | −0.53 [−0.82, −0.19] | 0.006 |
| $\hat{\sigma}^2_{sys}$ | latent | 0.00195 | 0.000370 | −0.00138 | −0.71 [−0.91, −0.44] | <0.001 |
| RR timing jitter (ms) | latent | 8.81 | 8.43 | −0.0953 | −0.01 [−0.38, 0.36] | 0.946 |
| SD of $x_p$ | latent | 0.0568 | 0.0218 | −0.0335 | −0.81 [−0.97, −0.58] | <0.001 |
| SD of $x_s$ | latent | 0.0199 | 0.0116 | −0.00856 | −0.61 [−0.85, −0.31] | 0.002 |
| SD ratio $x_p/x_s$ | latent | 2.61 | 2.08 | −0.605 | −0.54 [−0.81, −0.23] | 0.005 |
| $\ln(\kappa_p/\kappa_s)$ | latent | 3.97 | 3.40 | −0.537 | −0.69 [−0.92, −0.42] | <0.001 |
| Share of $x_p$ variance | latent | 0.872 | 0.810 | −0.0726 | −0.54 [−0.81, −0.23] | 0.005 |
| Mean HR | HRV | 61.8 | 57.9 | −4.80 | −0.33 [−0.65, 0.03] | 0.108 |
| SDNN | HRV | 65.1 | 33.7 | −31.0 | −0.71 [−0.92, −0.43] | <0.001 |
| RMSSD | HRV | 54.6 | 22.1 | −31.6 | −0.81 [−0.96, −0.59] | <0.001 |
| LF/HF | HRV | 1.25 | 1.96 | 0.506 | 0.26 [−0.11, 0.60] | 0.172 |

*Quantities that are comparable across people.* The tonic level of each drive is not identified from RR intervals ($\nu_0$, $\mu_p$ and $\mu_s$ enter the likelihood only through $\nu_0 - \mu_p + \mu_s$), so what can be compared is the variability of each drive in physical units, the time constants and the mean heart rate. The stationary SD of the vagal-like and sympathetic-like drives is $\sqrt{\lambda_P \sigma_{sys}^2}$ and $\sqrt{\sigma_{sys}^2}$ (Hz; multiplied by 60, a bpm equivalent); their ratio is the spectral anchor, so the split inherits its accuracy. Table S 11 lists them by group with the agreement between the two windows of the stability analysis.

Table S 11: Quantities comparable across people in the Fantasia database: medians by group, Cliff's delta (older versus young) with bootstrap 95% interval, and agreement between two windows of the same recording (ICC of the log values). Descriptive; no adjusted p-values.

| Quantity | Median (young) | Median (older) | Cliff's $\delta$ [95% CI] | ICC (log) |
|---|---|---|---|---|
| Mean heart rate (bpm) | 61.8 | 57.9 | −0.33 [−0.65, 0.03] | — |
| SD of the vagal-like drive (bpm equivalent) | 4.60 | 1.53 | −0.86 [−0.99, −0.66] | 0.95 |
| SD of the sympathetic-like drive (bpm equivalent) | 2.65 | 1.15 | −0.71 [−0.91, −0.44] | 0.90 |
| $\tau_p$ (s) | 3.16 | 5.37 | 0.72 [0.45, 0.93] | 0.88 |
| $\tau_s$ (s) | 165 | 167 | 0.04 [−0.32, 0.41] | 0.31 |

The stationary SD of the vagal-like drive was 1.5 bpm (equivalent) in older and 4.6 in young adults (Cliff's $\delta$ = −0.86), that of the sympathetic-like drive 1.2 and 2.6 bpm ($\delta$ = −0.71), and the vagal time constant 5.4 and 3.2 s ($\delta$ = 0.72). Between two windows of the same recording (ICC of the log values) the agreement was 0.95 for the vagal-like SD, 0.90 for the sympathetic-like SD, 0.88 for $\tau_p$ and 0.31 for $\tau_s$. These are descriptive comparisons of 20 young and 20 older adults; a net tone computed from an age-based intrinsic rate would differ between the groups by construction of the formula, not from the data.

*Methods.* The leave-one-out $R^2$ with which the conventional HRV metrics predict each latent summary (low values indicate that the summary is not linearly recoverable from them) and the leave-one-out discrimination (AUC) of young versus older adults using HRV metrics, latent summaries or both were computed on the same recordings. For the subjects with a respiration signal, the dominant respiratory frequency (spectral peak between 0.10 and 0.50 Hz) was compared with the HF band (0.15–0.40 Hz), since the spectral anchor assumes that vagal modulation lies mainly in that band, and the respiratory coupling of each reconstructed state was quantified by the SD of the state within the respiratory band (respiratory frequency $\pm$ 0.04 Hz), its share of the total variance of the state in that band and the respiratory-power-weighted coherence with respiration; coherence is insensitive to amplitude, so the band SD and variance share are the primary measures, and the vagal-like and sympathetic-like states were compared within recordings (paired Wilcoxon test). For the subjects with blood pressure, mean systolic pressure, ABP and pulse pressure were summarized and baroreflex sensitivity was estimated by the alpha index in the LF and HF bands at frequencies where the squared coherence exceeds 0.5; these quantities were compared between groups with the same contrasts as in the main text and related to the latent summaries by Spearman correlations (pooled over groups, exploratory). Estimators are detailed in section "Respiration, blood-pressure and lagged-correlation estimators". The stability of the representation was examined by refitting two non-overlapping windows of each recording (600–1200 s and 1200–1800 s).

*Results.* The leave-one-out AUC for separating the groups was 0.83 with HRV metrics, 0.86 with latent summaries and 0.93 with both, and HRV metrics predicted the latent summaries with a median leave-one-out $R^2$ of 0.42 (low values indicate variation that is not linearly predicted by these metrics, which may include estimation error as well as additional information). The AUC values are uncertain with these group sizes and were not compared formally. In the 40 recordings with respiration, the respiratory frequency lay within the HF band in 90%, and the SD of the vagal-like state within the respiratory band was larger than that of the sympathetic-like state in 100% of recordings (median ratio 8.7), as expected from the faster time scale assigned to the vagal-like state. The respiration measures are shown in Figure S 9.

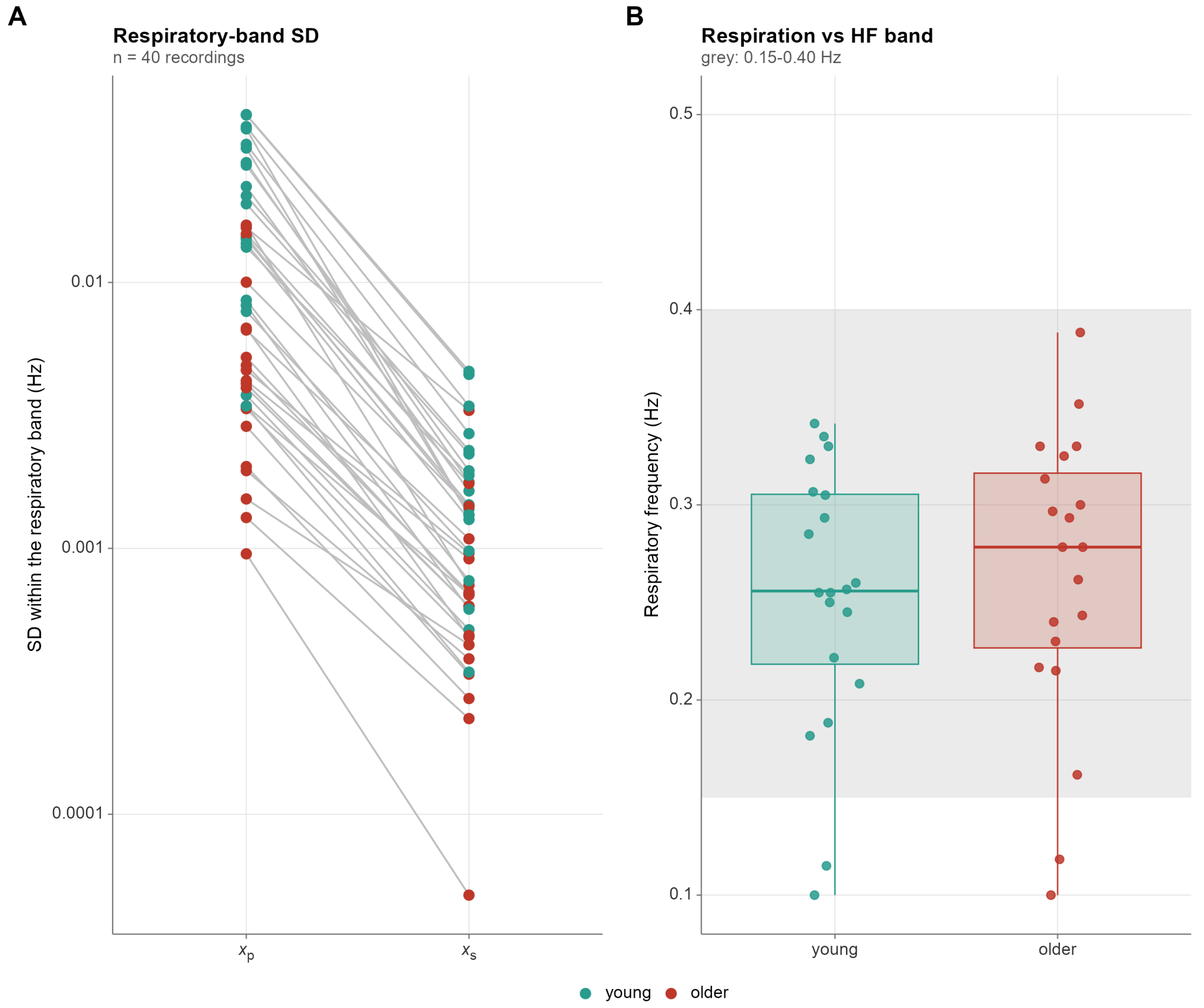


Figure S 9: Respiration in the Fantasia database (subjects with the signal; blood pressure is not available in these recordings). (A) SD of each reconstructed state within the respiratory band, paired within recordings. (B) Respiratory frequency by group, with the HF band shaded.

### 16 Fit and innovation diagnostics of the engine specification

The summaries of the filtered fit, the Kolmogorov–Smirnov distance and the partial autocorrelation of the standardized innovations of $M_3$ are given in the main text; Figure S 10 shows the distributions across recordings and the estimated time constants of the two drives against the 95% intervals of their priors. The calibration of the critical value under a correctly specified model is described in the “Calibration of the innovation diagnostics” section.

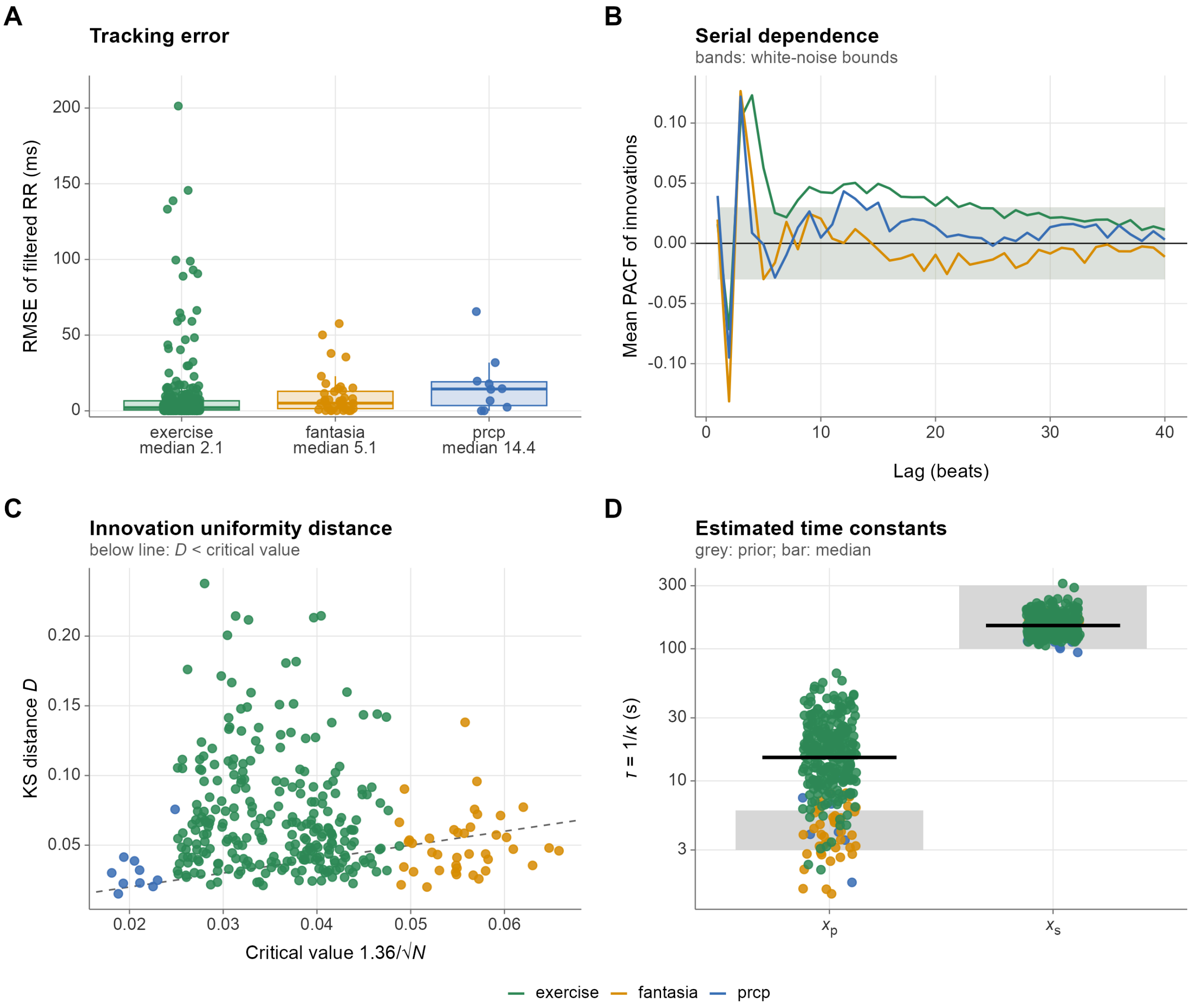


Figure S 10: Fit and innovation diagnostics of the engine specification. (A) Distribution of filtered (in-sample) tracking error across recordings. (B) Cohort-averaged partial autocorrelation function (PACF) of the standardized innovations, with approximate white-noise bounds. (C) Kolmogorov–Smirnov distances ($D$) from the probability-integral-transform analysis of the innovations across recordings, relative to recording-specific critical values. (D) Estimated time constants of the two drives ($\tau = 1/\kappa$) against the 95% intervals of their priors (grey).

### 17 Analysis pipeline and reproducibility

All results can be regenerated from the CSV files with the accompanying R project. The data folders (`data/prcp`, `data/fantasia`, `data/tmst`) are set in `config.R` and the scripts in `analysis/` are run in the order of Table S 12 (or with `Rscript tools/run_all.R`). The estimation engine (`engine/model_engine.R`) is the reference implementation of the model; the functions are in `R/`, in layers (`core`, `model`, `signals`, `steps`, `report`), and the scripts in `analysis/` only call them, so that each script reads as a short recipe (load, analyze every recording, combine, save). `R/model/model_variants.R` assembles the specifications $M_0$–$M_3$ from the engine's likelihood and prior terms, and an optional compiled port of the filter (`engine/fast_filter.cpp`) accelerates the multi-start analyses (its equivalence with the engine's filter is verified above). Random seeds are fixed in `config.R` and derived per recording. The figures are drawn from the saved results by `analysis/13_figures.R`; their appearance and the recordings showcased in the example figures are settings of `config.R` (section 6) and are recorded in `results/figure_info.rds`. The file `pipeline.R` of the project declares, for each script, the results it reads and writes (`docs/DATAFLOW.md` is generated from it), `Rscript tools/run_all.R --status` says which steps are up to date, and `results/provenance.txt` and `results/MANIFEST.csv` record the code version, the data and the checksum of every output of a run.

Table S 12: Analysis scripts and their outputs

| Script (analysis/) | Output (results/) | Manuscript content |
|---|---|---|
| 01_fit_all.R | fit_*.rds, df_population_summary.rds | Fit and innovation diagnostics; time constants (Figure S10) |
| 02_simulation_recovery.R | sim_recovery.rds | Recovery under known truth (Figure S1) |
| 03_identifiability.R | identifiability.rds | Results 1–2 (Figure 1, Table 3) |
| 04_sensitivity.R | sensitivity.rds | Sensitivity, prior contrast (Figure S2) |
| 05_prcp.R | prcp.rds | Result 3: every tilt and stand-up of each recording (protocol annotations data/prcp_events.csv when present), pressure response, lagged correlation, dose-response, responses by type of maneuver (Figure 2; Figures S6–S8) |
| 06_fantasia.R | fantasia.rds | Result 4: group contrasts, stability, respiration and baroreflex (Figure 3; Figure S9) |
| 07_recovery_benchmark.R | recovery_choice.rds | Recovery definition (Figure S3) |
| 08_event_detection_validation.R | event_detection.rds | Event detection (Figure S5) |
| 09_exercise.R | exercise.rds | Result 5 (Figure 4; Figure S6) |
| 10_hrv_comparators.R | hrv.rds | HRV comparators |
| 11_supplement_checks.R | supp_checks.rds | Bound, calibration, equivalence checks (Figure S4) |
| 12_preprocessing_checks.R | prep_checks.rds | Data-preparation checks (earlier vs current format, cleaning gate) |
| 13_figures.R | manuscript/figures/*.png | All figures |
